# The *wtf4* meiotic driver utilizes controlled protein aggregation to generate selective cell death

**DOI:** 10.1101/2020.02.05.935874

**Authors:** Nicole L. Nuckolls, Anthony C. Mok, Jeffrey J. Lange, Kexi Yi, Tejbir S. Kandola, Andrew M. Hunn, Scott McCroskey, Julia L. Snyder, María Angélica Bravo Núñez, Melainia L. McClain, Sean A. McKinney, Christopher Wood, Randal Halfmann, Sarah E. Zanders

**Affiliations:** Stowers Institute for Medical Research, Kansas City, MO 64110, USA; University of Missouri Kansas City, Kansas City, MO 64110, USA; Department of Molecular and Integrative Physiology, University of Kansas Medical Center, Kansas City, KS 66160, USA

## Abstract

Meiotic drivers are parasitic loci that force their own transmission into greater than half of the offspring of a heterozygote. Many drivers have been identified, but their molecular mechanisms are largely unknown. The *wtf4* gene is a meiotic driver in *Schizosaccharomyces pombe* that uses a poison-antidote mechanism. Here, we show that the Wtf4 proteins can function outside of gametogenesis and in a distantly related species, *Saccharomyces cerevisiae*. The Wtf4^poison^ protein forms dispersed, toxic aggregates. The similar Wtf4^antidote^ protein also forms aggregates but is sequestered within or near vacuoles and is mostly benign. The Wtf4^antidote^ can co-assemble with the Wtf4^poison^ and promote its trafficking to vacuoles. We show that neutralization of the Wtf4^poison^ requires both co-assembly with the Wtf4^antidote^ and aggregate sequestration, as mutations that disrupt either of these processes results in cell death. This work reveals that *wtf* parasites can exploit protein aggregate management pathways to selectively destroy gametes.

## Introduction

Meiotic drivers are selfish DNA sequences that break the traditional rules of sexual reproduction. Whereas most alleles have a 50% chance of being transmitted into a given offspring, meiotic drivers can manipulate gametogenesis to bias their own transmission into most or even all of an individual’s offspring (Burt and Trivers, 2006; Lindholm et al., 2016). This makes meiotic drive a powerful evolutionary force (Sandler et al., 1957). Meiotic drivers are widespread in eukaryotes and the evolutionary pressures they exert are thought to shape major facets of gametogenesis including recombination landscapes and chromosome structure (Crow, 1991; Dyer et al., 2007; Larracuente and Presgraves, 2012; Schimenti, 2000; Pardo-Manuel de Villena and Sapienza, 2001; Hammer et al., 1989; C. Grey et al., 2018).

Harnessing and wielding the evolutionary power of meiotic drive has the potential to greatly benefit humanity. Engineered drive systems, known as ‘gene drives,’ are being developed to spread genetic traits in populations (Lindholm et al., 2016; Burt, 2014; Gantz et al., 2015; Esvelt et al., 2014; Burt and Crisanti, 2018). For example, gene drives could be used to spread disease resistance alleles in crops. Alternatively, gene drives can be used to suppress human disease vectors, such as mosquitoes, or to limit their ability to transmit diseases (Lindholm et al., 2016; Burt, 2014; Gantz et al., 2015; Esvelt et al., 2014, reviewed in Burt and Crisanti, 2018). While there are many challenges involved in designing effective gene drives, natural meiotic drivers could serve as useful models or components for these systems (Lindholm et al., 2016; Burt, 2014). However, the molecular mechanisms employed by most meiotic drivers are unknown.

The recently characterized *wtf* gene family of *Schizosaccharomyces pombe* includes several meiotic drivers (Nuckolls et al., 2017; Hu et al., 2017; López Hernández and Zanders, 2018; Bravo Núñez et al., 2018; Bravo Núñez et al., 2020; Eickbush et al., 2019). The *wtf* coding sequences are small (∼1 kb) and encode autonomous drivers that specifically kill meiotic products (spores) that do not inherit the *wtf ^+^* allele from *wtf* ^+^/*wtf ^-^* heterozygotes. These drivers carry out targeted spore destruction using two proteins: a poison (Wtf ^poison^) to which all spores are exposed, and an antidote (Wtf ^antidote^) which rescues only the spores that inherit the *wtf ^+^* allele (Figure 1A and 1B). The two proteins of a given driver are encoded on largely overlapping coding sequences, but the antidote contains ∼45 additional N-terminal amino acids (Figure 1A). The small size and autonomy of the *wtf* drivers make them promising candidates for use in gene drive systems. It is important, however, to first understand more about the molecular mechanisms of the *wtf* proteins and whether they are likely to be functional in other species.

**Figure 1.**
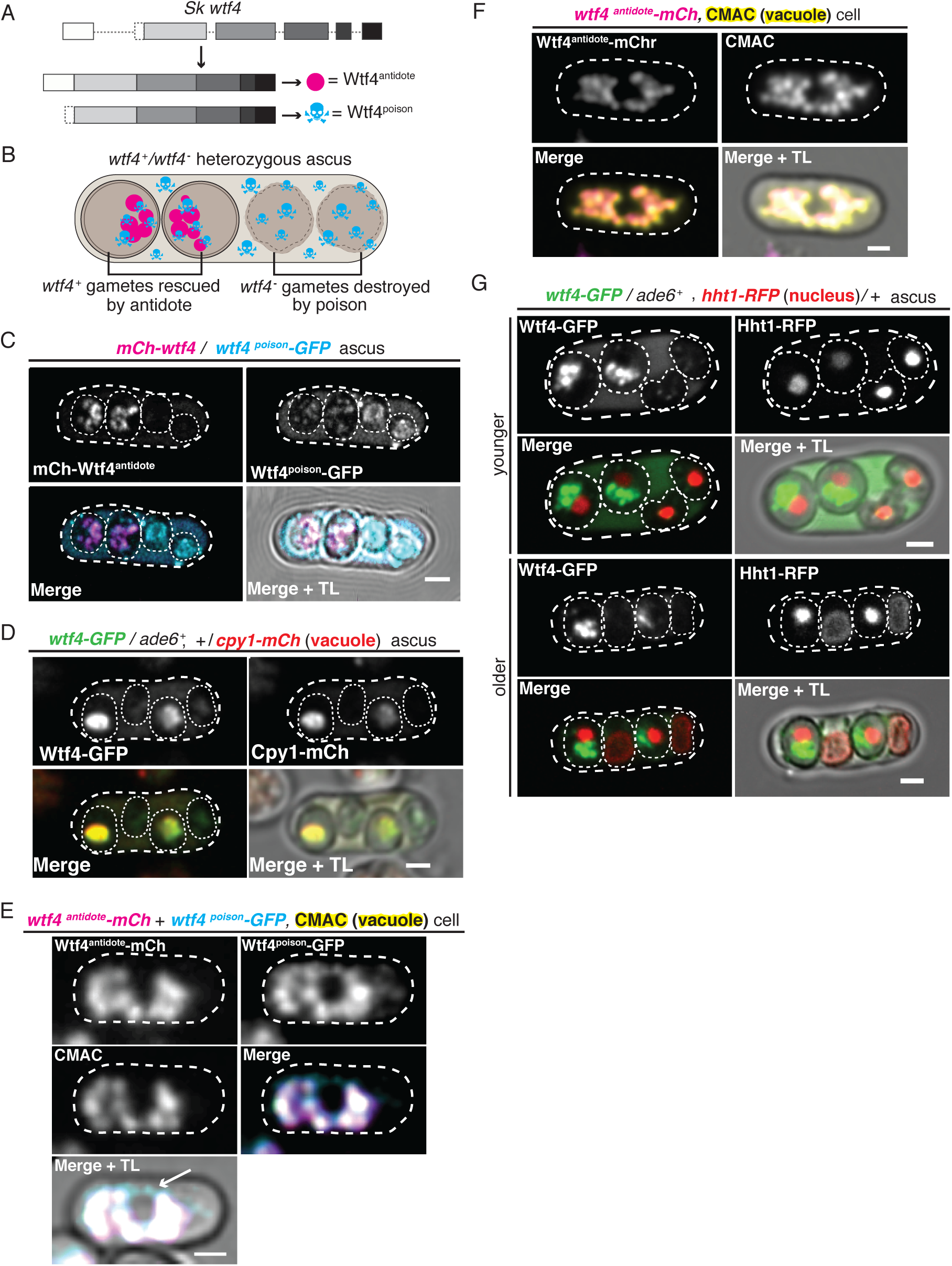
Wtf4^poison^ and Wtf4^antidote^ protein localization in both *S. pombe* meiosis and vegetative growth. (A) The *wtf4* gene utilizes alternate transcriptional start sites to encode Wtf4^antidote^ and Wtf4^poison^. (B) Model of a tetrad generated from a *wtf4*^+^/*wtf4*^-^ diploid. *wtf4*^+^ spores are rescued by the spore-enriched antidote (magenta circles), while the poison (cyan skulls) spreads throughout the ascus. (C) An ascus generated by an *mCherry-wtf4*/*wtf4^poison^-GFP* diploid showing the localization of mCherry-Wtf4^antidote^ (magenta in merged images) and Wtf4^poison^-GFP (cyan in merged images) (Nuckolls et al., 2017). (D) An ascus generated from a *wtf4-GFP*/*ade6*^+^, +/ *cpy1-mCherry* diploid showing localization of Wtf4-GFP (green in merged images) and Cpy1-mCherry (red in merged images). (E) A vegetatively growing haploid cell expressing Wtf4^poison^-GFP (cyan in merged images) and Wtf4^antidote^-mCherry (magenta in merged images) using the β-estradiol inducible system. CMAC is a vacuole lumen stain (yellow in merged images). Both Wtf4 proteins colocalize with the vacuole, except for a circle of Wtf4^poison^-GFP that lacks Wtf4^antidote^-mCherry (arrow). Cells were imaged 4 hours after induction with 100 nM β-estradiol. (F) A vegetatively growing haploid cell expressing Wtf4^antidote^-mCherry (magenta in the merged images) using the β-estradiol system and stained with the CMAC vacuole stain (yellow in the merged images). Cells were induced in the same way as in E. (G) Asci generated from a *wtf4-GFP/ade6^+^*, *hht1-RFP/+* diploid. Hht1-RFP (red in merged images) is a histone marker. All scale bars represent 2 µm.

Here, we investigate the mechanisms of *wtf* drive using the *wtf4* allele as a model. We demonstrate that the Wtf4 proteins are functional outside of gametogenesis and in the budding yeast *Saccharomyces cerevisiae*, despite over 350 million years since the two yeasts shared a common ancestor (*Hoffman et al.,* 2015). We also show that the two Wtf4 proteins assemble into distinct aggregated forms. Wtf4^poison^ forms small, toxic aggregates that are dispersed throughout the cytoplasm. The Wtf4^antidote^ forms aggregates that are recruited to vacuole-associated inclusions and are largely non-toxic. When the two Wtf4 proteins are expressed together, the Wtf4^antidote^ sequesters Wtf4^poison^ into vacuole-associated aggregates. This work adds to our understanding of how *wtf* meiotic drivers work. In addition, the conserved function of Wtf4^poison^’s toxicity and the fact that the Wtf4^antidote^ exploits conserved aggregate management processes suggests that *wtf* genes represent good candidates for gene drive systems.

## Results

### Wtf4 proteins localize to the vacuole and endoplasmic reticulum within *S. pombe* spores

The *wtf4* meiotic driver used in this work is from *S. kambucha*, an isolate that is almost identical (99.5% DNA sequence identity) to the commonly studied lab isolate of *S. pombe* (Rhind et al., 2011; Singh and Klar, 2002). Our previous work demonstrated that the Wtf4^antidote^ localizes to a region within the spores that inherit the *wtf4* gene. The Wtf4^poison^ protein, however, is found in all four spores and throughout the sac (ascus) that holds them (Nuckolls et al., 2017). We explored the localization of these proteins in greater depth to gain insight into their mechanisms.

We used fluorescently tagged alleles of *wtf4* to visualize the proteins. The two Wtf4 proteins have different translational start sites and thus different N-termini (Figure 1A, Figure1-figure supplement 1A). We took advantage of this feature to visualize the proteins separately. For the Wtf4^antidote^, we used an allele with an mCherry tag immediately upstream of the first start codon. This *mCherry-wtf4* allele tags only the Wtf4^antidote^ (mCherry-Wtf4^antidote^) but still encodes an untagged Wtf4^poison^. We previously demonstrated that this allele is fully functional (Nuckolls et al., 2017). To visualize Wtf4^poison^, we used the *wtf4^poison^-GFP* allele. This separation-of-function allele encodes only a C-terminally tagged poison, but no Wtf4^antidote^ protein. We previously demonstrated that this tagged allele is functional but has a slightly weaker phenotype than an untagged *wtf4^poison^* separation-of-function allele (Nuckolls et al., 2017).

We integrated the tagged alleles at the *ade6* locus in separate haploid *S. pombe* strains. We then crossed those two haploid strains to create heterozygous *mCherry-wtf4*/*wtf4^poison^-GFP* diploids and induced these diploids to undergo meiosis. We imaged the asci using both standard and time-lapse fluorescence microscopy (Figure 1C, Figure 1-figure supplement 1B). We confirmed our previous observations that the mCherry-Wtf4^antidote^ was enriched in two spores, whereas Wtf4^poison^-GFP was found throughout the ascus and often formed irregularly sized puncta. In the spores that did not inherit the antidote, Wtf4^poison^-GFP appeared dispersed throughout the spores. In the spores that inherited and thus expressed *mCherry-wtf4*, however, the localization of Wtf4^poison^-GFP was more restricted. Specifically, we observed that the Wtf4^poison^-GFP largely colocalized with mCherry-Wtf4^antidote^ in a limited region of the spore (Figure 1C). The Wtf4 proteins also co-diffused throughout the spore, suggesting the two proteins are either physically interacting or are in the same compartment (Figure 1-figure supplement 1B). It also appeared that the level of Wtf4^poison^-GFP protein is reduced in spores containing the antidote. We did not distinguish if this was due to technical reasons (i.e. quenching of the GFP molecules) or biological reasons such as degradation of Wtf4^poison^-GFP in spores with mCherry-Wtf4^antidote^ and/or due to a higher expression of Wtf4^poison^-GFP in the spores that inherit it (non-antidote spores) (Figure 1C). We completed Pearson correlation analysis (Adler and Parmryd, 2010) of mCherry-Wtf4^antidote^ and Wtf4^poison^-GFP in the spores (where a result of >0 is positive correlation; 0, no correlation; <0 negative correlation) and obtained a coefficient of 0.61, indicating strong colocalization between the two Wtf4 proteins (Figure 1-figure supplement 1C).

The limited distribution of the Wtf4 poison and antidote proteins within *wtf4^+^* spores suggested they may be confined to a specific cellular compartment. To test this, we looked for colocalization of Wtf4 proteins with the vacuole, endoplasmic reticulum (ER) and nucleus (see below). For these experiments, we used the fully functional *wtf4-GFP* allele, which tags both the poison and antidote proteins (Nuckolls et al., 2017).

To assay the localization of the Wtf4 proteins relative to the vacuole, we imaged asci produced by diploids that were heterozygous for both *wtf4-GFP* and *cpy1-mCherry*. Cpy1-mCherry localizes to the lumen of the vacuole in vegetative cells (Sun et al., 2013), but has not, to our knowledge, been imaged in spores. We could observe mCherry in two of the four spores– presumably the two that inherited the *cpy1-mCherry* allele. This 2:2 spore localization pattern has been previously observed in budding yeast for vacuolar proteins and several other organelles (Neiman, 2011; Roeder and Shaw, 1996; Suda et al., 2007). We found that the Wtf4-GFP and Cpy1-mCherry proteins colocalized within the spores that inherited both tagged alleles, suggesting the Wtf4 proteins are within the vacuole (Pearson coefficient of 0.89, Figure 1D, Figure 1-figure supplement 2A-B).

**Figure 2.**
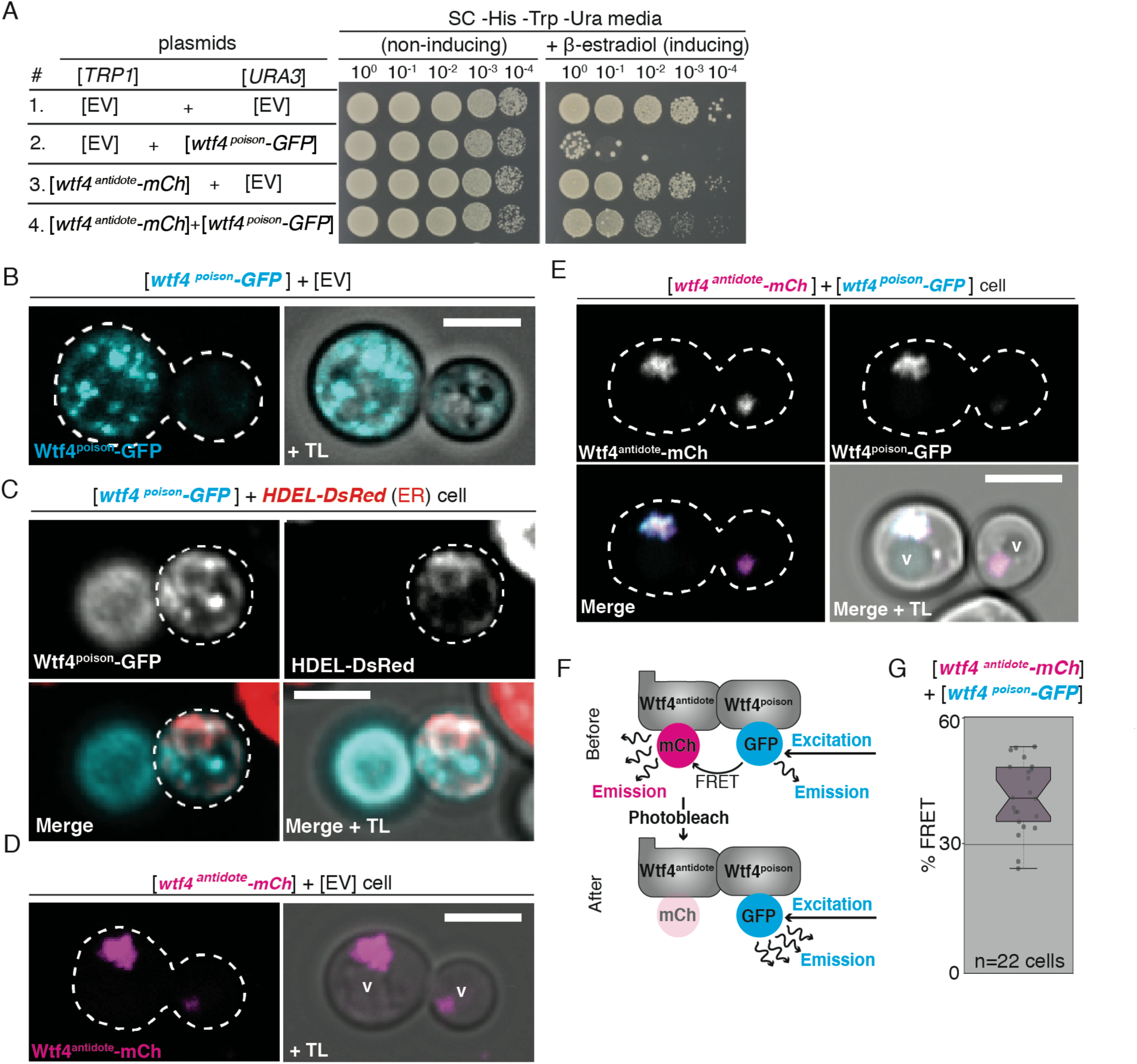
Wtf4^poison^ is toxic and Wtf4^antidote^ neutralizes that toxicity in *S. cerevisiae*. (A) Spot assay of serial dilutions on non-inducing (SC *-*His -Trp -Ura) and inducing (SC -His -Trp - Ura + 500 nM β-estradiol) media. Each strain contains [*TRP1*] and [*URA3*] ARS CEN plasmids that are either empty (EV) or carry the indicated β-estradiol inducible *wtf4* alleles. (B) A cell carrying an empty [*TRP1*] vector and a [*URA3*] vector with a β-estradiol inducible *wtf4^poison^-GFP* allele (cyan). (C) A cell with an integrated HDEL-dsRED allele that serves as an endoplasmic reticulum (ER) marker (red in merged images) and a [*URA3*] vector with a β-estradiol inducible *wtf4^poison^-GFP* allele (cyan in merged images). (D) A haploid cell carrying an empty [*URA3*] vector and a [*TRP1*] vector with a β-estradiol inducible *wtf4^antidote^-mCherry* allele (magenta). (E) A haploid cell carrying a [*URA3*] vector with a β-estradiol inducible *wtf4^poison^-GFP* allele (cyan in merged images) and a [*TRP1*] vector with a β-estradiol inducible *wtf4^antidote^-mCherry* allele (magenta in merged images). The vacuole is marked with ‘v’. (F) Cartoon of acceptor photobleaching <u>F</u>luorescence <u>R</u>esonance <u>E</u>nergy <u>T</u>ransfer (FRET). If the two proteins interact, Wtf4^poison^-GFP (the donor) transfers energy to Wtf4^antidote^-mCherry (the acceptor). After photobleaching of the acceptor, the donor emission will increase. (G) Quantification of FRET values measured in cells carrying β-estradiol inducible Wtf4^antidote^-mCherry and β-estradiol inducible Wtf4^poison^-GFP. The cells showed an average of 40% FRET. In all experiments, the cells were imaged approximately four hours after induction in 500 nM β-estradiol. All scale bars represent 4 µm.

Interestingly, we also saw colocalization of Wtf4-GFP proteins with an ER marker, pbip1-mCherry-AHDL (Zhang et al., 2012) in the spores generated by diploids heterozygous for alleles encoding those tagged proteins (Pearson coefficient of 0.74, Figure1-figure supplement 2C-D). However, we observed no colocalization of the Wtf4 proteins with the cortical ER. Due to the colocalization of Wtf4 with both organelles, we reasoned that the organelles themselves must colocalize in spores. This organelle colocalization could be due to nitrogen starvation, which is required to induce meiosis and promotes organelle autophagy in *S. pombe* (Zhao et al., 2016; Kohda et al., 2007).

### Wtf4^antidote^ localizes to the vacuole when expression is induced in vegetatively growing *S. pombe* cells

Because we could not distinguish the vacuole and ER within spores, we assayed the localization of the Wtf4 proteins in the absence of nitrogen starvation. To do this, we fluorescently-tagged the coding sequence of *wtf4^poison^* (*wtf4^poison^-GFP)* and *wtf4^antidote^* (*wtf4^antidote^-mCherry)* separation-of-function alleles under the control of β-estradiol–inducible promoters (Ohira et al., 2017). We then integrated the *wtf4^poison^-GFP* allele at the *ura4* locus and the *wtf4^antidote^-mCherry* allele at the *lys4* locus of the same haploid strain. Next, we observed the localization of the Wtf proteins relative to vacuole (visualized using the CellTracker Blue CMAC lumen stain) or the ER (using Sec63-YFP) following β-estradiol induction. Similar to our observations in spores, we saw that the Wtf4^poison^-GFP and Wtf4^antidote^-mCherry proteins largely colocalized, with a Pearson coefficient of 0.68 (Figure 1E, Figure1-figure supplement 3D-E). However, there were Wtf4^poison^-GFP puncta that lined the periphery of the cell and a circle in the middle of the cell, reminiscent of ER localization. These puncta were devoid of Wtf4^antidote^-mCherry (Figure 1E arrow, Figure1-figure supplement 3D). We also found that the Wtf4 proteins colocalized with the CMAC stain (Figure 1E), which suggests that the Wtf4 poison and antidote proteins are largely within the vacuole.

**Figure 3.**
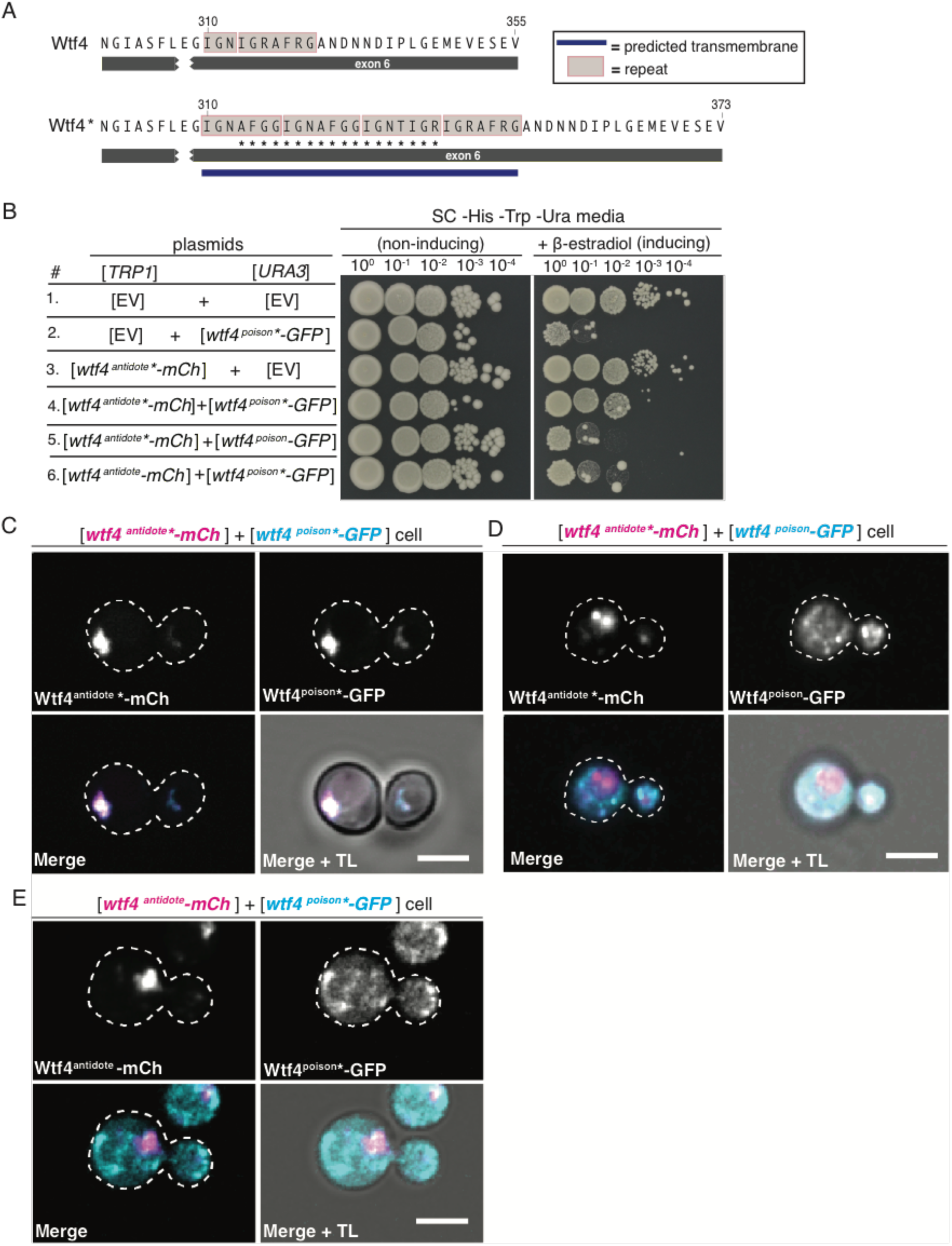
Homotypic interactions facilitate Wtf4^poison^-Wtf4^antidote^ co-assembly and Wtf4^antidote^ function. (A) Sequence encoding 18 amino acids (marked with an *) were added to the repeat sequences within exon 6 of the *wtf4* allele to create *wtf4*\* alleles. The addition of the amino acids introduces another predicted transmembrane domain (depicted by blue line; TMHMM model, Krogh et al., 2001). (B) Spot assay of serial dilutions on non-inducing (SC -His - Trp -Ura) and inducing (SC -His -Trp -Ura + 500 nM β-estradiol) media. Each strain contains [*TRP1*] and [*URA3*] ARS CEN plasmids that are either empty (EV) or carry the indicated β-estradiol inducible *wtf4* alleles. (C) Representative image of a haploid cell carrying a [*TRP1*] vector with a β-estradiol inducible *wtf4^antidote^*-mCherry* allele (magenta in merged images) and a [*URA3*] vector with a β-estradiol inducible *wtf4^poison^*-GFP* (cyan in merged images). (D) Representative image of a haploid cell carrying a [*TRP1*] vector with a β-estradiol inducible *wtf4^antidote^*-mCherry* allele (magenta in merged images) and a [*URA3*] vector with a β-estradiol inducible *wtf4^poison^-GFP* allele (cyan in merged images). (E) Representative image of a haploid cell carrying a [*TRP1*] vector with a β-estradiol inducible *wtf4^antidote^-mCherry* allele (magenta in merged images) and a [*URA3*] vector with a β-estradiol inducible *wtf4^poison^*-GFP* allele (cyan in merged images). In all experiments, the cells were imaged ∼4 hours after induction in 500 nM β-estradiol. All scale bars represent 4 µm.

We also attempted to assay the localization of the Wtf4 antidote and poison proteins individually to test if the localization of the Wtf4^poison^ was altered in the presence of the Wtf4^antidote^, as we observed in spores (Figure 1C). We found that the localization of the Wtf4^antidote^-mCherry to the vacuole was similar in the absence of the Wtf4^poison^ (Figure 1F, Figure1-figure supplement 3B), with a Pearson coefficient of 0.69 (Figure1-figure supplement 3C). This is analogous to previous observations of the localization of the slightly different (82.2% amino acid identity) Wtf4^antidote^ protein found in the *S. pombe* lab strain (Matsuyama et al., 2010). We failed, however, to generate cells carrying the *wtf4^poison^-GFP* allele without the *wtf4^antidote^-mCherry* allele by transformation or by crossing the strain carrying both *wtf4^poison^-GFP* and *wtf4^antidote^-mCherry* to a wild-type strain (Figure1-figure supplement 3A). This is likely due to leaky expression of the *wtf4^poison^-GFP* from the inducible promoter even without addition of β-estradiol. Overall, our results suggest that the Wtf4^poison^ protein is toxic in vegetative cells, but the antidote is still capable of neutralizing the poison, as we could obtain cells carrying both the Wtf4 poison and antidote proteins.

### *S. pombe* spores destroyed by *wtf4* display nuclear condensation followed by nuclear fragmentation

In the process of trying to understand the Wtf4 proteins’ localization patterns, we assayed the localization of the Wtf4 proteins relative to the nucleus. For this experiment, we imaged asci produced by *wtf4*-*GFP/ade6*^+^ heterozygotes also carrying a tagged histone allele, *hht1-RFP (Tomita and Cooper, 2017)*. Although we did not observe colocalization of Wtf4 proteins and the nucleus, we frequently (24/38 asci) observed that the nuclei in the *wtf4****^-^*** spores appeared more condensed (Figure 1G (younger ascus), see methods). Additionally, in 11 out of 38 asci, one or both of the nuclei in the *wtf4****^-^*** spores were disrupted and the nuclear contents were dispersed throughout the spores (Figure 1G (older ascus)). To address the timing of these nuclear phenotypes, we imaged diploids undergoing gametogenesis using time-lapse microscopy. We saw that all four nuclei tended to look similar shortly after the second meiotic division. As spores matured, however, we observed nuclear condensation sometimes followed by fragmentation in the spores that did not inherit *wtf4* (*i.e.* in spores lacking the enriched GFP expression and antidote function) (Figure 1-figure supplement 4A-B). This nuclear condensation and fragmentation are reminiscent of apoptotic cell death (Kerr et al., 1972; Carmona-Gutierrez et al., 2010).

### Wtf4 proteins function in the budding yeast, *Saccharomyces cerevisiae*

Our experiments in *S. pombe* suggest that the Wtf4 proteins can act when expressed outside of gametogenesis. Our inability to induce expression of the Wtf4^poison^ in the absence of the Wtf4^antidote^, however, limited our ability to explore their mechanisms of action in this system. We therefore tested if the Wtf4 proteins functioned in the budding yeast *Saccharomyces cerevisiae.* To do this, we cloned the coding sequences of *wtf4^poison^-GFP* and *wtf4^antidote^-mCherry* under the control of β-estradiol inducible promoters on separate plasmids (Ottoz et al., 2014). We then introduced these plasmids into *S. cerevisiae* individually and together. We found that cells carrying the *wtf4^poison^-GFP* plasmid were largely inviable when Wtf4^poison^-GFP expression was induced, indicating the poison is also toxic to *S. cerevisiae* (Figure 2A). However, cells expressing Wtf4^antidote^-mCherry had only a slight growth defect relative to control cells carrying empty plasmids (Figure 2A). Importantly, expression of the Wtf4^antidote^-mCherry plasmid largely ameliorated the toxicity of Wtf4^poison^-GFP (Figure 2A). Given that *S. pombe* and *S. cerevisiae* diverged >350 million years ago (Hoffman et al., 2015), our results suggest that the target(s) of Wtf4^poison^ toxicity are conserved and the Wtf4^antidote^ does not require cofactors that are specific to *S. pombe* or gametogenesis to neutralize Wtf4^poison^’s toxicity.

### Wtf4 poison and antidote proteins assemble into aggregates individually and together in budding yeast

We assayed the localization of the Wtf4 proteins in *S. cerevisiae* using the inducible *wtf4^poison^-GFP* and *wtf4^antidote^-mCherry* alleles described above. Similar to our observations in *S. pombe* meiosis, we saw that Wtf4^poison^-GFP localized as puncta of varying sizes throughout the cytoplasm (Figure 2B). We also observed some Wtf4^poison^-GFP localized to the ER (Figure 2C, Figure 2-figure supplement 1A). Analogous to our observation in *S. pombe* spores, we saw nuclear condensation in cells expressing Wtf4^poison^-GFP relative to wild-type cells (Figure 2-figure supplement 1B-D).

Wtf4^antidote^-mCherry, on the other hand, generally localized to one or two large amorphous regions adjacent to the vacuole (Figure 2D). When co-expressed, Wtf4^poison^-GFP and Wtf4^antidote^-mCherry co-localized to this region next to the vacuole (Figure 2E). In some cells, a faint circle of Wtf4^poison^-GFP could also be seen (likely ER localization); however, the majority colocalized with the antidote in the vacuole-associated region (Figure 2-figure supplement 1E, arrow). This localization was similar but not identical to our observations in *S. pombe* cells, where the Wtf4 proteins localize within, rather than adjacent to, the vacuole. To ensure the difference in localization (sequestration to a single puncta) and cell viability of the Wtf4^poison^-GFP protein observed in the cells co-expressing Wtf4^antidote^-mCherry was not due to the mCherry tag, we also confirmed these results with an untagged Wtf4^antidote^ (Figure 2-figure supplement 2A-B).

Because the Wtf4 proteins colocalize, we wondered if they physically interact. We tested this using acceptor photobleaching <u>F</u>luorescence <u>R</u>esonance <u>E</u>nergy <u>T</u>ransfer (FRET, Sekar et al., 2003, Figure 2F) in cells expressing both Wtf4^poison^-GFP and Wtf4^antidote^-mCherry proteins. This process involves bleaching the fluorescence of a tagged protein (the acceptor) and looking for a corresponding increase in fluorescence of another tagged protein (the donor). If an increase in fluorescence of the donor is observed, the proteins are said to be physically interacting, as they are in close enough proximity (less than 10 nanometers) to transfer energy to each other (Sekar et al., 2003). When we bleached Wtf4^antidote^-mCherry, we saw a corresponding increase in Wtf4^poison^-GFP emission, supporting the idea that the two proteins physically interact (Figure 2F-2G, Figure 2-figure supplement 1F).

The Wtf4 proteins localize as puncta of varying sizes, so we hypothesized that the proteins assemble into aggregates. To explore the nature of the Wtf4 protein assemblies, we utilized the recently developed Distributed Amphifluoric FRET (DAmFRET) assay (Khan et al., 2018). This approach looks for FRET between red and green fluorophores in a partially photoconverted population of mEos3.1-tagged proteins as a measure of the protein’s tendency to self-assemble (Figure 2-figure supplement 3A). We generated *wtf4^antidote^-mEos3.1* and *wtf4^poison^-mEos3.1* alleles, both under β-estradiol inducible promoters. Both tagged constructs encoded functional proteins in *S. cerevisiae,* but the mEos3.1-tagged Wtf4^poison^ allele was not as toxic as the GFP-tagged allele (Figure 2-figure supplement 3B). We then carried out DAmFRET analyses on cells expressing Wtf4^antidote^-mEos3.1 and on cells expressing Wtf4^poison^-mEos3.1. We observed high FRET signal between Wtf4^antidote^-mEos3.1 proteins and between Wtf4^poison^-mEos3.1 proteins. In fact, all cells expressing Wtf4^poison^-mEos3.1 or Wtf4^antidote^-mEos3.1 proteins exhibited FRET as compared to mEos3.1 negative control, regardless of the expression level of the proteins (Figure 2-figure supplement 3C). Collectively, these experiments confirm that the Wtf4 proteins self-assemble and assemble with each other.

### Homotypic interactions promote co-assembly of Wtf4 proteins and the neutralization of _Wtf4_poison

The Wtf4^poison^ and Wtf4^antidote^ proteins share the same 293 C-terminal amino acids (Figure 1A, Figure 1-figure supplement Figure 1A). All of the known active Wtf^antidote^ proteins are highly similar to the Wtf ^poison^ they neutralize (Bravo Núñez et al., 2020). In addition, mutations that disrupt the similarity between a given Wtf^antidote^ and Wtf^poison^ can eliminate the ability of the Wtf ^antidote^ to neutralize the Wtf ^poison^ (Hu et al., 2017; Bravo Núñez et al., 2018). Here, we tested the mechanism underlying that requirement using Wtf4 proteins. Given that each Wtf4 protein self-assembles, we hypothesized that homotypic interactions between Wtf4^poison^ and Wtf4^antidote^ mediated their co-assembly and neutralization of the poison. To test this idea, we mutated sequences at the C-termini of the inducible *wtf4^poison^-GFP* and *wtf4^antidote^-mCherry* alleles in the *S. cerevisiae* plasmids described above. Specifically, we targeted our mutagenesis to a seven amino acid repeat sequence (IGNAFRG) that is found in many members of the *wtf* gene family (Eickbush et al., 2019). We previously showed that a mismatched number of these repeats between a Wtf poison and antidote proteins is enough to disrupt their specificity (Bravo Núñez et al., 2018). The wild-type *S. kambucha wtf4* allele contains ∼1.5 repeat units (Figure 3A). To make the mutants, we inserted 18 additional codons into the repeat region of *wtf4* to make a total of four repeats. We denote these repeat insertion mutants with an * (Figure 3A).

As expected, the Wtf4^poison^*-GFP protein is functional (i.e. toxic) in *S. cerevisiae* and localizes similarly to the tagged wild-type Wtf4^poison^-GFP (Figure 3B, Figure 3-figure supplement 1A). Wtf4^poison^*-GFP is neutralized by the matching Wtf4^antidote^*-mCherry protein, and the two mutant proteins colocalized in vacuole-associated assemblies, just like the tagged wild-type proteins in *S. cerevisiae* (Figure 3B-C). Wtf4^antidote^*-mCherry protein on its own also resembled the wild-type Wtf4^antidote^-mCherry allele (Figure 3-figure supplement 1B). Wtf4^antidote^*-mCherry could not, however, suppress the toxicity of the wild-type Wtf4^poison^-GFP (Figure 3B). Similarly, the wild-type Wtf4^antidote^-mCherry could not neutralize Wtf4^poison^*-GFP (Figure 3B). The poison and antidote proteins did not colocalize in cells with incompatible poison and antidote proteins, and instead the poison proteins formed distributed aggregates, similar to cells expressing no antidote (Figure 3D, 3E).

### Electron microscopy reveals an association between Wtf4 aggregates and vesicles in *S. cerevisiae*

We next used transmission electron microscopy (TEM) to analyze the environment of Wtf proteins within the vacuole-associated aggregates. Similar to our observations made using fluorescence microscopy, we found using immuno-gold labeling that Wtf4-GFP largely clustered near the vacuole in cells also expressing untagged Wtf4^antidote^ (Figure 4A). These images also revealed that the Wtf4 protein aggregates appeared within a cluster of lightly staining organelles resembling lipid droplets (Figure 4A, Figure 4-figure supplement 1A, 1C). Very few immunogold particles were found in the cells carrying only empty vectors, suggesting minimal background and high specificity of the GFP antibody used for the immuno-labeling (Figure 4-figure supplement 1B).

**Figure 4.**
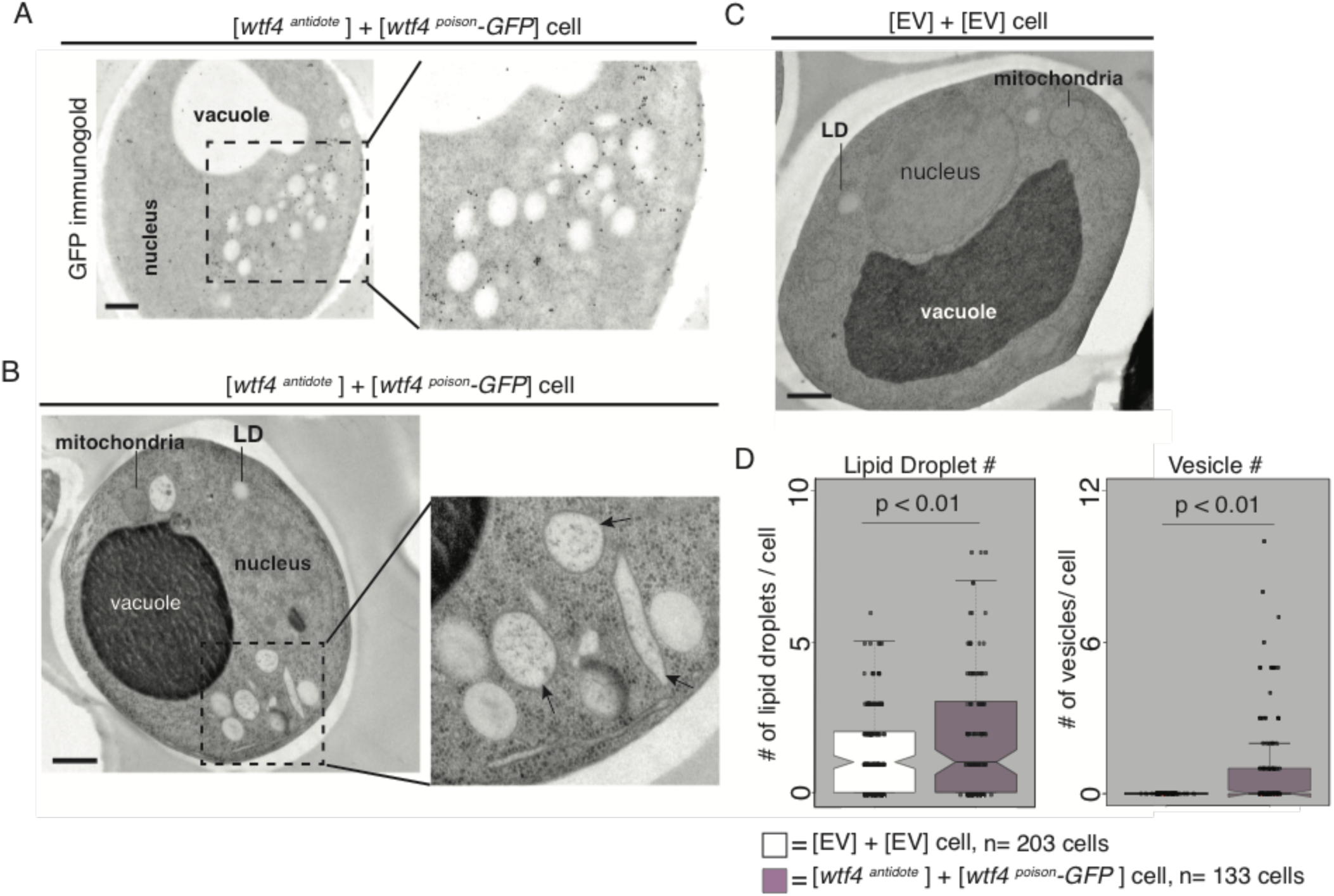
Cells expressing Wtf4^poison^ and Wtf4^antidote^ or Wtf4^antidote^ have increased vesicles that cluster within the Wtf4 aggregate. (A) Representative tomograph of Immuno-gold <u>T</u>ransmission <u>E</u>lectron <u>M</u>icroscopy (TEM) of a haploid cell carrying a [*TRP1*] vector with a β-estradiol inducible *wtf4^antidote^* allele and a [*URA3*] vector with a β-estradiol inducible *wtf4^poison^-GFP* allele. A monoclonal antibody against GFP was used. Immunogold particles (black dots) are enriched in a cluster near light staining organelles. (B) Representative TEM tomograph of a cell carrying a [*TRP1*] vector with a β-estradiol inducible *wtf4^antidote^* allele and a [*URA3*] vector with a β-estradiol inducible *wtf4^poison^-GFP*. Arrows point to vesicle structures. (C) Representative TEM tomograph of a cell carrying an empty [*TRP1*] vector and an empty [*URA3*] vector. (D) Quantification of the number of lipid droplets (left) or vesicles (right) per cell of two samples: 1. Cells carrying empty [*TRP1*] and [*URA3*] vectors (EV, white, n=203 cells) and 2. Cells carrying a [*TRP1*] vector with a β-estradiol inducible *wtf4^antidote^* allele and a [*URA3*] vector with a β-estradiol inducible *wtf4^poison^-GFP* allele (purple, n=133 cells), (p<0.01, t-test). All samples were processed ∼4 hours after induction in 500 nM β-estradiol. All scale bars represent 0.5 µm. Lipid Droplet=LD.

To look at these Wtf4 aggregate-associated organelles at higher resolution, we used TEM with a sample preparation method that better maintains cellular morphology (see methods). We found that the organelles were in fact a mix of lipid droplets and large vesicles with bilayer membranes (Figure 4B arrows, Figure 4-figure supplement 2A-C). We quantified the number of lipid droplets and large vesicles in cells carrying empty vectors, cells carrying a vector with β-estradiol inducible Wtf4^antidote^, and cells carrying both β-estradiol inducible Wtf4^antidote^ and β-estradiol inducible Wtf4^poison^-GFP. We found that cells expressing Wtf4 proteins had significantly more lipid droplets and large vesicles (Figure 4D, Figure 4-figure supplement 2F). These results indicate that the large aggregates that form in cells expressing Wtf4^antidote^ are embedded in a cluster of large vesicles and lipid droplets. This phenotype is reminiscent of another aggregation prone α-synuclein protein, a protein associated with Parkinson’s disease in humans, that when expressed in yeast forms cytoplasmic accumulations in association with clusters of vesicles (Soper et al., 2008). The α-synuclein vesicles, however, appear smaller and more numerous than the Wtf4-associated vesicles. To test if the increase in vesicles and lipid droplets was a common feature of aggregation prone proteins, we expressed (using the β-estradiol system) a different vacuole-associate prion aggregate, Rnq1-mCardinal (Figure 4-figure supplement 2E).

We did not observe any association with vesicles or lipid droplets, suggesting that the increase in vesicles is due to the Wtf4 aggregates, not a consequence of the over-expression system or a general feature of aggregation prone proteins.

We also imaged cells expressing only Wtf4^antidote^ or Wtf4^poison^. The morphology of cells expressing only the Wtf4^antidote^ was indistinguishable from cells expressing both Wtf proteins (Figure 4-figure supplement 2D). It is difficult to interpret the observations from the (dying) cells expressing only Wtf4^poison^ because the majority of the cells did not maintain cellular integrity during sample preparation (Figure 4-figure supplement 3A). In the few cells we could image, we observed diverse morphologies. We generally did not observe clustering of large vesicles and lipid droplets, as we saw in cells expressing Wtf4^antidote^. Instead, organelle integrity often looked disrupted and many cells expressing Wtf4^poison^ appeared to have undergone extensive autophagy (Figure 4-figure supplement 3B, 3C, 3D).

### Wtf4 poison-antidote protein aggregates localize to the IPOD and PAS in budding yeast

Our results were reminiscent of other studies in which toxic aggregated proteins were neutralized via sequestration at cellular inclusions (Kaganovich et al., 2008; Liu et al., 2010; Taylor et al., 2003; Chen et al., 2011; Hill et al., 2017; Tyedmers et al., 2010; Kryndushkin et al., 2012; Bagola et al., 2008; Arrasate et al., 2004). In *S. cerevisiae*, stable, misfolded proteins are generally sequestered to the Insoluble <u>P</u>r<u>O</u>tein <u>D</u>eposit (IPOD), a compartment located near the vacuole and <u>P</u>re-<u>A</u>utophagosomal <u>S</u>ite (PAS) (Kaganovich et al., 2008; Tyedmers et al., 2010, Suzuki and Ohsumi, 2010; Rothe et al., 2018). This compartmentalization of damaged/misfolded proteins mitigates their toxic effects, as well as facilitating their disposal, some of which occurs via autophagy (Marshall et al., 2016).

Given that the Wtf4^antidote^ and Wtf^poison^+Wtf4^antidote^ aggregates localize adjacent to the vacuole, we hypothesized that they could be at the IPOD in *S. cerevisiae*. To test this idea, we looked for the localization of the Wtf4 proteins relative to Rnq1-mCardinal and GFP-Atg8. Rnq1 localizes to the IPOD and Atg8 is a component of the pre-autophagosomal structure that is adjacent to the IPOD (Kaganovich et al., 2008; Tyedmers et al., 2010, Rothe et al., 2018). Consistent with our hypothesis, we found that Wtf4^antidote^-mCherry either colocalized or was adjacent to Rnq1-mCardinal (Figure 5A, Figure 5-figure supplement 1A) and was adjacent to GFP-Atg8 (Figure 5-figure supplement 1C). Wtf4^poison^-GFP did not colocalize with Rnq1-mCardinal on its own, supporting the idea that Wtf4^antidote^ recruits the poison to the IPOD (Figure 5-figure supplement 1B).

**Figure 5.**
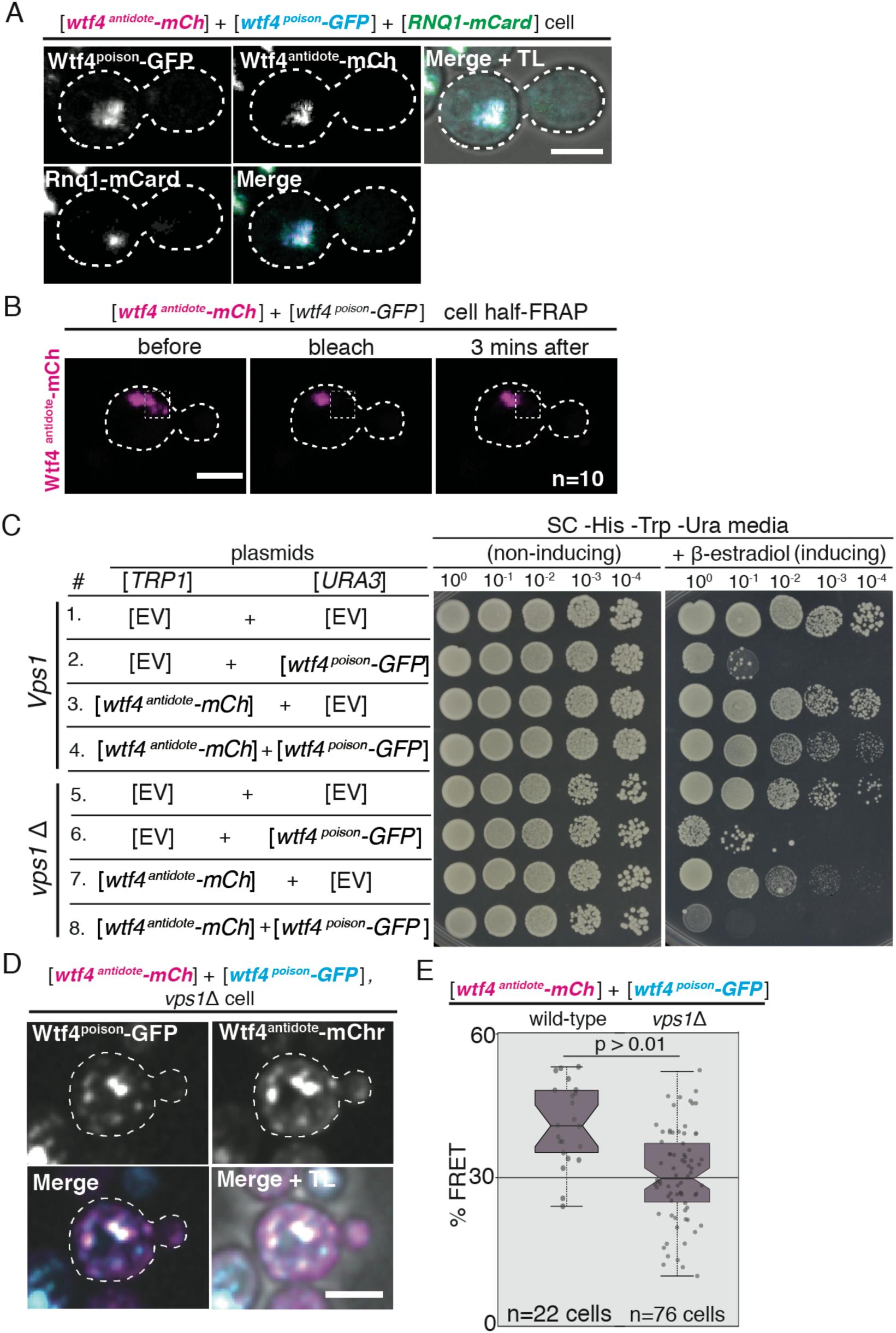
The Wtf4 proteins colocalize at the Insoluble <u>P</u>rotein <u>D</u>eposit (IPOD). (A) Representative image of a vegetatively growing haploid cell carrying a [*TRP1*] vector with a β-estradiol inducible *wtf4^antidote^-mCherry* (magenta in merged images) allele, a [*URA3*] vector with a β-estradiol inducible *wtf4^poison^-GFP* (cyan) allele, and a [*LEU2*] vector with a β-estradiol inducible *RNQ1-mCardinal* allele, acting as an IPOD marker (green in merged images). (B) half-<u>F</u>luorescence <u>R</u>ecovery <u>A</u>fter <u>P</u>hotobleaching (half-FRAP) of the Wtf4^antidote^-mCherry aggregate in cells carrying a [*TRP1*] vector with a β-estradiol inducible *wtf4^antidote^-mCherry* (magenta) allele and a [*URA3*] vector with a β-estradiol inducible *wtf4^poison^-GFP* allele. Cells were imaged for 3 minutes after bleaching and no recovery of mCherry fluorescence was seen. (C) Spot assay of serial dilutions on non-inducing inducing (SC -His -Trp -Ura) and inducing (SC -His -Trp -Ura + 500 nM β-estradiol) media of both wild-type (top, samples 1-4) and *vps1*Δ (bottom, samples 5-8) cells. Each strain contains [*TRP1*] and [*URA3*] ARS CEN plasmids that are either empty (EV) or carry the indicated β-estradiol inducible *wtf4* alleles. (D) Representative image of a vegetatively growing, haploid *vps1*Δ cell carrying a [*URA3*] vector with a β-estradiol inducible *wtf4^poison^-GFP* allele (cyan in merged images) and a [*TRP1*] vector with a β-estradiol inducible *wtf4^antidote^-mCherry* (magenta in merged images) allele. All fluorescence microscopy images acquired after ∼4 hours in 500 nM β-estradiol media. All scale bars represent 4 µm. (E) Quantification of FRET values of Wtf4^antidote^-mCherry and Wtf4^poison^-GFP measured in wild-type (same data as figure 2G) and *vps1*Δ cells carrying vectors with β-estradiol inducible *wtf4^antidote^-mCherry* allele and β-estradiol inducible *wtf4^poison^-GFP* allele (p >0.01, t-test).

Proteins in the IPOD tend to be insoluble (Kaganovich et al., 2008; Bagola et al., 2008). To test if the Wtf4^antidote^ shared this property in *S. cerevisiae*, we used <u>half</u> punctum-<u>F</u>luorescence <u>R</u>ecovery <u>A</u>fter <u>P</u>hotobleaching (half-FRAP) (Khan et al., 2018; Zhang et al., 2015). This analysis revealed that the Wtf4^antidote^-mCherry aggregate has very low internal mobility and is thus more solid-like than liquid-like (Figure 5B). We were curious if the Wtf4^antidote^ behaved similarly in its native context. To test this, we performed the half-FRAP assay on the Wtf4^antidote^-mCherry in *S. pombe* spores and again found very low protein mobility (Figure 5-figure supplement 1D-E).

### Genes involved in mitochondrial function, stress response pathways, and vesicle trafficking are necessary to neutralize Wtf4 protein toxicity in budding yeast

To better understand how toxic Wtf4 protein aggregates are neutralized, we screened for genes necessary for survival after induction of *wtf4^antidote^* and *wtf4^poison^*. Briefly, we screened the *S. cerevisiae MATa*, haploid deletion collection for mutants that failed to survive on galactose media when they carried plasmids encoding galactose-inducible *wtf4^antidote^-mCherry* and *wtf4^poison^-GFP* genes (Figure 5-figure supplement 2A-B). We found 106 mutants that could grow on galactose when carrying empty vector plasmids, but not when carrying both *wtf4* plasmids (Figure 5-source data 1).

Amongst our hits, the only significantly enriched (FDR p< 0.05) gene ontology groups were mitochondrial translation and organization (Figure 5-source data 1). We speculate this enrichment is due to two known roles of mitochondria in managing protein aggregates. The first is the <u>M</u>itochondria <u>A</u>s <u>G</u>uardian In <u>C</u>ytosol (MAGIC) mechanism by which mitochondria help degrade protein aggregates (Ruan et al., 2017). The second is that mitochondria mitigate the impact of toxic aggregates by promoting asymmetric aggregate segregation in mitosis (Zhou et al., 2014).

We also identified genes involved in <u>C</u>ell <u>W</u>all Integrity (CWI) pathways (*POP2*, *MPT5*, *SLT2* and *BCK1*) as necessary for survival after induction of Wtf4^antidote^ and Wtf4^poison^ (Jin et al., 2015; Li et al., 2016; Stewart et al.,2007). The CWI pathway is triggered by diverse stress stimuli (Fuchs and Mylonakis, 2009) and can promote stress-response gene expression and nuclear release of cyclin-C ((Ssn8), also a hit in our screen) (García et al., 2009). Release of cyclin-C into the cytoplasm promotes mitochondrial hyper-fission, stress response gene activation, and either apoptosis or repair of the stress-induced damage (Jin et al., 2015). Consistent with this, we observed separated, significantly smaller mitochondria in cells expressing the Wtf4 proteins (Figure 4-figure supplement 2C, 2G). Altogether, our screen hits suggest links between the CWI stress response pathways, mitochondrial fission, and Wtf4 antidote function.

Several other screen hits were genes with known roles in maintaining protein homeostasis and/or aggregate management. For example, we found that several genes involved in vesicle transport, endocytosis, and trafficking to the vacuole (e.g. *ATG11*, *SNF7*, and multiple *VPS* genes) are also required for survival when the Wtf4 proteins are expressed (Figure 5-source data 1). These hits suggest vacuolar trafficking pathways contribute to the neutralization of Wtf4 protein aggregates. This is consistent with our EM analyses showing that the Wtf4^antidote^ inclusion site is enriched with vesicles. Previous work demonstrated these pathways are also important for trafficking other proteins to the IPOD and for neutralizing the toxicity of the aggregation prone TDP-43 (Rothe et al., 2018; He et al., 2006; Liu et al., 2017).

Given our results, which suggest that Wtf4 protein localization is an important factor in mitigating toxicity, we next imaged the localization of Wtf4^poison^-GFP and Wtf4^antidote^-mCherry in all of the screen hits. We found that the localization of the Wtf4^poison^-GFP and Wtf4^antidote^-mCherry proteins was disrupted in all 106 hits relative to wild type (where the proteins coalesce to the IPOD). In 81 mutants, the Wtf4^poison^-GFP and Wtf4^antidote^-mCherry proteins localized as dispersed aggregates throughout the cell. These mutants included deletions of *YNL170W*, a reported dubious open reading frame, and *PHD1*, a transcriptional activator (Figure 5-figure supplement 2B-C). We noted that there were often cells with dispersed Wtf4^antidote^-mCherry aggregates or cells with dispersed Wtf4^poison^-GFP aggregates, but rarely cells with both. We speculate this is due to toxicity of distributed aggregates and cells expressing both aggregates at the same time being destroyed quickly. Another common feature we saw throughout the screen hits was Wtf4^antidote^-mCherry signal in the vacuole. We also observed this vacuolar localization in the C-terminal mutants depicted in Figure 3D-E, so this appears to be a common feature of the Wtf4^antidote^-mCherry protein in cells being destroyed by Wtf4^poison^. Five mutants appeared to have wild-type looking Wtf4^antidote^ (single inclusion outside the vacuole) but dispersed Wtf4^poison^, suggesting that the mutations may disrupt the interaction of the poison and antidote (Figure 5-figure supplement 2D-E). Twenty hits showed very little Wtf4 signal and soluble cytoplasmic localization (Figure 5-figure supplement 2F).

Because the Wtf4^antidote^ protein is quite similar to the Wtf4^poison^ and also assembles into aggregates, we were curious if the Wtf4^antidote^ alone was toxic in the absence of any of our screen hits. We therefore assayed the viability of the 106 deletion mutants when only Wtf4^antidote^ was expressed. We saw that in approximately half (44/106) of the deletion stains, Wtf4^antidote^ expression reduced viability (Figure 5-source data 1). These results are consistent with the idea that active aggregate management pathways are often required for cells to mitigate the toxicity of even the Wtf4^antidote^ protein in *S. cerevisiae*.

We also investigated one hit from our screen, *VPS1,* more thoroughly using our β-estradiol-inducible system (described above). *VPS1* is a dynamin-like GTPase that is necessary for trafficking of aggregates to the IPOD and/or other inclusions sites (Kumar et al., 2016; Kumar et al., 2017; Hill et al., 2016, Marshall et al., 2016). In the absence of *VPS1*, we found that the Wtf4^antidote^-mCherry and Wtf4^poison^-GFP proteins still physically interact (Figure 5D-E, Figure 5-figure supplement 3A). The Wtf4 protein aggregates did not, however, coalesce to form large inclusions (Figure 5D) and Wtf4^antidote^-mCherry failed to neutralize the toxicity of Wtf4^poison^-GFP (Figure 5C).

Together, these experiments indicate that the physical interaction between the Wtf4^poison^ and Wtf4^antidote^ proteins is insufficient to neutralize the toxicity of Wtf4^poison^ protein aggregates. Sequestering the aggregates to a vacuole-associated inclusion is also required. Interestingly, we also observed enhanced toxicity of the Wtf4^antidote^-mCherry protein in the absence of Vps1 and many of our other screen hits (Figure 5C, Figure 5-source data 1). These results suggest that the antidote aggregates are more detrimental to cells when they are distributed in the cytoplasm. Importantly, however, even in the *vps1*Δ mutant, expression of Wtf4^antidote^-mCherry is less toxic to cells than Wtf4^poison^-GFP. This, and the fact that not all of the 106 hits caused Wtf4^antidote^-mCherry to become toxic, suggests there are fundamental differences in the poison and antidote aggregates beyond their propensity to be trafficked to a vacuole-associated inclusion.

## Discussion

### Wtf4 proteins exploit conserved aspects of cell physiology to cause selective cell death

Here we explored how the Wtf4^poison^ protein kills cells and how the Wtf4^antidote^ protein neutralizes the toxicity of the Wtf4^poison^. We used a combination of genetics and cell biology to study these proteins in three contexts: 1) their endogenous context of *S. pombe* gametogenesis, 2) vegetatively growing *S. pombe* cells, and 3) vegetatively growing *S. cerevisiae* cells. In all three contexts, expression of Wtf4^poison^ alone kills cells and expression of the Wtf4^antidote^ rescues the toxicity. The simplest interpretation of these observations is that Wtf4^poison^ exploits or disrupts a conserved aspect of cellular physiology that is important during both vegetative growth and gametogenesis. Similarly, the Wtf4^antidote^ neutralizes the Wtf4^poison^ using conserved cofactors that can act in both vegetative growth and gametogenesis. This conservation suggests that *wtf*-derived gene drives could be a useful tool for genetically altering populations.

### Wtf4^poison^ proteins assemble into toxic aggregates

In *S. pombe* gametogenesis and in vegetative *S. cerevisiae* cells, we observed the Wtf4^poison^-GFP proteins assembled into small foci (aggregates) in the absence of Wtf4^antidote^. The aggregates were largely dispersed throughout the cytoplasm, with some ER localization in *S. cerevisiae*. The assembly of Wtf4 proteins is reminiscent of another meiotic drive element, Het-s, which employs prion-like amyloid polymerization to convert Het-S proteins to a lethal form (Dalstra et al.,2003; Riek and Saupe, 2016). We therefore evaluated whether Wtf4^poison^ proteins exhibit prion activity in *S. cerevisiae* using DAmFRET (Khan et al., 2018). We found that Wtf4^poison^-mEos proteins assembled with themselves even at very low expression levels (Figure2-figure supplement 3C). In fact, we were unable to detect cells that lacked self-assemblies, revealing that the toxic form of the protein is not appreciably supersaturated, as would be required for Wtf4^antidote^ to detoxify it through a simple prion-like mechanism. Nevertheless, the sequence-dependent self-assembly of Wtf4 remains consistent with amyloid polymerization. However, given its intimate association with vesicles, extensive testing would be required to further evaluate the structural basis of Wtf4 activity.

The significance of the Wtf4^poison^ aggregation is not clear. We speculate that the aggregation propensity is intimately tied to the toxicity of Wtf4^poison^. We propose that distributed Wtf4 aggregates interact broadly with other proteins and disrupt their folding or localization. Compounding effects of these hypothesized interactions could disrupt protein homeostasis or cellular integrity, leading to cell death. This death may occur via a programmed cell death pathway, as in both *S. pombe* gametogenesis and in vegetative *S. cerevisiae,* cells succumbing to the Wtf4^poison^ exhibit nuclear condensation (followed by nuclear fragmentation in *S. pombe*). The death may also be related to loss of cell wall integrity, as CWI pathways are necessary for cell survival upon expression of the Wtf4 proteins. Testing these ideas may be challenging, especially if understanding Wtf4^poison^ toxicity proves to be as elusive as understanding the intensely studied neurotoxic aggregating proteins TDP-43 and α-Synuclein (Johnson et. al, 2011; Cookson et. al, 2007).

### Wtf4^antidote^ promotes neutralization of Wtf4^poison^ via recruitment to vacuole-associated sites

Like Wtf4^poison^, the Wtf4^antidote^ also assembles into aggregates in both *S. pombe* and *S. cerevisiae* cells. Unlike the Wtf4^poison^, however, the Wtf4^antidote^ aggregates have little effect on the viability of wild-type vegetative cells or *S. pombe* spores. This is surprising given the similarity of the two proteins (the Wtf4^poison^ shares 292 of the Wtf4^antidote^’s 337 amino acids). Our data suggest that the localization of the aggregates and/or the exposed aggregate surface area could underlie their differences in toxicity. The Wtf4^poison^ aggregates (without Wtf4^antidote^) remain largely dispersed in the cytoplasm, whereas the Wtf4^antidote^ proteins are trafficked to a confined region near or within the vacuole. In *S. pombe* cells, the Wtf4^antidote^ aggregates enter the vacuole. In *S. cerevisiae* cells, the Wtf4^antidote^ accumulates outside the vacuole in the IPOD. However, it is unclear if some of Wtf4^antidote^ aggregates also enter the vacuole and are quickly degraded in *S. cerevisiae*.

When we disrupted the ability of *S. cerevisiae* cells to transport the Wtf4^antidote^ aggregates with the *vps1*Δ mutation, we found that the Wtf4^antidote^ aggregates were distributed and more toxic than in wild-type cells. This is consistent with the idea that a key feature of Wtf4 protein toxicity relies in the aggregates being widely dispersed in the cytoplasm. When Wtf4^poison^ and Wtf4^antidote^ are found together in wild-type cells, the proteins co-assemble into aggregates. The co-assembled aggregates then behave similarly to the Wtf4^antidote^ aggregates and are trafficked into the vacuole (in *S. pombe* cells) or to the IPOD adjacent to the vacuole (in *S. cerevisiae* cells) where they cause limited toxicity. Also, like the Wtf4^antidote^ aggregates, the toxicity of the Wtf4^poison^+Wtf4^antidote^ co-assembled aggregates is greatly enhanced if aggregate transport to the vacuole is disrupted by mutations (*e.g. vps1*Δ).

Together, our observations suggest a mechanistic model for *wtf4* function. In this model, *wtf4* exploits protein aggregation control pathways to induce selective cell death. The Wtf4^poison^ forms distributed toxic aggregates and the Wtf4^antidote^ co-assembles with the Wtf4^poison^ and neutralizes the aggregate’s toxicity via sequestration (Figure 6). This mechanism is unlike the mechanism of any other meiotic driver described to date (Grognet et al., 2014;Didion et al., 2015; Long et al., 2008; Dawe et al., 2018; Rhoades et al., 2019; Dalstra et al., 2005; Hammond et al., 2012; Vogan et al., 2019, Chen et al., 2008; Akera et al., 2017; Bauer et al., 2012; Pieper et al., 2018; Herrmann et al., 1999; Shen et al.,2017; Yu, et al., 2018; Bauer et al., 2007; Wu et al., 1988; Xie, et al., 2019; Kruger et al., 2019; Lin et al., 2018), but there are very few mechanistically characterized gamete-killing drive systems (reviewed in Bravo Núñez et al., 2018).

**Figure 6.**
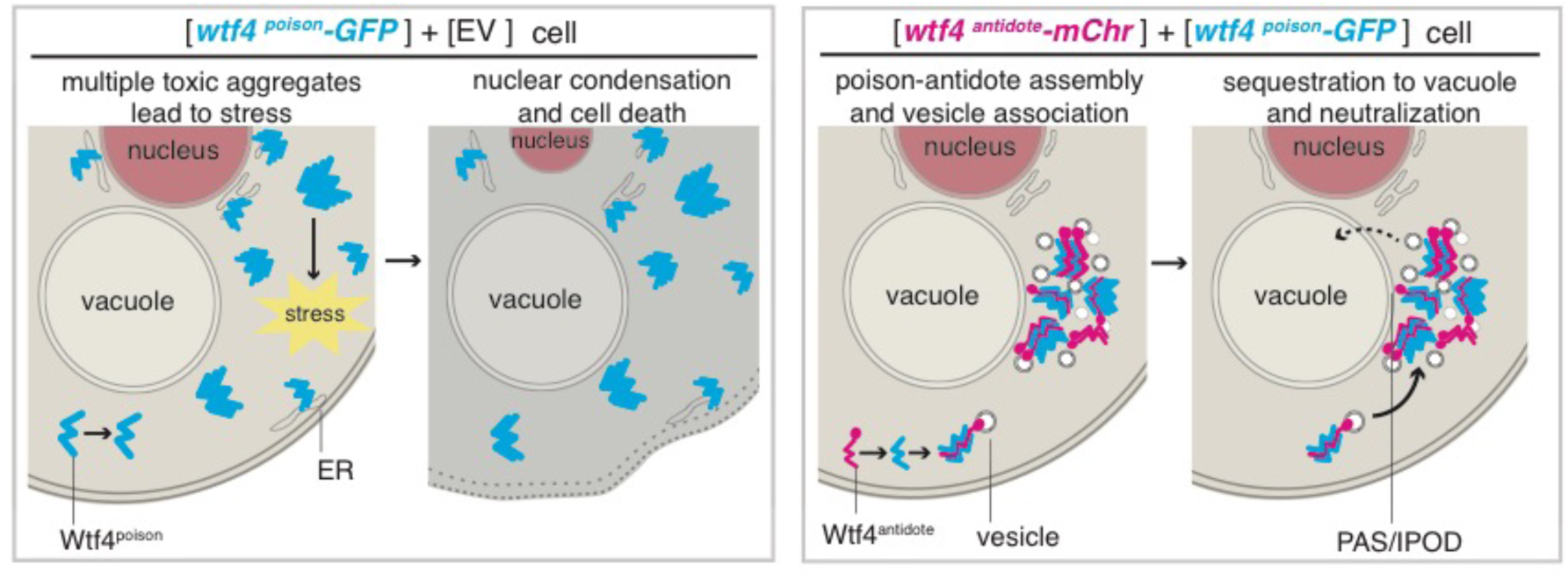
Model of Wtf4^poison^ and Wtf4^antidote^ mechanism in *S. cerevisiae*. Wtf4^poison^ assembles into toxic aggregates that spread throughout the cell, causing stress. This stress leads to nuclear condensation and cell death. Wtf4^antidote^ co-assembles with Wtf4^poison^, at least partially driven by shared sequences within the C-terminus. These assemblies are recruited to a vacuole-associated compartment that is associated with vesicles. This sequestration neutralizes the Wtf4^poison^ toxicity and rescues cell viability.

Continued investigation into how fission yeast and budding yeast species handle Wtf4 aggregates may elucidate important biological features of the two yeasts as well as a more detailed Wtf4 mechanism. Much less is known about aggregate management in *S. pombe*, but it offers an excellent model for understanding the process in symmetrically dividing cells (Coelho et al., 2014).

### A controlled protein aggregation model offers a solution to the *wtf* diversity paradox

This study focused on the *wtf4* meiotic driver. There is, however, an incredibly diverse array of *wtf* genes that cause meiotic drive. For example, the poison protein encoded by *wtf35* (from the FY29033 isolate) shares less than 23% amino acid identity with Wtf4^poison^ (Bravo Núñez et al., 2020). Despite that extreme divergence, both genes cause essentially the same phenotype: drive of the gene into > 90% of the progeny of a heterozygote (Bravo Núñez et al., 2020). The conserved protein aggregation model offers an explanation for how such a diverse array of proteins can cause the same phenotype. Under our model, the mechanism of the Wtf^poison^ proteins is dependent upon their aggregation propensity. Presumably, the evolution of a protein that must self-aggregate could be less constrained than the evolution of a protein that must maintain a specific enzymatic activity or interaction partner.

Exon 1 of the *wtf4* driver encodes the antidote-specific residues (45 amino acids) that facilitate the recruitment of Wtf4 aggregates to the vacuole in *S. pombe* (or the IPOD in *S. cerevisiae*). This antidote-specific function likely relies on unidentified interacting partners (perhaps amongst our screen hits). This specific functional requirement could explain the greater conservation of exon 1-encoded residues amongst *bona fide wtf* drivers (68-100% amino acid identity) compared to the conservation amongst the remaining exons (30-90% amino acid identity) (Bravo Núñez et al., 2020).

Importantly, our model also suggests that aggregate management may be a major feature of gametogenesis in *S. pombe*. The number of *wtf* genes varies between different isolates, but most have 30 or more *wtf* genes (Eickbush et al., 2019). Four of these genes are widely diverged from *wtf4* and are either not expressed in gametogenesis or their proteins exhibit distinct cellular localization from known Wtf^poison^ and Wtf^antidote^ proteins (*Bravo Núñez et al., 2020)*. The rest of the genes are similar to *wtf4* in expression and localization. It is not clear how many are expressed in a given cell, but they all appear to be transcribed at some level (Eickbush et al., 2019; Kuang et al., 2016). It will be interesting to explore the direct and indirect impacts of these Wtf proteins on *S. pombe* gametogenesis.

### Additional cellular factors are required to neutralize toxicity of Wtf4 proteins

The genetic screen presented in this work identified a number of factors required for cell viability in *S. cerevisiae* cells expressing Wtf4^poison^ and Wtf4^antidote^. Many of these genes informed our model for Wtf4 protein function and therefore fit nicely within our proposed model. For example, our screen implicated genes involved in the Cytoplasm-to-Vacuole Targeting (CVT) pathway as necessary for survival of the Wtf4 proteins. This pathway has been previously implicated in aggregate management (Kumar et al., 2016; Kumar et al., 2017).

Not all of our screen hits, however, are in genes or pathways with annotated roles that clearly fit our model. Some of the genes have no annotated functions. It is possible that at least some of these genes are not directly involved in aggregate management, but the mutants are especially sensitive to the stresses imposed by Wtf4 aggregates. It is also possible that some of the genes do have roles in mitigating the effects of toxic aggregates. Indeed, in deletions of some genes with unknown functions, we saw distributed Wtf4 aggregates, suggesting these unknown proteins could play a role in sequestration of aggregates. Interestingly, other hits are in well-studied genes, such as multiple acetyltransferases and various kinetochore proteins. Future analysis of these hits will be essential to refine or to potentially reject our current model.

### Insight into protein cellular response to aggregates via studying meiotic drive

Studying how parasites manipulate their hosts can uncover unexpected insights on the host’s biology. For example, studies of the mouse *t*-haplotype meiotic driver revealed that gene expression in spermatids can create sperm-autonomous phenotypes, even though spermatids are connected by intercellular bridges (Herrmann et al., 1999). Under our model, a fine line exists between protein aggregates that cells can manage (*i.e.* Wtf4^antidote^) and lethal aggregates that are not effectively managed (*i.e.* Wtf4^poison^). We propose that the *wtf4* meiotic driver has exploited this feature for its own selfish advantage. Future studies can now exploit the Wtf4 proteins to learn about protein aggregate toxicity and cellular aggregate management strategies.

## Materials and Methods

We confirmed all the vectors we generated (described below) via sequencing.

### Generation of tagged *wtf4* alleles for expression in *S. pombe* gametogenesis

#### Generation of a vector containing wtf4^antidote^-mCherry expressed from the endogenous promoter

We amplified the beginning of *wtf4* (including the endogenous promoter) from pSZB260 (described below) using oligos 688+719. The rest of the *wtf4^antidote^-mCherry* sequence was amplified (using oligos 605+1751) from pSZB708 (described below). The *ADH1* transcriptional terminator sequence was amplified from pSZB203 (Nuckolls et al., 2017) using oligos 1750+634. We then used overlap PCR (using oligos 688+634) to combine the three pieces. We then cloned the complete *wtf4^antidote^-mCherry* cassette into the SacI site of pSZB331 (Bravo Núñez et al., 2020) to generate pSZB891.

#### Generation of a vector containing wtf4^antidote^-GFP expressed from the endogenous promoter

We amplified the upstream sequence and the beginning of the *wtf4* allele from pSZB203 (Nuckolls et al., 2017) using oligos 620+736. We amplified the rest of the *wtf4-GFP* sequence (with an *ADH1* transcriptional terminator) from pSZB203 using oligos 735+634. Oligos 734 and 735 introduced mutations that interrupt the Wtf4^poison^ start site within intron 1. We then used overlap PCR with oligos 620+634 to unite the two pieces. We digested the complete *wtf4^antidote^*-GFP cassette with SacI site and cloned it into the SacI site of pSZB188 (Nuckolls et al., 2017) to generate pSZB260.

#### Generation of a vector containing the predicted wtf4^antidote^ coding sequence expressed from the endogenous promoter

We amplified the *wtf4* coding sequence in three pieces. We amplified the promoter with oligos 633+604 using SZY13 DNA as a template. We amplified the coding sequence from a gBlock DNA fragment (Integrated DNA Technologies, Inc., Coralville) using oligos 605+614. We amplified the sequence downstream of *wtf4* using oligos 613+635 and SZY13 genomic DNA as a template. We then stitched the three pieces together using overlap PCR with oligos 633+635. We then digested the product with SacI and ligated the cassette into SacI-digested pSZB188 (Nuckolls et al., 2017)^15^ to generate pSZB199. Intron 5 was predicted wrong, so there is a mutation at the C-terminus. Within this study, this plasmid was only used to build other plasmids, and when used in subsequent steps, we repaired the C-terminal mutation with the PCR oligos.

### *S. pombe* Z_3_EV β-Estradiol inducible system

Z_3_EV promoter system is a titratable inducible promoter system (Ohira et al., 2017). The system requires the Z_3_EV transcription factor and a Z_3_EV-responsive promoter (Z_3_EVpr). β-estradiol induces nuclear import of the Z_3_EV protein; therefore, genes placed immediately downstream of Z_3_EVpr in a strain expressing Z_3_EV become expressed upon β-estradiol addition to the media.

#### Background strain construction

To integrate the Z_3_EV transcription factor at the *leu1* locus of *S. pombe,* we digested plasmid pFS461 (Addgene #89064, Ohira et al., 2017) with XhoI and transformed it into the yeast strain SZY643 (selecting for Leu+) via standard lithium acetate protocol (Gietz, et al., 1995). This generated the yeast strain SZY2690, into which we transformed all of the proteins with Z_3_EV promoters (see below).

#### Generation of a strain that expresses wtf4^antidote^-mCherry under the control of a β-estradiol inducible promoter

We amplified the Z_3_EVpr from pFS478 (Addgene #89066, Ohira et al., 2017) using oligos 1734+1735. We then amplified the *wtf4^antidote^-mCherry* sequence (with an *ADH1* transcriptional terminator) from pSZB891 (described above) using oligos 1738+634. We used overlap PCR to add the Z_3_EV promoter piece to the *wtf4^antidote^-mCherry* piece using oligos 1738+634. We then digested this cassette with SacI and ligated it into the SacI site of pSZB322 (*Bravo Núñez et. al, 2018)*, a *lys4* integrating vector with a *hphMX6* cassette, to create pSZB892. We cut pSZB892 with KpnI and integrated into the *lys4* locus of SZY2690 to create SZY2740.

#### Generation of a strain that expresses wtf4^antidote^-mCherry and wtf4^poison^-GFP under the control of β-estradiol inducible promoters

To create an estradiol inducible Wtf4^poison^-GFP vector, we amplified the Z_3_EVpr on pSZB892 (see above) using oligos 1734+2068. We amplified the Wtf4^poison^-GFP (with an *ADH1* transcriptional terminator) from pSZB203 (Nuckolls et al., 2017) using oligos 2069+634. We then completed overlap PCR (using oligos 1734+634) on the two pieces. We then digested the completed *wtf4^poison^-GFP* cassette with SacI and ligated it into the SacI site of pSZB331 (see above) to create pSZB975. We cut pSZB975 with KpnI and integrated into the *ura4* locus of SZY2740 to generate SZY2888.

For the above transformations, we used high-efficiency, lithium acetate transformation protocol (Gietz, et al., 1995) to integrate the vectors, selecting first for drug resistance and then screening for the relevant auxotrophy.

#### Induction of Wtf proteins

For imaging cells (Figures 1E-F, Figure 1-Figure Supp 3), we created 5 ml saturated overnight cultures in rich YEL broth (0.5% yeast extract, 3% glucose, 250 mg/L of adenine, leucine, lysine, histidine, and uracil) supplemented with 100 ug/ml G418 and Hygromycin B (to select against pop-outs of the *lys4* and *ura4* integrating plasmids described above). The next day, we diluted 1 ml of each saturated culture into 4 mls of fresh YEL+G418+HYG media. We then added β-estradiol (from VWR, #AAAL03801-03) to a final concentration of 100 nM and shook the cultures at 32°C for four hours. We then used these induced cultures for imaging (see below for microscopy details).

### *S. cerevisiae* LexA-ER-AD β-estradiol inducible system

The LexA-ER-AD system (Ottoz et al. 2014) utilizes a heterologous transcription factor containing a LexA DNA-binding protein, the human estrogen receptor (ER) and an activation domain (AD). β-estradiol binds the ER and tightly regulates the activity of the LexA-ER-AD transcription factor. The LexA DNA-binding domain recognizes *lexA* boxes in the target promoter.

#### Background strain construction

To integrate the PACT1-LexA-ER-haB42 transcription factor into the *his3-11,15* locus of *S. cerevisiae,* we digested plasmid FRP718 (Addgene #58431, Ottoz et al. 2014) with NheI and transformed it into the yeast strain SLJ769 via standard lithium acetate protocol (Gietz, et al., 1995) selecting for His+ cells. This generated SZY1637, the β-estradiol inducible PACT1-LexA-ER-haB42 transcription factor strain, into which we transformed all the plasmids carrying genes under the control of LexA box containing promoters (LexApr) (see below). We generally used the ‘Sleazy’ transformation protocol (incubate 240 μl 50% PEG3500 + 36 μl 1M Lithium Acetate + 50 μl boiled salmon sperm + 1-5 μl DNA + a match head amount of yeast overnight at 30°C; modified from Elble, 1992) to introduce plasmids into *S. cerevisiae*. We selected transformants on Synthetic Complete (SC) media (6.7 g/L yeast nitrogen base without amino acids and with ammonium sulfate, 2% agar, 1X amino acid mix, 2% glucose) lacking appropriate amino acids for selection of the transformed plasmids.

#### Generation of a vector containing wtf4^poison^-GFP under the control of a β-estradiol inducible promoter

We amplified the LexApr from on FRP1642 (Addgene #58442, Ottoz et al. 2014) using oligos 1195+1240. We cloned the promoter into the KpnI/XhoI sites of pSZB464 (see below) to create pSZB585.

#### Generation of empty vectors containing only the β-estradiol inducible promoter

We amplified the LexApr from FRP1642 (Addgene #58442, Ottoz et al. 2014) using oligos 1195+1240. We cloned that promoter into KpnI+XhoI-digested pRS314 (ARS CEN *TRP1* vector, Sikorski et al., 1989) to generate pSZB668. We also cloned it into the KpnI+XhoI site of pRS316 (ARS CEN *URA3* vector, Sikorski et al., 1989) to generate pSZB670.

#### Generation of a vector containing wtf4^antidote^-mCherry under the control of a β-estradiol inducible promoter

We first amplified *wtf4^antidote^* (using oligos 1402+1401), *mCherry* (using oligos 1400+1399), and the *CYC1* transcriptional terminator (using oligos 1398+964) individually, using pSZB700 (see below) as the PCR template in all three reactions. We then used overlap PCR (using oligos 1402+964) to unite the three pieces into the *wtf4^antidote^-mCherry* cassette. We then digested the LexApr out of pSZB668 (with KpnI+XhoI) and ligated it, along with the *wtf4^antidote^-mCherry* cassette (digested with XhoI+BamHI), into KpnI+BamHI-digested pRS314 (Sikorski et al, 1989) to generate pSZB708.

#### Generation of a vector containing mCherry-wtf4^antidote^ under the control of a β-estradiol inducible promoter

We amplified *mCherry-wtf4^antidote^* coding sequence from pSZB248 (Nuckolls et al., 2017) using oligos 1066+604 and amplified the C-terminus of *wtf4^antidote^* plus the *CYC1* terminator from pSZB497 (see below) using oligos 1065+964. We then used overlap PCR (using oligos 1326+964) to join the two pieces. We digested the LexApr out of pSZB668 using KpnI and XhoI, and ligated that promoter, along with the XhoI-BamHI-digested *mCherry-wtf4^antidote^* cassette made by overlap PCR into KpnI-BamHI-digested pRS314 (Sikorski et al, 1989) to generate pSZB700.

#### Generation of a vector containing wtf4^antidote^ under the control of a β-estradiol inducible promoter

We digested the galactose inducible promoter out of pSZB497 (see above) w/ KpnI and XhoI. We amplified the LexApr (using oligos 1195+1240) from FRP1642 (Addgene #58442, Ottoz et al. 2014), digested with KpnI+XhoI, and ligated it into KpnI-XhoI-digested pSZB497 to generate pSZB589.

#### Induction of Wtf4 proteins with β-estradiol

For imaging, we grew 5 mL saturated overnight cultures in SC -His -Ura -Trp (without agar). The next day, we diluted 1 ml of the saturated culture into 4 mls of media of the same type. We then added β-estradiol to a final concentration of 500 nM and shook the cultures at 30°C to induce. Cells were induced for four hours and then imaged at one or multiple timepoints, depending on the experiment. For spot assays, we diluted saturated cultures to an OD of ∼1, then serial diluted (10^0^,10^-1^,10^-2^,10^-3^,10^-4^) in a 96-well plate.

We then spotted 10 ul of each dilution onto SC -His -Ura -Trp media and the same media with 500 nM β-estradiol. We grew the plates 2 to 3 days at 32°C and imaged on a SpImager (S&P Robotics).

### *S. cerevisiae* galactose inducible system

#### Generation of a vector containing wtf4^antidote^ under the control of a galactose inducible promoter

We amplified the beginning of the *wtf4^antidote^* coding sequence from pSZB388 (see below) using oligos 1065+678 and amplified the rest of *wtf4^antidote^* (and the *CYC1* transcriptional terminator) from pSZB392 (see below) using oligos 679+964. We then used overlap PCR with oligos 1065+964 to join the two pieces. We then digested the complete *wtf4^antidote^* cassette with XhoI and BamHI and ligated them into XhoI+BamHI-digested pDK20 (DasGupta et. al, 1998) to generate pSZB497.

#### Generation of a vector containing wtf4^poison^-GFP under the control of a galactose inducible promoter

We first amplified *wtf4^poison^* followed by a *CYC1* terminator from pSZB388 (see below) using oligos 963+964. We then digested the PCR product with XhoI and BamHI and ligated it into XhoI+BamHI-digested pDK20 (DasGupta et. al, 1998). This created pSZB392, a *URA3* integrating vector with *wtf4^poison^* under the control of a *GAL* promoter. We then amplified *wtf4^poison^* (including the *GAL* promoter) from pSZB392 with oligos 1045+606 and amplified GFP followed by an *ADH1* transcriptional terminator from pSZB203 (Nuckolls et al., 2017) using oligos 998+1040. We then stitched those two PCRs together using overlap PCR (amplifying with oligos 1040+1045). Finally, we digested the PCR product with KpnI and BamHI and cloned it into KpnI+BamHI-digested pRS316 (Sikorski et al, 1989) to generate pSZB464 and into KpnI+BamHI-digested pRS314 (Sikorski et al, 1989) to generate pSZB463.

#### Generation of a vector containing wtf4^antidote^-mCherry under the control of a galactose inducible promoter

We amplified the *wtf4^antidote^* sequence with the galactose inducible promoter from pSZB497 (see above) using oligos 1929+997 and the *wtf4^antidote^-mCherry* sequence (with a *CYC1* terminator) from pSZB708 (see above) using oligos 1072+964. We then used overlap PCR using oligos 1929+964 to combine the two pieces. We then digested the complete *wtf4^antidote^-mCherry* cassette with BamHI and ligated it into BamHI-digested PRS315 (Sikorski et al, 1989)^63^ to generate pSZB1005.

#### Generation of a vector containing wtf4^poison^ coding sequence

We amplified the *wtf4^poison^* coding sequence from pSZB199 (see above) using oligos 916+926. We then digested the PCR product with SfiI and cloned into SfiI-digested pBT3-STE (P03233DS from DUALsystems Biotech) to generate pSZB388. Within this study, this plasmid was only used to build other plasmids.

#### Induction of Wtf4 proteins with galactose

For imaging, we grew 5 ml saturated overnight cultures in SC media lacking appropriate amino acids for selection of the plasmids. The next day, we pelleted the cultures, resuspended in YP raffinose media, and grew overnight. The next day, we diluted 1 ml of the saturated raffinose culture into 4 mLs of SC galactose media lacking amino acids for selection of plasmids. We then added β-estradiol to a final concentration of 500 nM and shook the cultures at 30°C for four hours to create induced samples. For spot assays, we diluted saturated cultures to an OD of ∼1, then serial diluted (10^0^,10^-1^,10^-2^,10^-3^,10^-4^) in a 96-well plate. We then spotted 10 μl of each dilution onto both SC media (lacking amino acids appropriate for selection of the plasmids) and SC galactose media lacking the same amino acids. We grew the plates 2 to 3 days at 32°C and imaged them on a SpImager (S&P Robotics).

### Construction of the ER marker in *S. pombe*

To create the Sec63-YFP strain, we PCR amplified the C-terminus of *sec63* (using oligos 939+941) and the sequence downstream of *sec63* (using oligos 945+946) using SZY643 as a template. We also amplified a YFP-*HIS3* cassette from pYM41 (Janke et al., 2004) using oligos 944+943. We then used overlap PCR (using oligos 939+943) to unite those three PCR products. We then transformed this PCR product into GP1163 with standard lithium acetate protocol (Gietz, et al., 1995) (selecting for His+) to integrate the tagged *sec63-YFP* at its endogenous locus to generate SZY1277. We confirmed the strain via PCR using oligos 2037+2038.

### Generation the IPOD marker for expression in *S. cerevisiae*

#### Generation of a vector containing RNQ1-mCardinal under the control of a β-estradiol inducible promoter

We amplified the LexApr from pSZB708 using oligos 1835+1834, the *RNQ1* sequence from pDK412 (Kryndushkin et. al, 2012) using oligos 1833+1832 and the mCardinal-*CYC1* terminator from V08_mC (a gift from the Halfmann lab) using oligos 1831+964. We then used overlap PCR (using oligos 1835+964) to stitch the three pieces together. We then digested the cassette with BamHI and ligated it into BamHI-digested pRS315 (Sikorski et al, 1989) to generate pSZB942.

### Construction of *vps1*Δ *S. cerevisiae* strain

We PCR amplified the *vps1*Δ::*kanMX* locus out of strain YKR001C from the haploid yeast knockout *MATa* collection (Open Biosystems) (using oligos 1850+1851) and transformed the PCR product into SLJ769 using high efficiency lithium acetate protocol (selecting for G418 resistance) to create strain SZY2539. We used PCR (using oligos 1712+1713) and sequenced the locus to confirm the deletion. To add the PACT1-LexA-ER-haB42 transcription factor, we digested FRP718 (Addgene #58431, Ottoz et al. 2014) with NheI and integrated it into SZY2539 at the *his3-11,15* locus (using standard lithium acetate protocol (Gietz, et al., 1995) and selecting for His+) to create SZY2552.

### Generation of the *wtf4* exon 6 mutant alleles for expression in *S. pombe*

#### Generation of a vector containing wtf4*-GFP allele under the endogenous promoter

We introduced the mutation within exon 6 of *wtf4* using PCR (oligos 1280 and 1281 contain the desired mutation). We amplified the endogenous promoter and beginning of *wtf4* using oligos 688+1280 and pSZB203 as a template. We amplified the rest of *wtf4* and the downstream sequence using oligos 1281+686 and pSZB203 as a template. We used overlap PCR using oligos 688+686 to join the two pieces. We then digested the PCR product with SacI and cloned it into the SacI site of pSZB386 (Bravo Núñez et al., 2018) to generate pSZB647. Within this study, this plasmid was only used to build other plasmids.

### Generation of the wtf4 exon 6 mutant alleles for expression in S. cerevisiae

#### Generation of a vector containing wtf4^antidote^*-mCherry under the control of a β-estradiol inducible promoter

We amplified the beginning of *wtf4^antidote^* (using oligos 1402+1021) from pSZB700 (see above), the mutated section of *wtf4^antidote^\** (using oligos 1072+997) from pSZB647 (see above) and mCherry-*CYC1* terminator (using oligos 998+964) from pSZB708 (see above). We then stitched the three pieces together (using 1402+964) to generate the complete *wtf4^antidote^*-mCherry* cassette and digested it with XhoI and BamHI. We also digested the LexApr from pSZB708 using KpnI and XhoI. We then cloned those digested pieces into KpnI-BamHI digested pRS314 to generate pSZB774.

#### Generation of a vector containing wtf4^poison^*-GFP under the control of a β-estradiol inducible promoter

We amplified the beginning of *wtf4^poison^* (using oligos 1419+1021) from pSZB585 (see above). We amplified *wtf4^poison^\** (using oligos 1072+997) from pSZB647 (see above) and GFP+ the *ADH1* terminator (using oligos 998+1040) from pSZB585 (see above). We then used overlap PCR (using oligos 1419+1040) to join the three pieces. We digested the cassette with XhoI and BamHI and also digested the LexApr from pSZB668 (see above) using KpnI and XhoI. We ligated the digested *wtf4^poison^*-GFP* cassette and the LexApr into KpnI-BamHI-digested-pSZB668 to generate pSZB786.

### Generation of the mEos3.1 tagged *wtf4* alleles

#### Generation of vector containing wtf4^poison^-mEos3.1 under the control of a galactose inducible promoter

We used Golden Gate assembly (New England Biolabs) to insert the *wtf4^poison^* sequence that had been codon-optimized for *S. cerevisiae* (ordered from Addgene) into the BsaI-site of V08, a Gal-inducible vector with mEos3.1 and a rigid structure linker made up of a quadruple repeat of amino acids EAAAR [4x(EAAAR)] (Khan et al., 2018). This generated plasmid rhx1389.

#### Generation of vector containing wtf4^antidote^-mEos3.1 under the control of a galactose inducible promoter

We used Gibson assembly (New England Biolabs) using oligos rh1282+rh1283 to insert a sequence that encodes the 45 amino acids of the codon-optimized *wtf4* exon1 into the Aar1-digested rhx1389 (see above) to create pSZB1120.

#### Generation of a vector containing wtf4^poison^-mEos3.1 under the control of a β-estradiol inducible promoter

We amplified *wtf4^poison^-mEos3.1* with a *CYC1* terminator sequence (using oligos 1466+964) from rhx1389 (see above) and digested with BamHI. We then ligated the cassette into the BamHI site of pSZB668 to generate pSZB732.

#### Generation of a vector containing wtf4^antidote^-mEos3.1 under the control of a β-estradiol inducible promoter

We amplified the *wtf4^antidote^-mEos3.1* with a *CYC1* transcriptional terminator (using oligos 1465+964) from pSZB1120 (see above) and digested with BamHI. We then ligated the cassette into the BamHI site of pSZB670 (see above) to generate pSZB756.

#### Generation of a vector containing wtf4^antidote^ under the control of β-estradiol inducible promoters

We digested the LexApro-*wtf4^antidote^*-*CYC1* terminator construct out of pSZB589 (see above) and ligated it into pRS316 (Sikorski et al, 1989) to create pSZB782.

### DAmFRET

We induced samples of SZY2072, SZY2070, SZY2159, and SZY2059, with β-estradiol as described above. We then aliquoted these induced samples into a 96-well plate. We then partially photoconverted the mEos3.1 protein by exposing the plate, while shaking at 800 RCF, to 405 nm illumination for 25 mins using an OmniCure® S1000 fitted with a 320–500 nm (violet) filter and a beam collimator (Exfo), positioned 45 cm above the plate. This exposure yielded a total photo dose of 16.875 J/cm^2^. This photo dose reproducibly achieves the maximum fluorescence of the acceptor (red) form of mEos3.1 while minimizing photobleaching of the green form (Khan et al., 2018). For Figure 2-figure supplement 3C, we assayed the photoconverted samples on a Bio-Rad ZE5 cell analyzer with high throughput automation. We analyzed 20 µL of each sample to collect approximately 100,000 events per well. We excited the mEos3.1 donor (green form) with a 488 nm laser at 100 mW and collected with 525/35 nm and 593/52 nm bandpass filters, respectively. We excited the acceptor fluorochrome with a 561 nm laser at 50 mW and collected with a 589/15 nm bandpass filter. We performed manual compensation on-instrument at acquisition. We used DeNovo FCS Express for data analysis and visualization and calculated ratiometric FRET as FRET/acceptor signals.

### Fission Yeast microscopy

For imaging during gametogenesis (Figure 1C-D, 1G, Figure 1-figure supplement 1B, Figure 1-figure supplement 2A and 2C), we crossed the two haploid yeast strains to generate heterozygous diploids as previously described (Nuckolls et al., 2017). We placed the diploids on sporulation agar (SPA, 1% glucose, 7.3 mM KH_2_PO_4_, vitamins, agar) for 2-3 days. We then scraped the cells off of the SPA plates and onto slides for imaging.

For vegetatively growing samples (Figure 1E-F, Figure 1-figure supplement 3B and 3D), we induced gene expression with β-estradiol as described above. If we used vacuole staining, we took 1 mL of the induced culture, spun to pellet, and resuspended in 1 mL of 10 mM HEPES buffer, pH 7.4, containing 5% glucose with 100 μM CellTracker^TM^ Blue CMAC (Component B; Invitrogen C2110). We incubated these cells at room temperature for 30 minutes. We then washed with YEL media and imaged. For imaging, we used the LSM-780 (Zeiss) microscope with a 40x C-Apochromat water-immersion objective (NA 1.2) in photon-counting channel mode. For GFP, we used 488 nm excitation and collected through a 491–552 bandpass filter. For mCherry, we used 561 nm excitation and collected through a 572 longpass filter. For YFP, we used 514 nm excitation and collected through a 500-589 nm bandpass filter. For CMAC, we used 405 nm excitation and collected through a 411-509 nm bandpass filter. Brightness and contrast are not the same for all images. We analyzed at least 20 cells for each strain and chose a representative image. For experiments assaying meiosis/gametogenesis, we used at least two independent progenitor diploids; if cells were imaged during vegetative growth, we imaged at least three different starting cultures.

We carried out Pearson correlation analysis (Adler and Parmryd, 2010) as previously described (Slaughter et al., 2013). Briefly, we drew a segmented line (width of two pixels) throughout the spore, randomly covering as much of the spore as we could. We then used an in-house custom written plugin for ImageJ (https://imagej.nih.gov/ij/) to generate a two-color line profile. We calculated the Pearson correlation of the line profile with varying degrees of shifts in at least eight spores or six vegetatively growing cells per sample. We then combined and averaged the trajectories with standard error.

To quantify nuclear size, we calculated the full width at half maximum of the fluorescence intensity of RFP. We quantified 42 spores that inherited *wtf4-GFP* and 19 that did not, all from a *wtf4-GFP/ade6+* heterozygote after 2 days on SPA media. We excluded any nuclei that appeared to have already fragmented.

For the nuclear timelapse (Figure 1-figure supplement 4), we grew diploid cultures to saturation at 32°C overnight in YEL media. We then plated 100 µL of the cultures on a SPA plate, cut a circle punch of agar from the plate, and placed this punch upside down (cells facing down) in a 35 mm glass bottom poly-D-lysine coated dish (MatTek corporation). We placed grease around the edge of the MaTeK dish and a moist kim wipe inside to control for humidity. We then imaged the cells using the Nikon Ti Eclipse coupled to a Yokogawa CSU W1 Spinning Disk, using the 60x oil objective, acquiring images every ten minutes. Here, we excited RFP at 561nm and collected its emission through a 605-70 nm bandpass filter.

For the gametogenesis timelapse (Figure 1-figure supplement 1B), we grew diploid cultures to saturation at 32°C overnight in YEL media. The next day, we diluted 100 µL of the saturated diploid culture into 5 mLs of PM media (20 mLs of 50x EMM salts, 20 ml 0.4 M Na_2_HPO_4_, 25 mL 20% NH_4_Cl, 1 mL 1000x Vitamins, 100 µL 10,000x mineral stock solution, 3 g potassium hydrogen phthalate, 950 mL ddH_2_O, 25 mL of sterile 40% glucose after autoclaving, supplemented with 250 mg/L uracil). We grew the PM culture overnight at 32°C. The next day, we spun to pellet and resuspended the pellet in PM-N media (20 mLs of 50x EMM Salts, 20 mL 0.4 M Na_2_HPO_4_, 1 mL 1000x Vitamins, 100 µL 10,000x mineral stock solution, 25 mL of sterile 40% glucose after autoclaving, supplemented with 250 mg/L uracil, volume up to 1 L with ddH_2_O). We shook the PM-N cultures for 4 hours at 28°C. Then, we took 100 µL of the PM-N culture and mixed it with 100 µL of lectin (Sigma). We took 150 µl of this mixture and added it to a 35 mm glass bottom poly-D-lysine coated dish (MatTek corporation). We waited five minutes to allow the cells to adhere. We then added 3 mLs of fresh PM-N to the dish (protocol modified from Klutstein et al., 2015). We imaged using a Zeiss Observer.Z1 wide-field microscope with a 63x (1.2 NA) oil-immersion objective and collected the emission onto a Hamamatsu ORCA Flash 4.0 using μManager software. We acquired the mCherry with BP 530–585 nm excitation and LP 615 emission, using an FT 600 dichroic filter, acquiring images every 10 minutes.

### Budding yeast microscopy

For all budding yeast images except for the two experiments described below, we induced samples as described above and imaged on an LSM-780 (Zeiss) microscope, with a 40x C-Apochromat water-immersion objective (NA 1.2) in photon-counting channel mode. For GFP and mCherry, we used the same conditions as we did in *S. pombe.* For mCardinal, we used 633 nm excitation and collected through a 632-696 nm bandpass filter. Brightness and contrast are not the same for all images. We imaged at least 20 cells from at least three starting cultures and chose a representative image for each figure.

For imaging *vps1*Δ cells (Figure 5D) and the nuclear timelapse (Figure 2-figure supplement 2A-B), we placed samples in a Millipore Onix 2 Cellasic system to allow for a constant flow of media. We initiated flow of inducing media (SC with 500 nM b-estradiol for *vps1*Δ and SC galactose for the nuclear timelapse) and took images every 10 minutes. We used a Perkin Elmer Ultraview Vox spinning disc microscope with a Hamamatsu EMCCD (C9100-23B) with a 40x C-Apochromat water-immersion objective (NA 1.2). We collected GFP and mCherry with 488 and 651 nm excitation as above but collected GFP through a 525-50 nm bandpass filter and mCherry through a 615-70 nm bandpass filter. We had two independent starting cultures for the sample. We chose a representative cell and timepoint. Brightness and contrast are not the same for all images.

To quantify nuclear size (Figure 2-figure supplement 2C), we calculated the full width at half maximum of the fluorescence intensity of RFP per cell. We quantified at the beginning of the timelapse (early) and 14 hours into the timelapse (late). We quantified 72 Wtf4^poison^-GFP expressing cells and 65 wild-type cells at the early timepoint. We quantified 62 Wtf4^poison^-GFP expressing cells and 79 wild-type cells at the later timepoint.

### Acceptor Photobleaching FRET in *S. cerevisiae*

We carried out acceptor photobleaching FRET with β-estradiol induced (described above) SZY1954 (wild type) using a LSM-780 (Zeiss) microscope, with a 40x C-Apochromat water-immersion objective (NA 1.2) in photon-counting channel mode. For *vps1*Δ cells (SZY2570), we used a Perkin Elmer Ultraview Vox spinning disc microscope with a Hamamatsu EMCCD (C9100-23B) with 488 and 561 nm excitation. For both samples, we photobleached the acceptor (mCherry) with 561 nm excitation (for bleaching images, see Figure 2-figure supplement 2F for wildtype and Figure 5-figure supplement 3A for *vps1*Δ). We analyzed 22 wildtype and 76 *vps1*Δ cells.

### Fluorescence Recovery After Photobleaching half-FRAP of Wtf4 Aggregates

#### In S. cerevisiae

We induced SZY1954 with β-estradiol as described above and mounted into a lectin-coated 35 mm glass bottom poly-D-lysine coated dish (MatTek corporation) and imaged on a Perkin Elmer Ultraview Vox spinning disc with a Hamamatsu EMCCD (C9100-23B). We excited GFP with a 488 nm laser and collected its emission through a 100x alpha plan Apochromat objective (NA=1.4) and a 525-50nm bandpass filter. For each cell (n=10), we bleached half of the visible aggregate. We then acquired recovery images every second for three minutes total time.

#### In S. pombe

We placed SZY1142/SZY1049 heterozygous diploids on SPA plates for 2 days. We then scraped the sample off of the SPA plates into a 35 mm glass bottom poly-D-lysine coated dish (MatTek corporation) and carried out half-FRAP as above (n=10 spores), except that recovery images were then acquired for three minutes total time.

### Electron Microscopy

We made 50 mL saturated overnight cultures of SZY1821, SZY1952, SZY1954, and SZY2731 in SC media lacking histidine, tryptophan, and uracil (to select for retention of the plasmids). The next day, we diluted 10 mLs of the saturated cultures into 90 mLs of the same media with 500 nM β-estradiol. We shook these cultures for four hours at 30°C, reaching log phase. We then pelleted the yeast cells by filtering and carried out high pressure freezing with the Leica ICE system (Leica Biosystems). We further processed the frozen cell pellets by freeze substitution (FS) using acetone containing 0.2% uranyl acetate (UA) and 2% H_2_O was used as FS medium. The FS program was −90° to −80° over 70 hrs, −80° to −60° over 6 hrs, −60° for 5 h, −60° to −50° over 6 hrs, and −50° to −20°C over 4 hrs. After washing extensively with acetone, we then infiltrated, embedded and polymerized the samples into resin.

For Immuno-EM, we used HM-20 resin. We cut 60 nm sections with a Leica Ultra microtome (Leica UC-6) and picked up onto a carbon-coated 150 mesh nickel grid. The grids were labeled with anti-GFP primary antibody (a gift from M. Rout, Rockefeller University, New York, NY) and 12 nm colloidal gold goat anti-Rabbit secondary antibody (Jackson Immuno Research Laboratories, Inc). After immuno labeling, we post-stained the samples with 1% UA for 3 minutes. We acquired images using a FEI Tecnai Biotwin electron microscope. For non-immuno-EM, we used Epon resin to better maintain morphology, but the rest of the procedure was the same. We analyzed the tomographs of at least 10 cells per condition.

For quantification purposes, we also used completed array tomography. For array tomography, we cut 60 nm serial sections with a Leica Ultra microtome (UC-6) using an Ultra 35 Jumbo diamond knife (Diatome) and picked up on ITO coated coverslips using the ASH-100 Advanced Substrate Holder (RMC Boeckeler). We post-stained serial sections with Sato’s triple lead stain for two minutes, 4% UA in 70% methanol for two minutes, and Sato’s lead stain again for two minutes. The coverslips were coated with 5 nm of carbon and imaged in a Zeiss Merlin Gemini 2 SEM with 4QBSD detector at 10 kV and 700 pA using Atlas 5 Array Tomography software (Fibics). The obtained dataset was aligned with Midas of the IMOD software package (Kremer et al., 1996) and manually quantified. For better visualization, an image series of an individual yeast cell was cropped and further aligned with registration tools in Image J.

For model building, the segmentation was done based on intensity and known organelle structure with Microscopy Image Browser (Belevich et. al, 2016) and with IMOD. We used Amira (Thermo Fisher Scientific) software for model rendering and visualization.

Further quantification of mitochondrial volumes was performed on selected cells after training a Unet (Ronneberger et al., 2015). Hand annotation of training data was performed in Fiji. A suite of internally developed Fiji plugins, macros and CherryPy scripts called DeepFiji (see below) sent training data to a pair of in-house NVIDIA Tesla-equipped deep learning machines running Tensorflow. Representative cells were selected, and segmented images inferred using the same macros and deep learning machines before being aligned using a StackReg variant. Mitochondrial volumes were quantified in Fiji using the 3D Segmentation tools.

### DeepFiji training

DeepFiji is a suite of macros and plugins in Fiji, Python, and CherryPy (a Python web framework) that enable end users on any machine with a reasonable amount of RAM to request deep learning training and inference on a remote deep learning box as long as both machines have access to a shared file system.

First, a user selects example sub-images that span the realm of potential objects, background levels and signal levels. Manual annotations are made using Fiji’s Region of Interest (ROI) tools and manager, and individual ROI files are saved for each image (in our case the ROIs were each individual cell and the total of the mitochondria inside). A user chooses a small subset of the annotated image/ROI pairs to be used as a validation set, while the remainder becomes the training set. For each image, two binary channels are added from the associated ROIs: a mask channel and an outline channel. The mask channel has all pixels contained within an ROI painted true, while the outline channel only paints true the pixels that were in the ROI’s outline. As the training network expects standard image sizes and runs more efficiently with smaller images, the macro next makes image stacks for both annotated training and validation images that break the original images up into 512×512 sub-images with 50% overlap between sub-images. For the training set, it also applies a series of random rotations and translations to help the neural network generalize. Both the validation and new training images are saved to a shared file system and the deep learning boxes are notified to begin processing through a call to a webserver running on the box. The sub-image size, file location, and other parameters are configurable by the user at the time of running.

For our in-house system, the deep learning boxes are running ubuntu with NVIDIA Tesla boards and configured with Tensorflow 1.13.1 and CUDA 10.0. CherryPy is configured to listen for web calls on each and, once initiated from a user, begins processing files from the selected directory by calling trainer.py. The trainer first finds the standard deviation (STDEV) and mean (MEAN) of the non-zero pixel intensities and stores those values. The training, validation, and all future inference sets will be processed by (Intensity-MEAN)/STDEV+0.5 first to keep numbers roughly between 0 and 1. The model used is a modified Unet(ref): convolutional layers are alternated with max pool layers, doubling the channel depth at each layer while halving the resolution. The final layer is 32×32×512. At each convolutional layer the image is passed through a leaky rectified linear unit. Convolutional up-sampling brings the image back to its original resolution where it is passed through a tanh() function and the mean square error is calculated with respect to the ground truth image for back-propagation. Every 100th iteration of the training is applied to the validation set and the images are visualized using TensorBoard (which opens automatically on the user’s host computer). Training proceeds with the learning rate adjusting over time, until 2200 iterations have passed at which point most networks have either converged or never will.

Once training is completed, and a reasonable iteration point is found in TensorBoard, the user can run Inferer.ijm in Fiji on their host machine to apply their model to a new dataset. Inferer will similarly parse images into sub-images, and contact a deep learning box to initiate processing.

Once processing is complete the user can run a second script to blank out border regions and de-window their images. The output outline and mask channels are ranged from 0-1 and represent probabilities. Typically, thresholding pixel values above 0.5 in the mask channel will suffice for finding objects of interest. However, in cases with frequent object touching, one can subtract the outline probability from the mask.

If consistent mistakes are found in the inferred data, the user can annotate them properly using Fiji and use Retrain.ijm to retrain the network using the new data together with the old to generate a new training set. Retraining starts from the original model so that it does not have to relearn from scratch.

Plugins necessary to run in Fiji are available from the Stowers update site within Fiji. Macros, python code, and CherryPy configurations are available at https://github.com/cwood1967/DeepFiji.

### Genetic screen for suppressors of Wtf4^antidote^ function in *S. cerevisiae*

#### Screen design

Two separate plasmids carrying galactose inducible Wtf4 proteins [(Wtf4^poison^-GFP (pSZB464) and Wtf4^antidote^-mCherry (pSZB1005)] were transformed into the *MATa* haploid *S. cerevisiae* deletion collection (purchased from Open Biosystems) using lithium acetate protocol (Gietz, et al., 1995). The transformed collection was then spot inoculated on SC and SC galactose media lacking leucine and uracil. As a control, a strain expressing the galactose inducible poison and antidote in a wild-type background was used for comparison. We grew the plates for three days at 30°C, imaged them using the SpImager (S&P Robotics), and manually scored growth. For the antidote-only screen, we transformed the galactose driven Wtf4^antidote^-mCherry (pSZB1005) plasmid into the 106 hits from the first screen and scored them as mentioned above.

#### Confirmation of hits

The initial screen identified 250 strains that grew poorly on inductive media. To confirm that this poor growth was due to the Wtf4 proteins and not due to the background strain being sick or a poor grower on galactose media in general, we completed a follow-up screen. We transformed the 250 strains we identified as “poor growers” with empty [*URA3*] and [*LEU2*] vectors and assayed the strains as above to identify those that grew poorly on galactose media independent of *wtf4* gene expression. We found 106 strains that passed this secondary screen, which we then called hits. We imaged this 106 strains after a short galactose induction (∼4) to ensure we saw Wtf4 protein

#### Analysis

To look for enriched gene ontology terms in the hits from the screen, we used the PANTHER overrepresentation Test (Thomas et al., 2003; Thomas et al., 2006). The background list we used for the analysis was the list of *MATa* deletion collection strains that we successfully transformed with our plasmids of interest (n= 4793). We used Fisher’s Exact test and corrected with false discovery rate. We imaged the cells as described above for Gal-inductions, but we added 80 mg/L adenine to the inducing media to circumvent any potential autofluorescence introduced by the adenine auxotrophy.

## Acknowledgments

We thank Alejandro Rodríguez Gama, Julie Cooper, Li-Lin Du, Snezhana Oliferenko, Risa Mori, and Frank Shewmaker for strains or plasmids. We thank Kausik Si for valuable advice on the project, Sue Jaspersen for advice and reagents, and Xia Zhao for assistance in electron microscopy. Original data underlying this manuscript can be accessed from the Stowers Original Data Repository at http://www.stowers.org/research/publications/libpb. This work was performed to fulfill, in part, requirements for NLN’s thesis research in the Graduate School of the Stowers Institute for Medical Research. This work was supported by The Stowers Institute for Medical Research (SEZ, RH); the Searle Award (SEZ); National Institutes of Health (NIH) R00GM114436 and DP2GM132936 (SEZ); NIH Director’s Early Independence Award DP5-OD009152 (RH); National Cancer Institute of the NIH under award number F99CA234523 (MABN). Eunice Kennedy Shriver National Institute Of Child Health & Human Development of the NIH under Award Number F31HD097974 (NLN). The funders had no role in study design, data collection and analysis, or manuscript preparation. The content is solely the responsibility of the authors and does not necessarily represent the official views of the funders.

---

**Figure 1-figure supplement 1.**
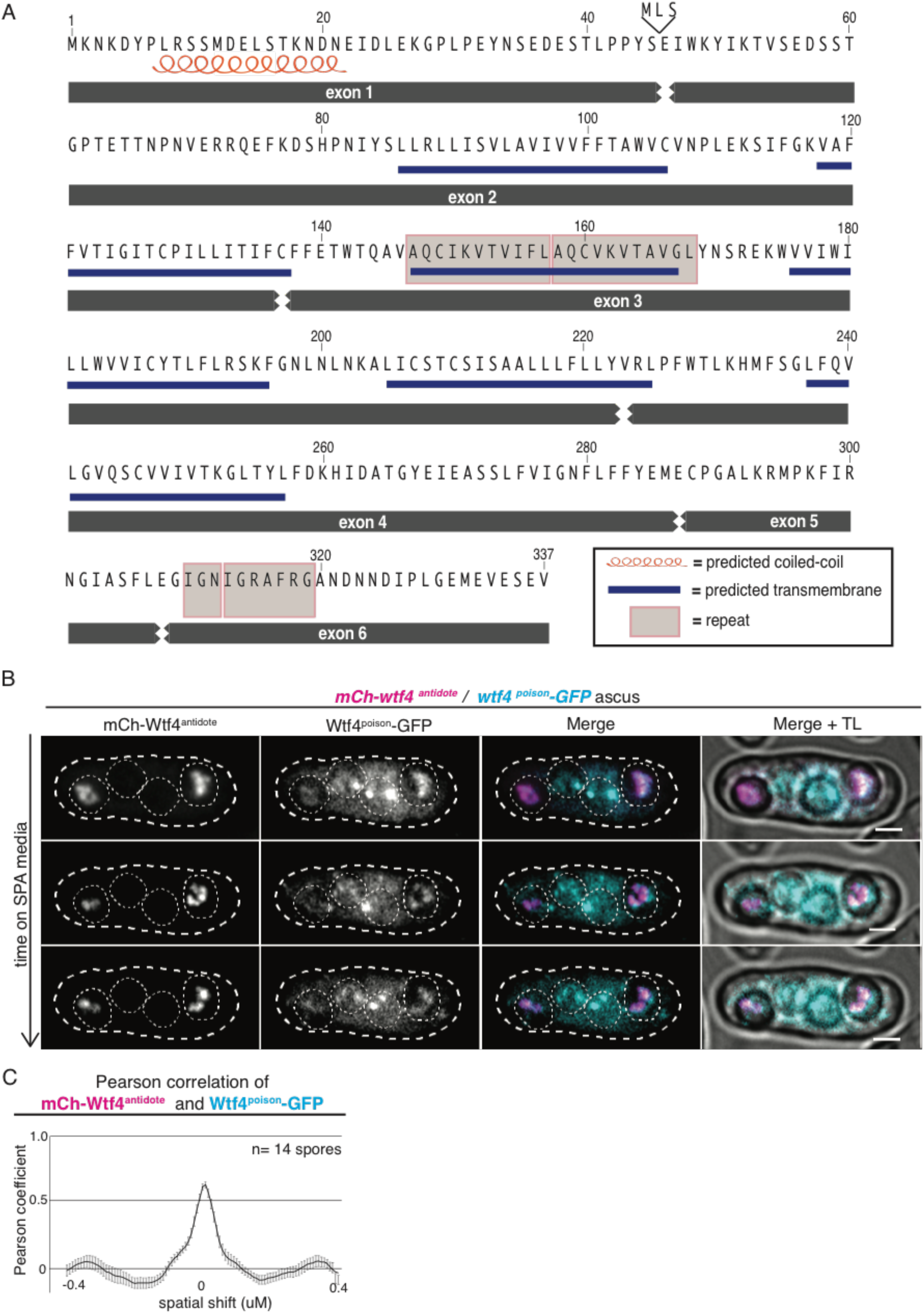
Wtf4^poison^ and Wtf4^antidote^ protein sequences and localization in *S. pombe* asci. (A) Wtf4 protein sequence. Wtf4^antidote^ is encoded by all 6 exons. Wtf4^poison^ is encoded by exons 2-6 and gains three amino acids (MLS) from intron 1. Exon 1 contains a predicted coiled-coil domain (depicted with an orange coil, Lupas, Dyke, and Stock, 1991). There are six predicted transmembrane domains (depicted by blue lines) (TMHMM model, Krogh et al., 2001) found throughout the amino acid sequences shared by both proteins. There are also two regions of repetitive sequences: one found in exon 3 and one found in exon 6 (Eickbush et al., 2019). These are depicted with gray boxes. (B) Time-lapse microscopy of an ascus generated by a *mCherry-wtf4/wtf4^poison^-GFP* diploid. mCherry-Wtf4^antidote^ is shown in magenta and Wtf4^poison^-GFP as cyan in merged images. All scale bars represent 2 µm. (C) Pearson correlation of mCherry-Wtf4^antidote^ and Wtf4^poison^-GFP from spores within asci generated from *mCherry-wtf4/wtf4^poison^-GFP* diploids is shown. Error bars represent standard error.

**Figure 1-figure supplement 2.**
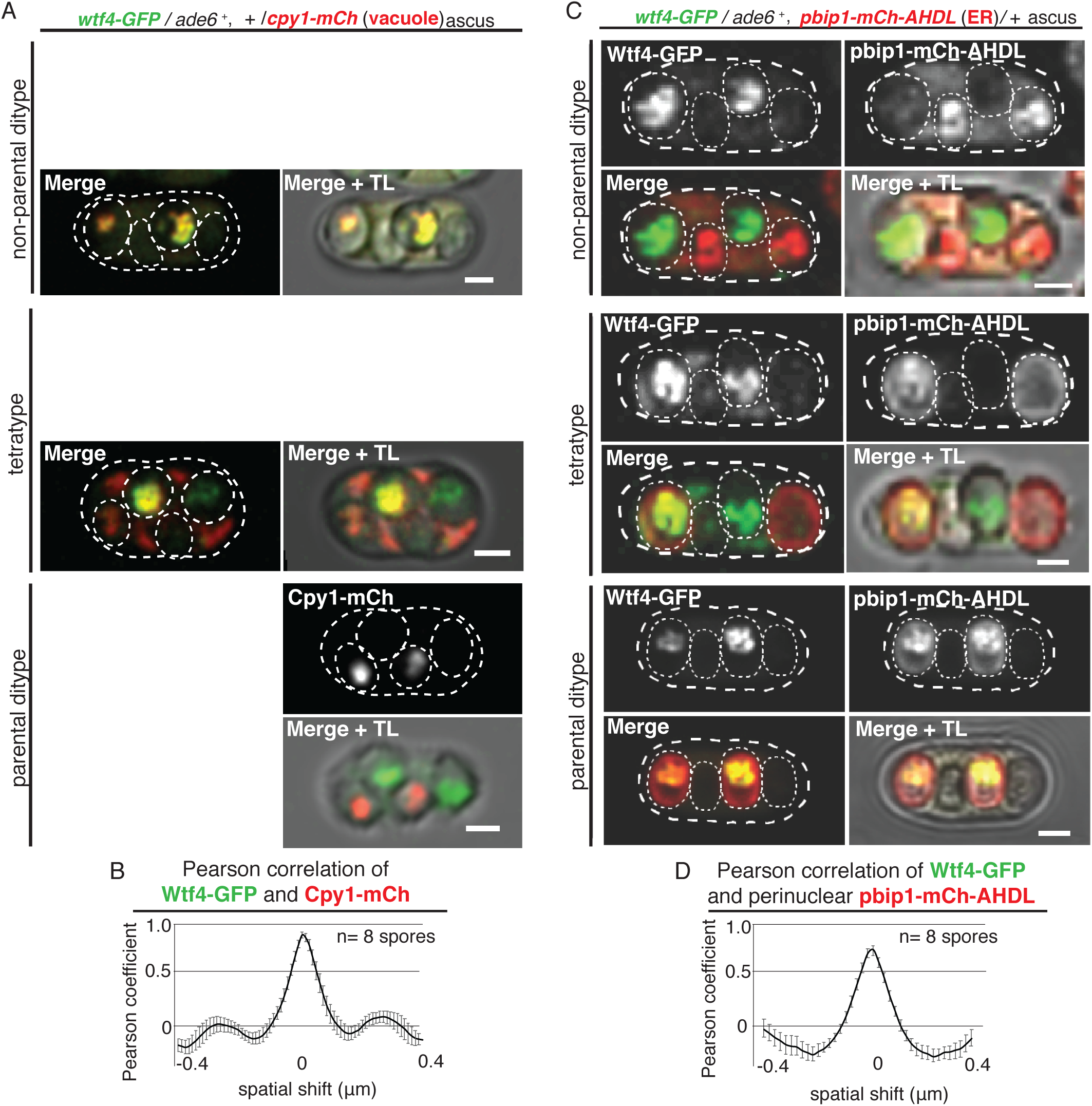
Wtf4^poison^ and Wtf4^antidote^ colocalize in the vacuole and endoplasmic reticulum (ER) in spores. (A) Non-parental ditype, tetratype, and parental ditype asci generated by *wtf4-GFP/ade6^+^, +/cpy1-mCherry* diploids. Cpy1-mCherry (red in merged images) is a vacuole luminal marker and Wtf4-GFP is shown in green in merged images. (B) Pearson correlation of Wtf4-GFP and Cpy1-mCherry in spores that inherited both alleles generated by *wtf4-GFP/ade6^+^, +/cpy1-mCherry* diploids shows a coefficient of approximately 0.89. (C) Non-parental ditype, tetratype, and parental ditype asci generated by *wtf4-GFP/ade6^+^*, *pbip1-mCherry-AHDL/+* diploids. pbip1-mCherry-AHDL (red in merged images) is an ER marker and Wtf4-GFP is shown in green in merged images. All scale bars represent 2 µm. (D) Pearson correlation of Wtf4-GFP and pbip1-mCherry-AHDL from spores that inherited both alleles generated by *wtf4-GFP/ade6^+^, pbip1-mCherry-AHDL/+* diploids shows a coefficient of approximately 0.74. Error bars represent standard error.

**Figure 1-figure supplement 3.**
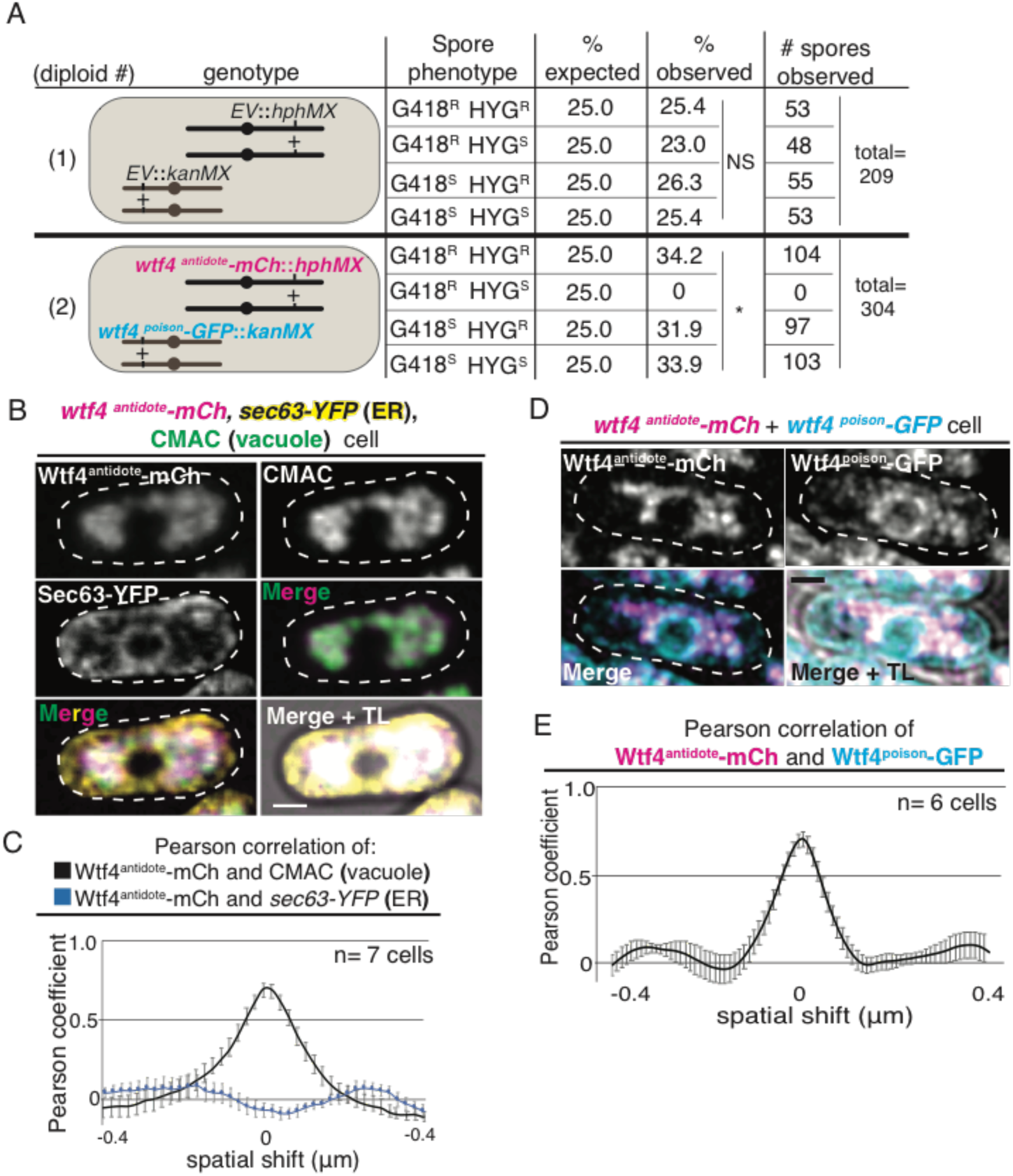
Wtf4^poison^ and Wtf4^antidote^ function outside of gametogenesis. (A) Results of allele transmission assays from two diploids. Diploid 1 has an empty vector (EV) with a hygromycin resistance gene (*hphMX*) integrated at *lys4* and another EV with a G418 resistance gene (*kanMX*) integrated at *ade6*. Diploid 2 has a vector with a *wtf4^antidote^-mCherry* allele (under the control of the β-estradiol promoter) and hygromycin resistance integrated at *lys4*. It also has a vector with a *wtf4^poison^-GFP* allele (under the control of the β-estradiol promoter) and G418 resistance integrated at *ade6*. The *ade6* and *lys4* loci are unlinked, and there are four possible spore genotypes in relation to drug resistance (R) or sensitivity (S). The diploids were sporulated on regular SPA media (without β-estradiol) and random spore analysis was completed (Smith, 2009). The expected percentage of each spore genotype and the observed percentages are shown, as well as the raw numbers (* = p < 0.01, g-test; NS= not significant). (B) A vegetatively growing haploid cell expressing a β-estradiol inducible *wtf4^antidote^-mCherry* allele (magenta in merged images). It also contains a tagged *sec63-YFP* allele to mark the endoplasmic reticulum (ER) (yellow in merged images) and is stained with CMAC, a vacuole stain (green in merged images). Cells were imaged four hours after induction with 100 nM β-estradiol. (C) Pearson Correlation of Wtf4^antidote^-mCherry and CMAC (vacuole, black line) and *sec63-YFP* (ER, blue line) from cells carrying a β-estradiol inducible *wtf4^antidote^-mCherry* allele, integrated *sec63-YFP* allele, and stained with CMAC, imaged four hours after induction with 100 nM β-estradiol. Wtf4^antidote^-mCherry and CMAC shows a Pearson coefficient of approximately 0.69, while Wtf4^antidote^-mCherry and Sec63-YFP shows -0.06. (D) A vegetatively growing haploid cell expressing a β-estradiol inducible *wtf4^antidote^-mCherry* allele (magenta in merged images) and a β-estradiol inducible *wtf4^poison^-GFP allele* (cyan in merged images) imaged four hours after induction with 100 nM β-estradiol. (E) Pearson correlation of Wtf4^antidote^-mCherry and Wtf4^poison^-GFP from cells carrying a β-estradiol inducible *wtf4^antidote^-mCherry* allele and a β-estradiol inducible *wtf4^poison^-GFP* allele 4 hours after induction with 100 nM β-estradiol shows a Pearson coefficient of approximately 0.64. All scale bars represent 2 µm.

**Figure 1-figure supplement 4.**
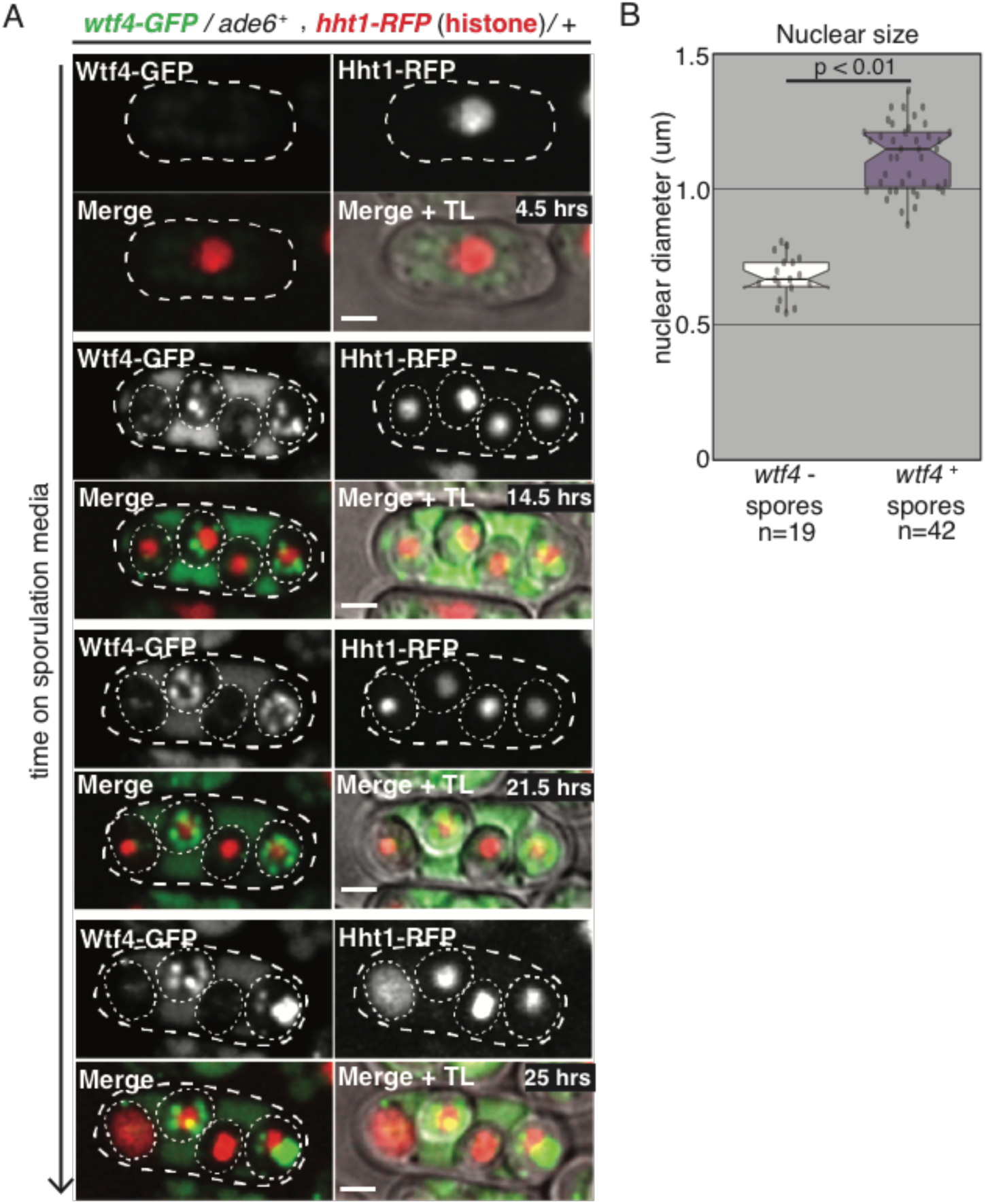
Spores destroyed by Wtf4^poison^ exhibit nuclear condensation followed by fragmentation. (A) A time course showing the nuclear phenotypes representative of gametogenesis starting from a *wtf4-GFP/ade6*^+^, *hht1-RFP/*+ diploid. The nucleus was visualized using the histone Hht1-RFP marker (red in merged images). The spores that inherit the *wtf4-GFP* allele can be distinguished from those that do not, because the *wtf4*^+^ spores have enriched GFP signal (green in merged images) from the accumulation of the Wtf4^antidote^. During spore development, the nuclei of the *wtf4-* spores appear to condense (14.5 hours) and, in some spores, fragment (25 hours). All scale bars represent 2 µm. (B) Quantification of nuclear diameter (µm) in *wtf4*^-^ and *wtf4*^+^ spores in asci produced by *wtf4-GFP*/*ade6*^+^, *hht1-RFP*/+ diploids, imaged two days after placement on SPA media. We excluded any nuclei that appeared to have already exploded (p< 0.01, t-test).

**Figure 2-figure supplement 1.**
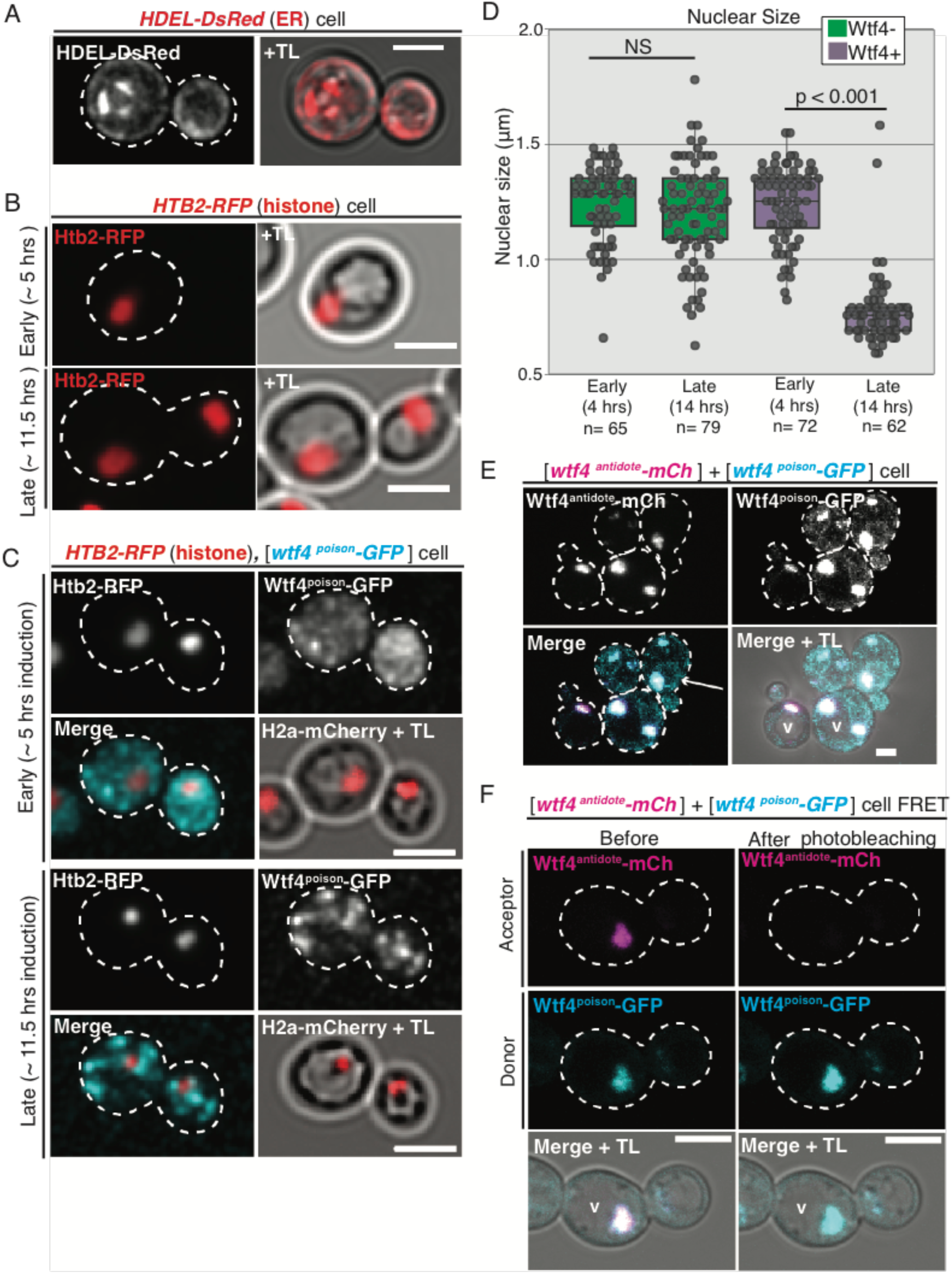
Wtf4^poison^ toxicity and Wtf4^antidote^ neutralization function in *S. cerevisiae*. (A) A vegetatively growing cell with an integrated HDEL-DsRed allele acting as an endoplasmic reticulum (ER) marker (red in merged image) imaged 4 hours after induction in SC galactose media. (B) A cell containing an integrated *HTB2-RFP* histone marker (red) ∼5 hours and ∼11.5 after switching to SC galactose (inducing) media. (C) A cell containing the *HTB2-RFP* histone marker (red in merged images) and carrying a [*URA3*] vector with a galactose inducible *wtf4^poison^-GFP* allele (cyan in merged images) ∼5 hours and ∼11.5 after switching to SC galactose (inducing) media. (D) Quantification of nuclear size (μm) in wild-type cells and in cells carrying the [*URA3*] vector with the galactose inducible *wtf4^poison^-GFP* allele 4 and 14 hours after being placed in SC galactose (inducing) media (p<0.01, t-test, ns= not significant). (E) A cell carrying both a [*TRP1*] vector with a β-estradiol inducible *wtf4^antidote^-mCherry* allele and a [*URA3*] vector with a β-estradiol inducible *wtf4^poison^-GFP* allele imaged 4 hours after being placed in 500 nM β-estradiol media. Both Wtf4 proteins colocalize in a large puncta next to the vacuole (v), but there is a faint circle of Wtf4^poison^-GFP (cyan in merged images) that is devoid of Wtf4^antidote^-mCherry (magenta in merged images, arrow). (F) Representative image of Acceptor Photobleaching <u>F</u>luorescence <u>R</u>esonance <u>E</u>nergy <u>T</u>ransfer (FRET) of Wtf4^poison^-GFP (the donor, cyan) and Wtf4^antidote^-mCherry (the acceptor, magenta) before and after photobleaching of Wtf4^antidote^-mCherry in cells carrying both a [*TRP1*] vector with a β-estradiol inducible *wtf4^antidote^-mCherry* allele and a [*URA3*] vector with a β-estradiol *wtf4^poison^-GFP* allele. FRET was assayed four hours after induction in 500 nM β-estradiol media. All scale bars represent 4 µm.

**Figure 2-figure supplement 2.**
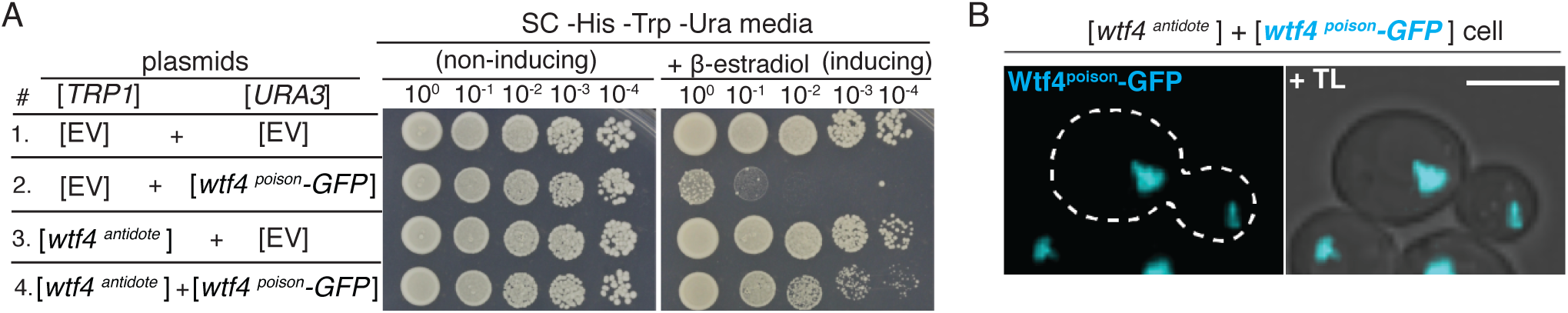
Wtf4^poison^ toxicity and Wtf4^antidote^ neutralization function in *S. cerevisiae*. (A) Spot assay of serial dilutions on non-inducing (SC *-*His -Trp -Ura) and inducing (SC -His -Trp -Ura + 500 nM β-estradiol) media. Each strain contains [*TRP1*] and [*URA3*] ARS CEN plasmids that are either empty (EV) or carry the indicated β-estradiol inducible *wtf4* alleles. (B) A haploid cell carrying a [*TRP1*] vector with an β-estradiol inducible *wtf4^antidote^* allele and a [*URA3*] vector with a β-estradiol inducible *wtf4^poison^-GFP* allele (cyan) imaged four hours after induction in 500 nM β-estradiol. Scale bar represents 4 µm.

**Figure 2-figure Supplement 3.**
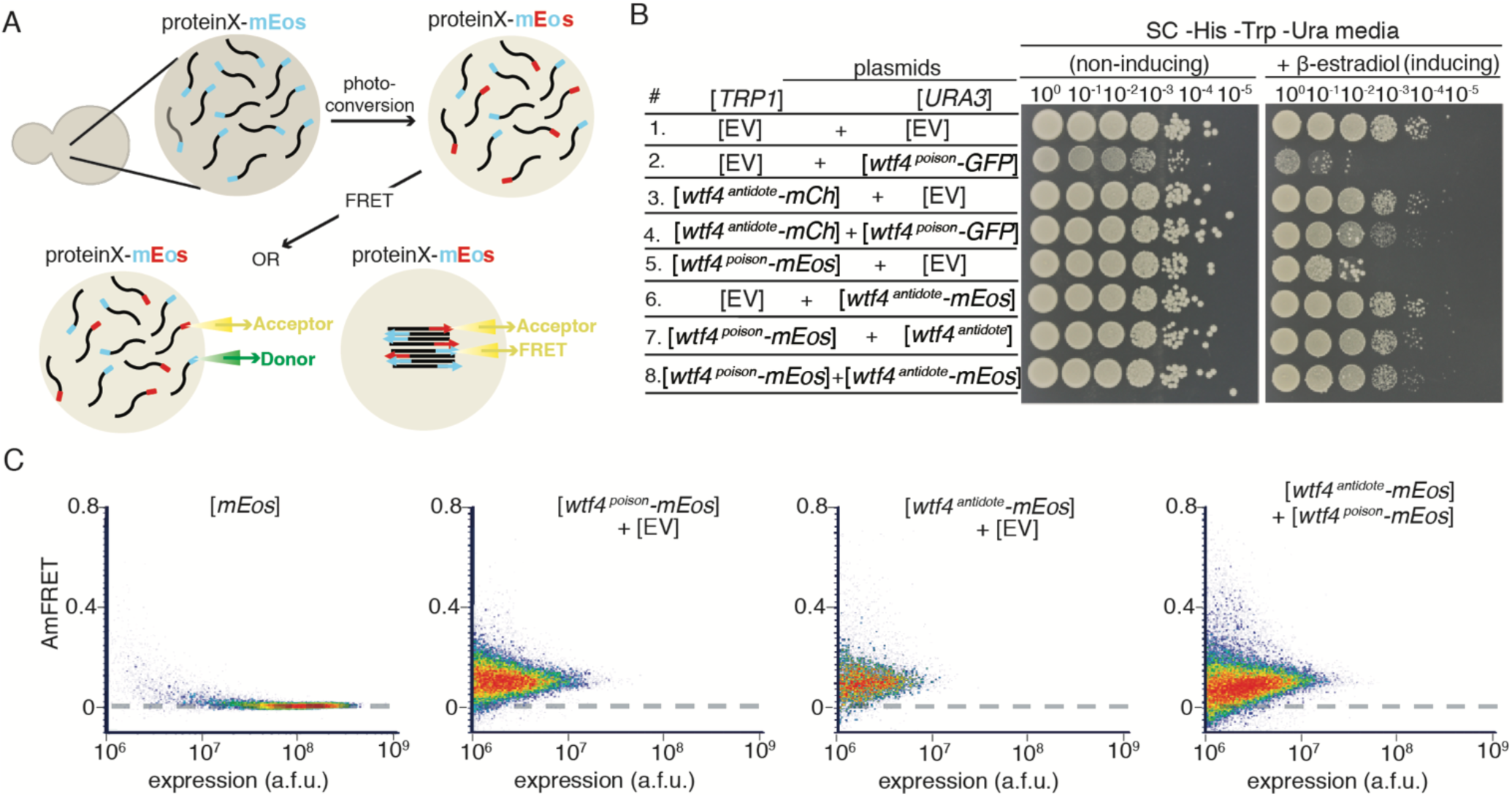
Wtf4^poison^ and Wtf4^antidote^ proteins assemble into aggregates. (A) Model of <u>D</u>istributed <u>A</u>mphifluoric FRET (DAmFRET) assay. (B) Spot assay of serial dilutions on non-inducing (SC *-*His -Trp -Ura) and inducing (SC -His -Trp -Ura + 500 nM β-estradiol) media. Each strain contains [*TRP1*] and [*URA3*] ARS CEN plasmids that are either empty (EV) or carry the indicated β-estradiol inducible *wtf4* alleles. (C) DAmFRET plots of flow cytometry data of cells carrying a [*URA3*] vector galactose inducible mEos3.1 (control for a monomeric protein population), cells carrying a [*TRP1*] vector with a β-estradiol inducible *wtf4^poison^-mEos3.1* allele and an empty [*URA3*] vector, cells carrying a [*URA3*] vector with a β-estradiol inducible *wtf4^antidote^-mEos3.1* allele and an empty [*TRP1*] vector, and cells carrying a [*URA3*] vector with a β-estradiol inducible *wtf4^antidote^-mEos3.1* allele and a [*TRP1*] vector with a β-estradiol inducible *wtf4^poison^-mEos3.1*. Cells carrying vectors with the β-estradiol inducible Wtf4-mEos3.1 proteins were assayed four hours after induction in 500 nM β-estradiol. Cells carrying the galactose inducible mEos3.1 control were assayed 16 hours after induction in SC galactose media.

**Figure 3-figure supplement 1.**
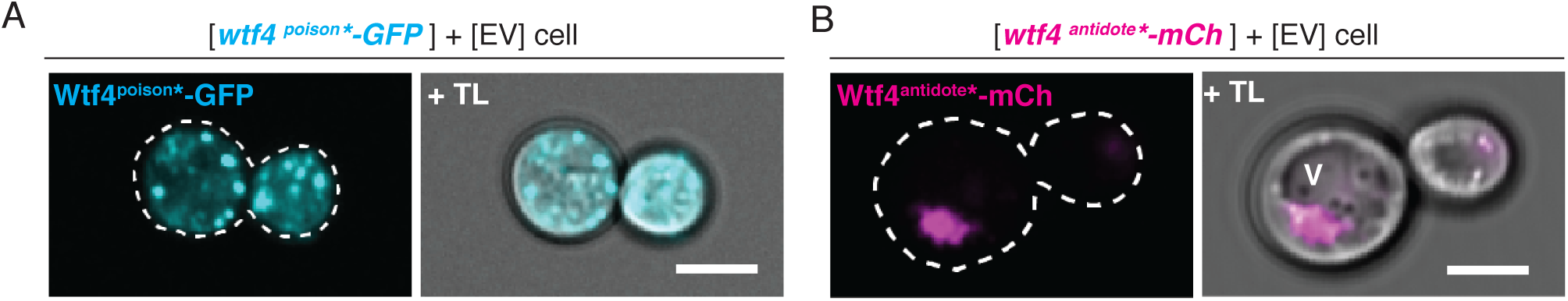
Wtf4^poison^* and Wtf4^antidote^* have the same localization as the wild-type Wtf4 proteins in *S. cerevisiae*. (A) Representative image of a haploid cell carrying a [*URA3*] vector with a β-estradiol inducible *wtf4^poison^-GFP* allele (cyan) and an empty [*TRP1*] vector. (B) Representative image of a haploid cell with carrying a [*TRP1*] vector with a β-estradiol inducible *wtf4^antidote^-mCherry* allele (magenta) and an empty [*URA3*] vector. In both experiments, the cells were imaged ∼4 hours after induction in 500 nM β-estradiol. All scale bars represent 4 µm.

**Figure 4-figure supplement 1.**
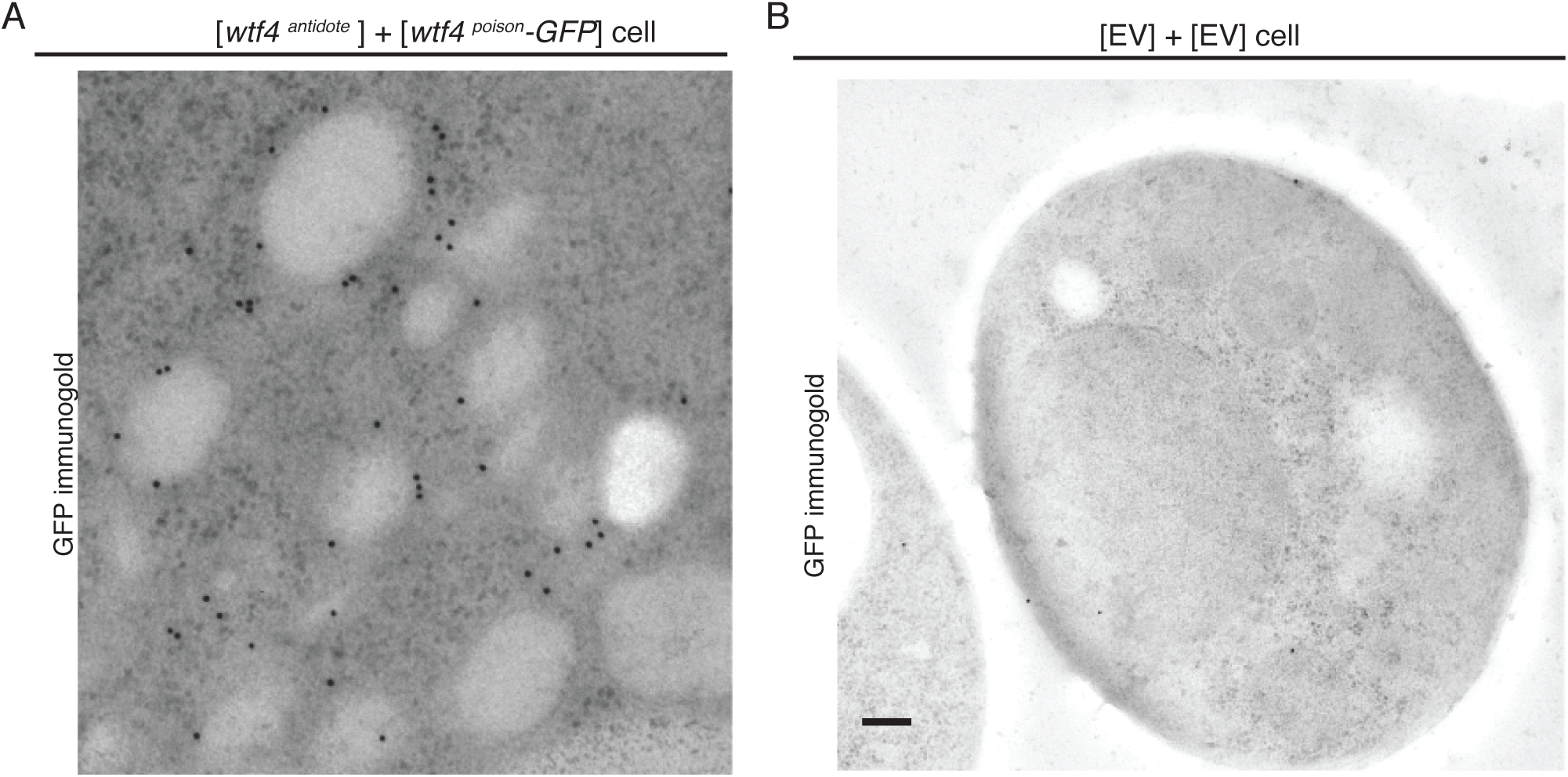
Cells expressing Wtf4^poison^ and Wtf4^antidote^ or Wtf4^antidote^ have increased vesicles that cluster within the Wtf4 aggregate. (A) A cropped, magnified immuno-gold <u>T</u>ransmission <u>E</u>lectron <u>M</u>icroscopy (TEM) tomograph of a region of a haploid cell carrying a [*TRP1*] vector with a β-estradiol inducible *wtf4^antidote^* allele and a [*URA3*] vector with a β-estradiol inducible *wtf4^poison^-GFP* allele. A mono-clonal antibody against GFP was used. Immunogold particles (black dots) are enriched in a cluster near light staining organelles. (B) Representative Immuno-gold TEM tomograph of a cell carrying an empty [*TRP1*] vector and an empty [*URA3*] vector. The same antibody was used as in A, but very few particles are seen, suggesting little background staining is present. Both samples were processed after ∼4 hours of induction in 500 nM β-estradiol. Scale bar represents 0.2 µm.

**Figure 4-figure supplement 2.**
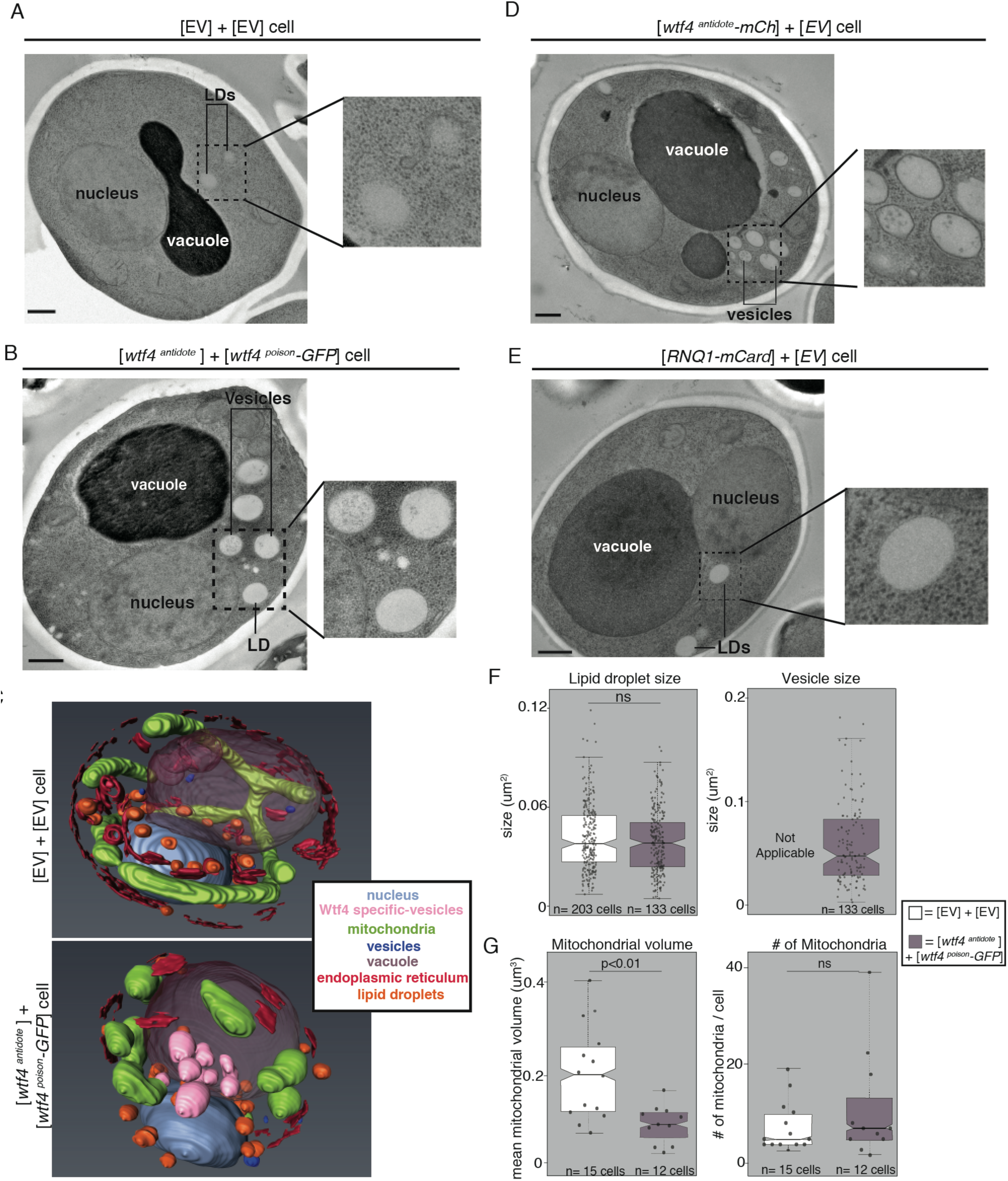
Cells expressing Wtf4^poison^ and Wtf4^antidote^ or Wtf4^antidote^ have increased vesicles that cluster within the Wtf4 aggregate. (A) <u>T</u>ransmission <u>E</u>lectron <u>M</u>icroscopy (TEM) tomograph of a haploid cell carrying an empty [*TRP1*] vector and an empty [*URA3*] vector. (B) TEM tomograph of a haploid cell carrying a [*TRP1*] vector with a *wtf4^antidote^* allele and a [*URA3*] vector with a *wtf4^poison^-GFP* allele. (C) Models generated using array tomography, a process that utilizes serial slices and <u>S</u>canning <u>E</u>lectron <u>M</u>icroscopy (SEM) to generate a 3D reconstruction of the cell. Top is the reconstruction of a haploid cell carrying an empty [*TRP1*] vector and an empty [*URA3*] vector. The bottom is the 3D reconstruction of a haploid cell carrying a [*TRP1*] vector with a *wtf4^antidote^* allele and a [*URA3*] vector with a *wtf4^poison^-GFP* allele. The key shows the colors representative of the cellular structures in the images. (D) TEM tomograph of a haploid cell carrying a [*TRP1*] vector with a *wtf4^antidote^-mCherry* allele and an empty [*URA3*] vector. (E) TEM tomograph of a haploid cell carrying a [*TRP1*] vector with a *RNQ1-mCardinal* allele and a [*URA3*] vector with a *wtf4^poison^-GFP* allele. (F) Quantification of the size of lipid droplets (left) or vesicles (right) in two samples: cells carrying an empty [*TRP1*] vector and an empty [*URA3*] vector (EV, white, n=203) and cells carrying a [*TRP1*] vector with a *wtf4^antidote^* allele and a [*URA3*] vector with a *wtf4^poison^-GFP* allele (purple, n=133 cells). (G) Quantification of the average volume of mitochondria per cell (left) or number of mitochondria per cell (right) in two samples: cells carrying an empty [*TRP1*] vector and an empty [*URA3*] vector (EV, white, n=15 cells) and cells carrying a [*TRP1*] vector with a *wtf4^antidote^* allele and a [*URA3*] vector with a *wtf4^poison^-GFP* allele (purple, n=12 cells). (ns= not significant) (p< 0.01, t-test). All samples were processed after ∼4 hours of induction in 500 nM β-estradiol. Scale bar represents 0.5 µm.

**Figure 4-figure supplement 3.**
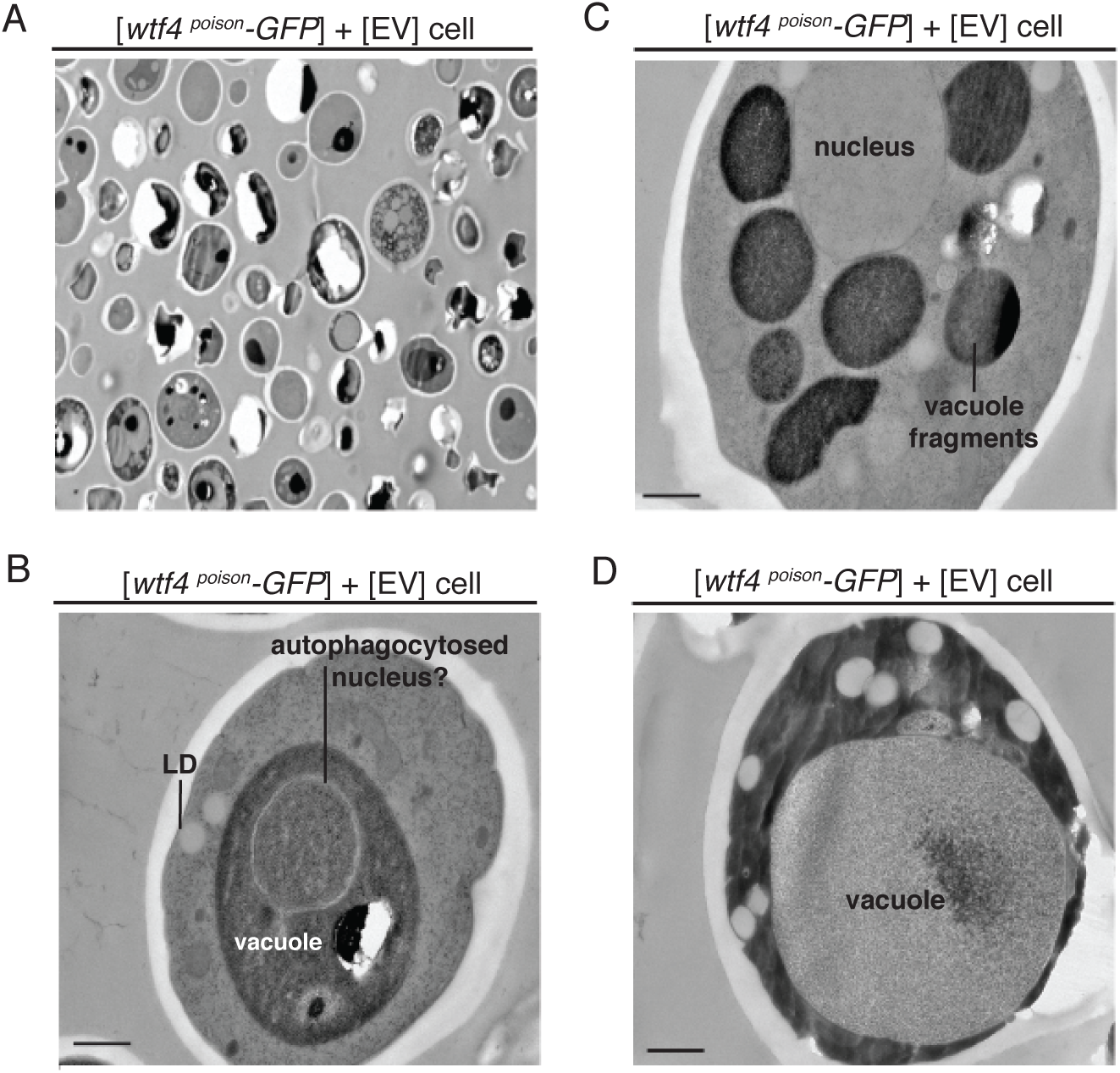
Cells expressing only Wtf4^poison^ have variable phenotypes. (A-D) are representative <u>T</u>ransmission <u>E</u>lectron <u>M</u>icroscopy (TEM) tomographs of cells carrying a vector with a [*URA3*] vector with a β-estradiol inducible *wtf4^poison^-GFP* and an empty [*TRP1*] vector. Cells were processed four hours after induction with 500 nM β-estradiol. (A) TEM micrograph at a lower magnification, showing a field of cells. Many of the cells appear dead and have lost integrity through the TEM process. There are few cells surviving long enough to have recognizable phenotypes. We grouped the remaining cells into phenotype classes listed in B-D. (B) One phenotype of the remaining cells is increased autophagy. The cell shown has an organelle, presumed to be the nucleus, inside the darker staining vacuole. (C) Some cells show numerous, fragmented vacuoles. (D) Others show enlarged vacuoles, with minimal, darker staining cytoplasm. All scale bars represent 0.5 µm. LD= lipid droplet.

**Figure 5-figure supplement 1.**
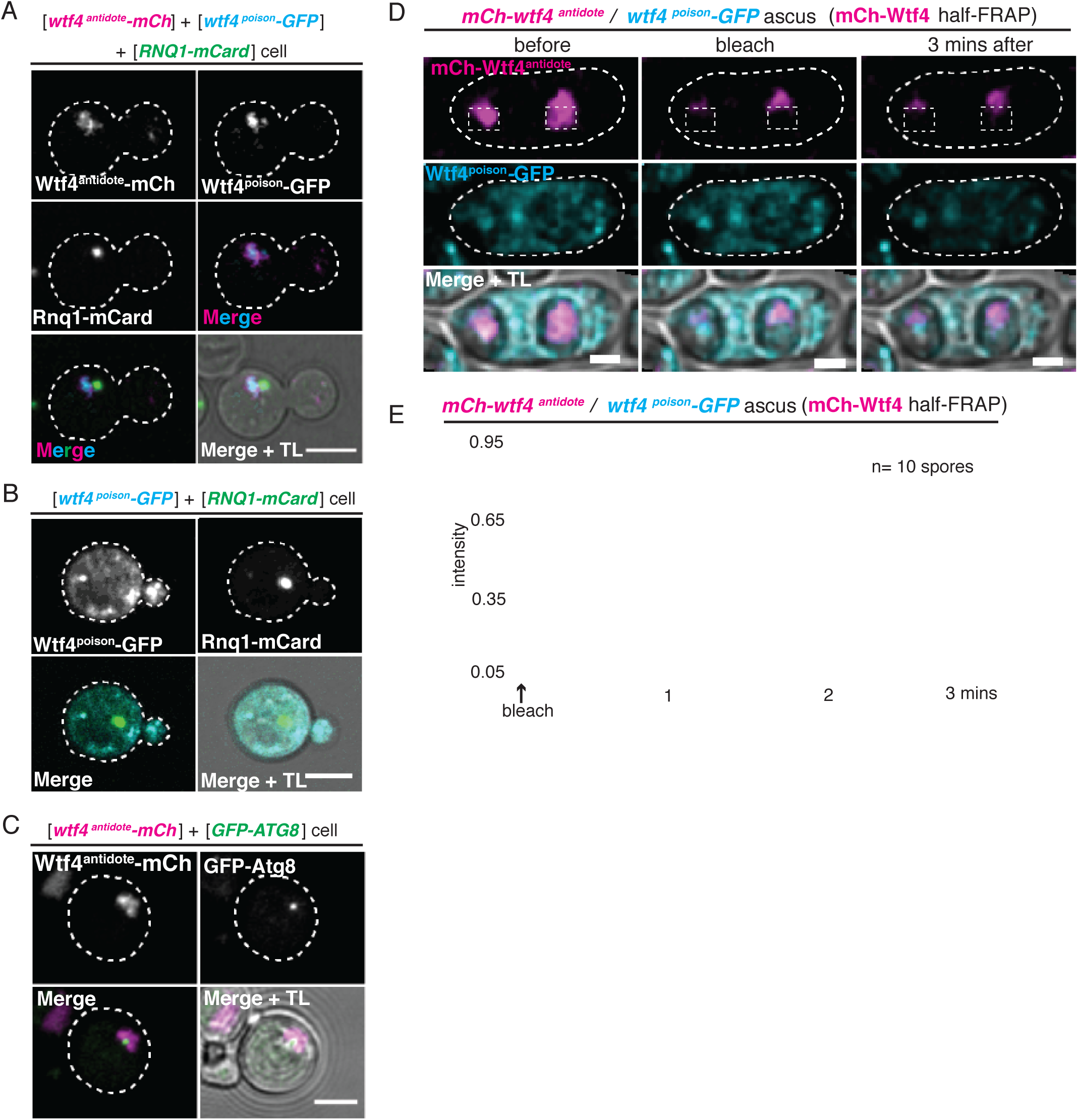
The Wtf4 proteins colocalize at the Insoluble <u>P</u>rotein <u>D</u>eposit (IPOD). (A) Representative image of a cell carrying a [*URA3*] vector with a β-estradiol inducible *wtf4^poison^-GFP* allele, a [*TRP1*] vector with a β-estradiol inducible *wtf4^antidote^-mCherry* allele (magenta in merged images), and a [*LEU2*] vector with a β-estradiol inducible *RNQ1-mCardinal* allele (green in merged images) acting as an IPOD marker. The images were acquired after ∼4 hours in 500 nM β-estradiol media. (B) Representative image of a cell carrying a [*URA3*] vector with a β-estradiol inducible *wtf4^poison^-GFP* allele (cyan in merged images), an empty [*TRP1*] vector, and a [*LEU2*] vector with a β-estradiol inducible *RNQ1-mCardinal* acting as an IPOD marker (green in merged images). Cells were images ∼4 hours in 500 nM β-estradiol media. (C) Representative image of a cell carrying a [*LEU2*] vector with a galactose inducible *wtf4^antidote^-mCherry* (magenta in merged images) and a [*URA3*] vector with a *GFP-ATG8* allele under its endogenous promoter acting as a Pre-Autophagosomal Site (PAS) marker (green in merged images). Acquired after ∼4 hours in SC galactose media. (D) Representative image of half-<u>F</u>luorescence <u>R</u>ecovery <u>A</u>fter <u>P</u>hotobleaching (half-FRAP) of the Wtf4^antidote^-mCherry (magenta) aggregate in asci that were generated from heterozygous *mCherry-wtf4/wtf4^poison^-GFP* diploids. (E) Quantification of the FRAP data shown in D. Cells were imaged for 3 minutes after bleaching and very little recovery was seen, suggesting the mCherry-Wtf4^antidote^ protein is stable. All scale bars represent 4 µm.

**Figure 5-figure supplement 2.**
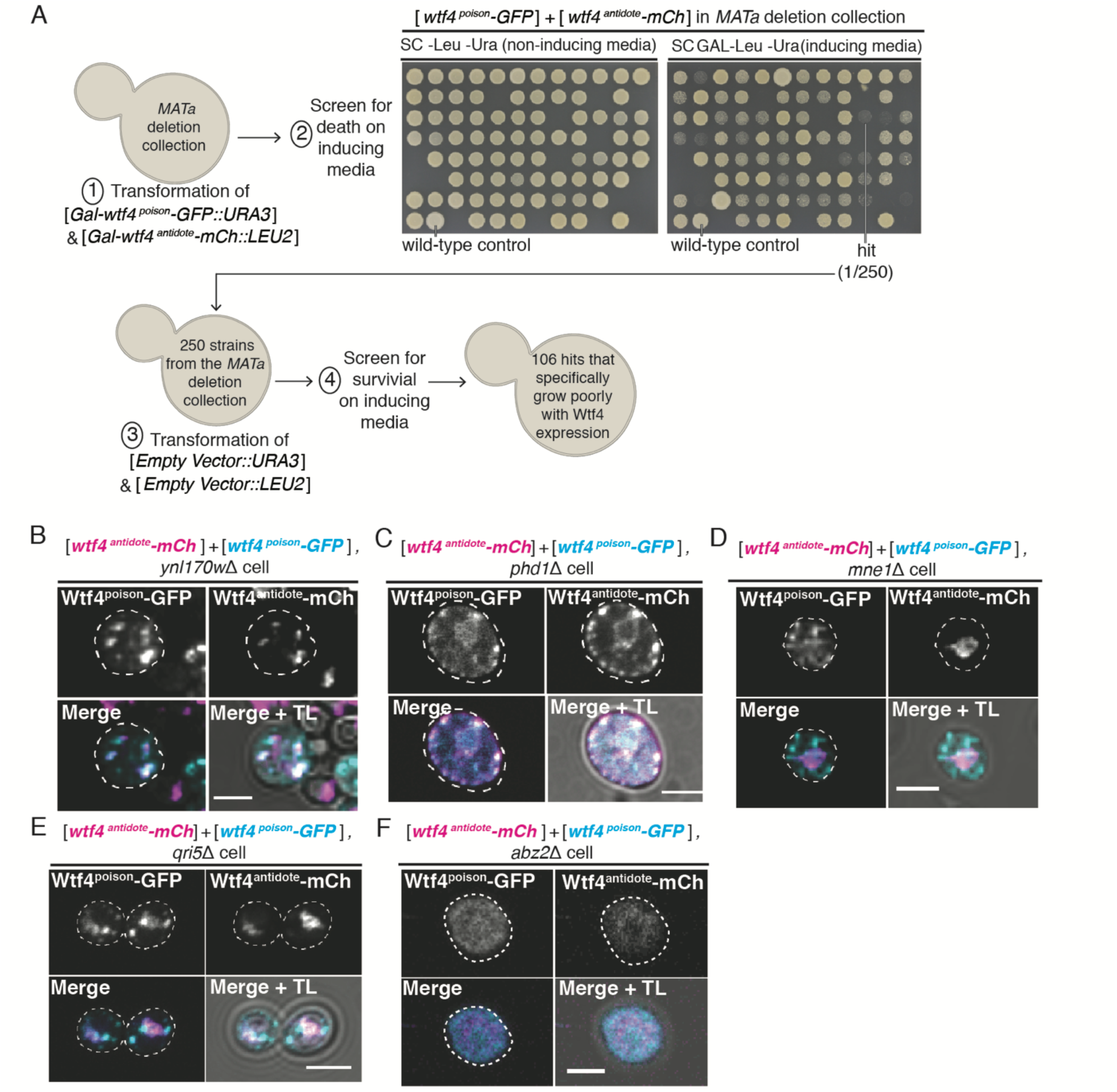
A screen of the *S. cerevisiae* deletion collection to identify genes necessary for survival upon Wtf4^poison^ and Wtf4^antidote^ expression. (A) Cartoon of the screen designed to identify genes necessary for survival of cells after co-induction of the Wtf4 poison and antidote proteins. Representative images of a plate from the screen are included. The control is a wild-type cell carrying a [*LEU2*] vector with a galactose inducible *wtf4^antidote-^ mCherry* and a [*URA3*] vector with a galactose inducible *wtf4^poison^-GFP* allele. The 250 strains that showed reduced growth in comparison to the control on SC galactose (inducing) media were deemed as hits. Those 250 strains were re-assayed from the *MATa* deletion collection with only [*LEU2*] and [*URA3*] empty vectors. (B)-(F) are representative images of vegetativelygrowing, haploid cells carrying a [*LEU2*] vector with a galactose inducible *wtf4^antidote^-mCherry* (magenta in merged images) and a [*URA3*] vector with a galactose inducible *wtf4^poison^-GFP* allele (cyan in merged images). Cells were imaged four hours in galactose (inducing) media. (B) *ynl170w*Δ and (C) *phd1*Δ are representative of multiple samples (81/106) that showed the phenotype of the Wtf4 proteins localizing as dispersed puncta throughout the cell. (D) *mne1*Δ, and (E) *qrl51*Δ, are representative of multiple samples (5/106) that showed the phenotype of the Wtf4^antidote^-mCherry protein localizing as a puncta, while the Wtf4^poison^-GFP protein looked more dispersed. (F) *abz2*Δ, is representative of multiple samples (20/106) that showed the Wtf4^antidote^-mCherry and Wtf4^poison^-GFP proteins localizing as a soluble haze throughout the cell. All scale bars represent 4 µm.

**Figure 5-Figure Supplement 3.**
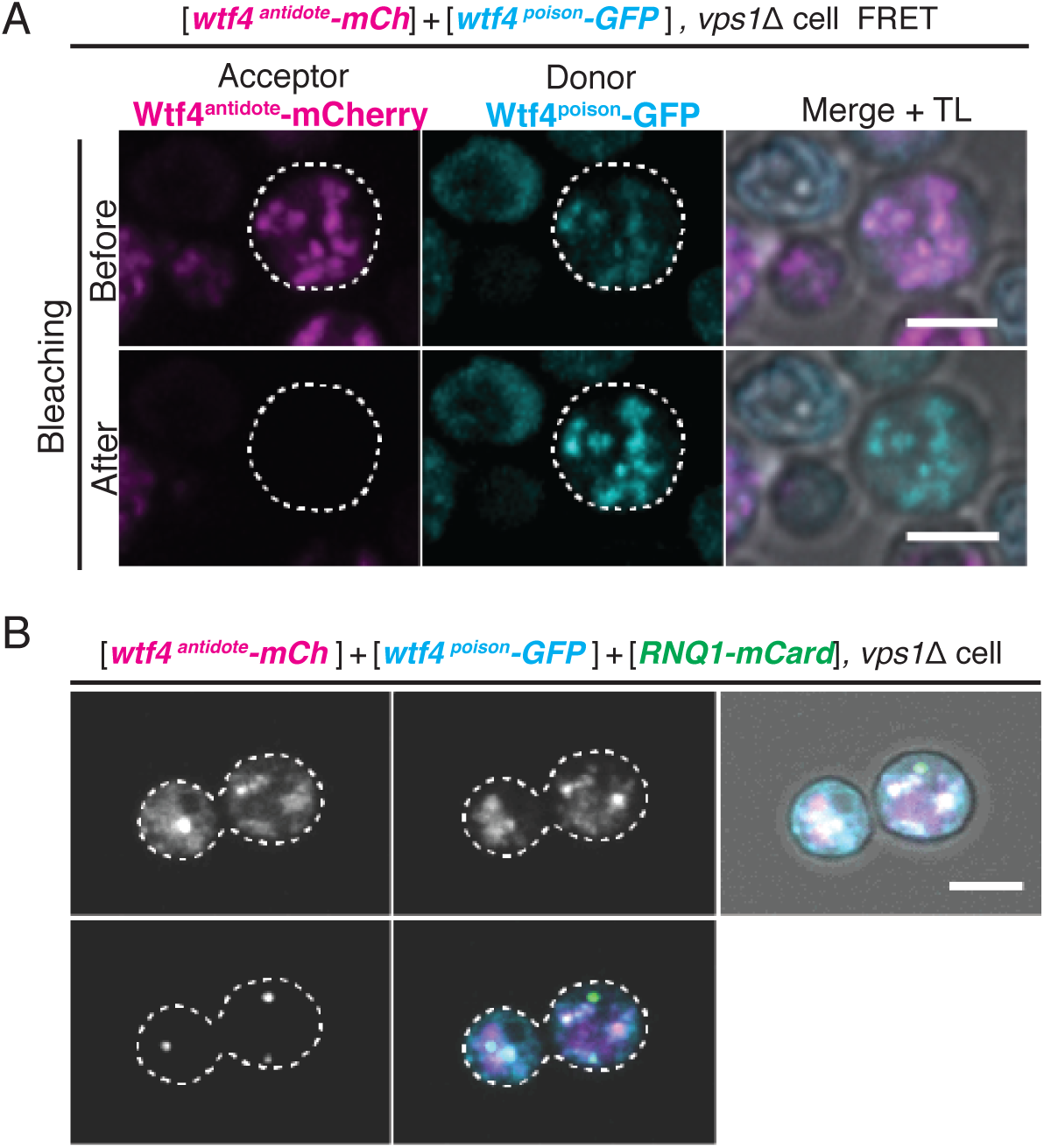
Vps1 is necessary for the recruitment of Wtf4 proteins to the Insoluble Protein Deposit (IPOD). (A) Representative image of acceptor photobleaching Fluorescence Resonance Energy Transfer (FRET) of Wtf4^antidote^-mCherry and Wtf4^poison^-GFP in *vps1*Δ cells carrying a carrying a [*TRP1*] vector with a β-estradiol inducible *wtf4^antidote^-mCherry* (magenta in merged images) allele and a [*URA3*] vector with a β-estradiol inducible *wtf4^poison^-GFP* (cyan in merged images) allele. After photobleaching of Wtf4^antidote^-mCherry (the acceptor), Wtf4^poison-^GFP (the donor) signal increases. (B) Representative image of a vegetatively growing haploid *vps1*Δ cell carrying a [*TRP1*] vector with a β-estradiol inducible *wtf4^antidote^-mCherry* (magenta in merged images) allele, a [*URA3*] vector with a β-estradiol inducible *wtf4^poison^-GFP* (cyan in merged images) allele, and a [*LEU2*] vector with the β-estradiol inducible IPOD marker, *RNQ1-mCardinal* (green in merged images). All cells were imaged four hours after induction in 500 nM β-estradiol media. All scale bars represent 4 µm.

**Supplemental Table 1.**
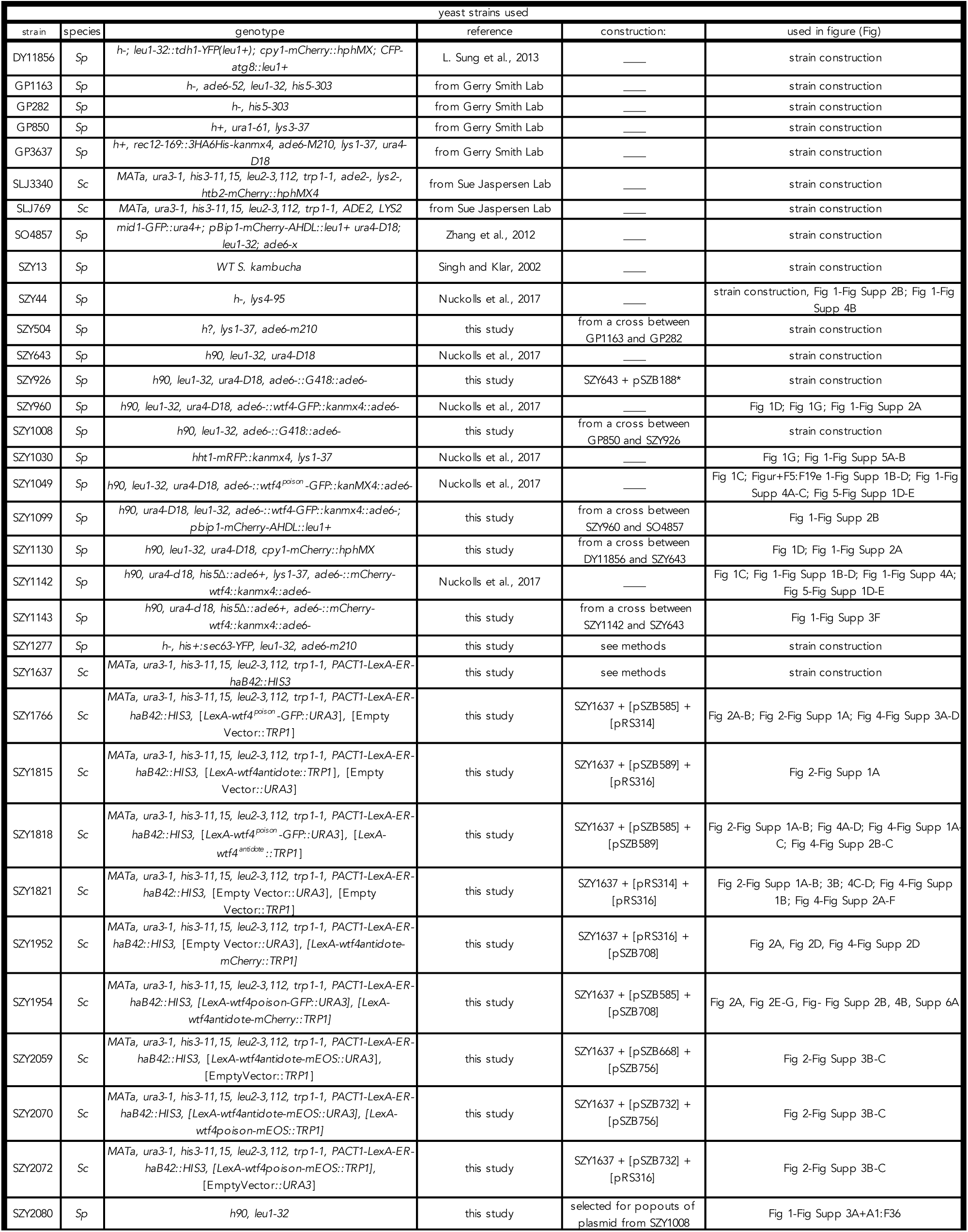
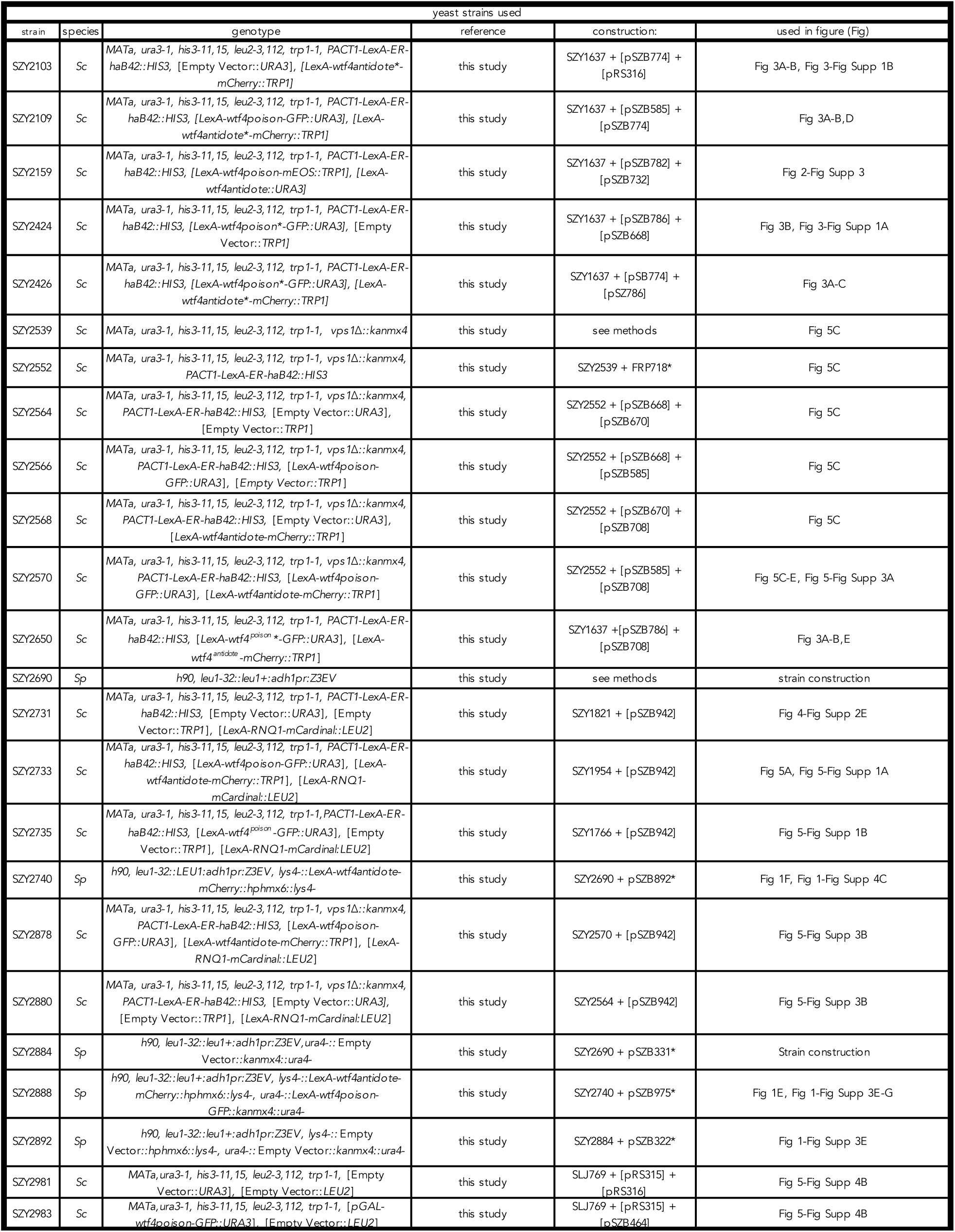
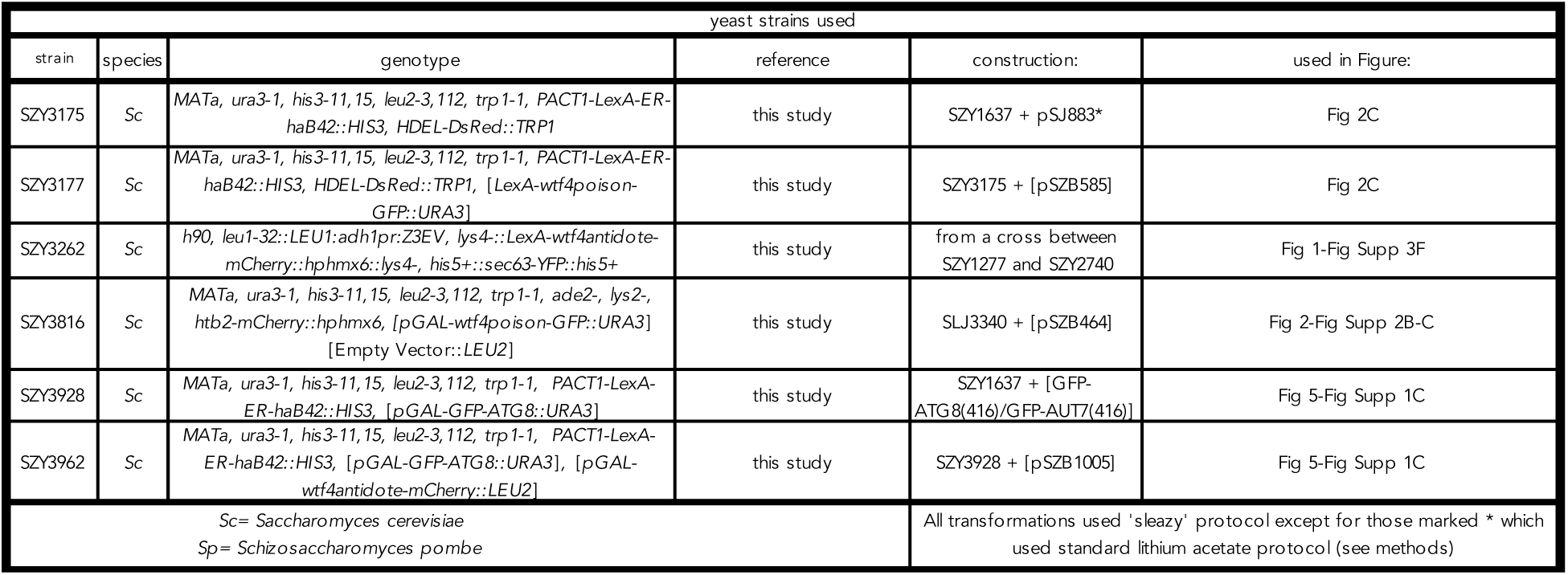
**Yeast Strains Used**

**Supplemental Table 2.** **Plasmids Used**

| plasmids used |  |  |
| --- | --- | --- |
| plasmid | description | Reference |
| P03233DS | pBT3-STE::LEU2, bait vector Y2H | DUALsystems Biotech |
| V08_mC | pGAL-mCardinal::LEU2 | from the Halfmann lab |
| FRP718 | PACT1-LexA-ER-haB42::HIS3 | Ottoz et al., 2014 |
| FRP1642 | PTEF-TTEF-insul-4LEXABOX-pCYC1::hphMX6 | Ottoz et al., 2014 |
| GFP-ATG8(416)/GFP-AUT7(416) | GFP-ATG8::URA3 | Guan et al., 2001 |
| pDK20 | pGAL::URA3 | DasGupta et al., 1998 |
| pDK412 | pGAL-RNQ1-RFP::LEU2 | D. Kryndushkin et al., 2012 |
| pFS461 | adh1pr:Z3EV::Leu1+ | Ohira et al., 2017 |
| pFS478 | kanMX4::pZ3EV | Ohira et al., 2017 |
| pRS314 | Empty Vector::TRP1 | Sikorski et al., 1989 |
| pRS315 | Empty Vector::LEU2 | Sikorski et al., 1989 |
| pRS316 | Empty Vector::URA3 | Sikorski et al., 1989 |
| pSJ883 | HDEL-DsRed::TRP1 | Friederichs et al., 2011 |
| pYM41 | pFA6-eYFP-his3MX6 | Janke et al., 2004 |
| pSZB188 | Empty Vector::kanMX4::ade6- | Nuckolls et al., 2017 |
| pSZB199 | wtf4 (coding sequence)::kanMX4::ade6- | this study |
| pSZB203 | wtf4-GFP::kanMX4::ade6- | Nuckolls et al., 2017 |
| pSZB248 | mcherry-wtf4::kanMX4::ade6- | Nuckolls et al., 2017 |
| pSZB257 | wtf4<sup>poison</sup>-GFP::kanMX4::ade6- | Nuckolls et al., 2017 |
| pSZB260 | wtf4<sup>antidote</sup>-GFP::kanMX4::ade6- | this study |
| pSZB322 | Empty Vector::hphMX6::lys4- | Bravo Núñez et al., 2018 |
| pSZB331 | Empty Vector::kanMX4::ura4- | Bravo Núñez et al., 2020 |
| pSZB386 | Empty Vector::hphMX6::ade6- | Bravo Núñez et al., 2018 |
| pSZB388 | pBT3-STE-wtf4<sup>poison</sup>::LEU2 | this study |
| pSZB392 | pGAL-wtf4<sup>poison</sup>::URA3 | this study |
| pSZB464 | pGAL-wtf4<sup>poison</sup>-GFP::URA3 | this study |
| pSZB463 | pGAL-wtf4<sup>poison</sup>-GFP::TRP1 | this study |
| pSZB497 | pGAL-wtf4<sup>antidote</sup>::TRP1 | this study |
| pSZB585 | LexA-wtf4<sup>poison</sup>-GFP::URA3 | this study |
| pSZB589 | LexA-wtf4<sup>antidote</sup>::TRP1 | this study |
| pSZB668 | LexA-Empty Vector::TRP1 | this study |
| pSZB670 | LexA-Empty Vector::URA3 | this study |
| pSZB700 | LexA-mCherry-wtf4<sup>antidote</sup>::TRP1 | this study |
| pSZB708 | LexA-wtf4<sup>antidote</sup>-mCherry::TRP1 | this study |
| pSZB732 | LexA-wtf4<sup>poison</sup>-mEOS::TRP1 | this study |
| pSZB756 | LexA-wtf4<sup>antidote</sup>-mEOS::URA3 | this study |
| pSZB774 | LexA-wtf4<sup>antidote</sup>*-mCherry::TRP1 | this study |
| pSZB782 | LexA-wtf4<sup>antidote</sup>::URA3 | this study |
| pSZB786 | LexA-wtf4<sup>poison</sup>*-GFP::URA3 | this study |
| pSZB898 | pGAL-mCardinal::LEU2 | this study |
| pSZB891 | wtf4 (coding sequence)-mCherry::kanMX4::ura4- | this study |
| pSZB892 | LexA-wtf4<sup>antidote</sup>-mCherry::lys4::hphMX6 | this study |
| pSZB942 | LexA-RNQ1-mCardinal::LEU2 | this study |
| pSZB975 | LexA-wtf4<sup>poison</sup>-GFP::ura4::kanMX4 | this study |
| pSZB1005 | pGAL-wtf4<sup>antidote</sup>-mCherry::LEU2 | this study |
| pSZB1120 | pGAL-wtf4<sup>antidote</sup>-mEos::URA3 | this study |
| rhx1389 | pGAL-wtf4<sup>poison</sup>-mEos::URA3 | this study |

**Supplemental Table 3.**
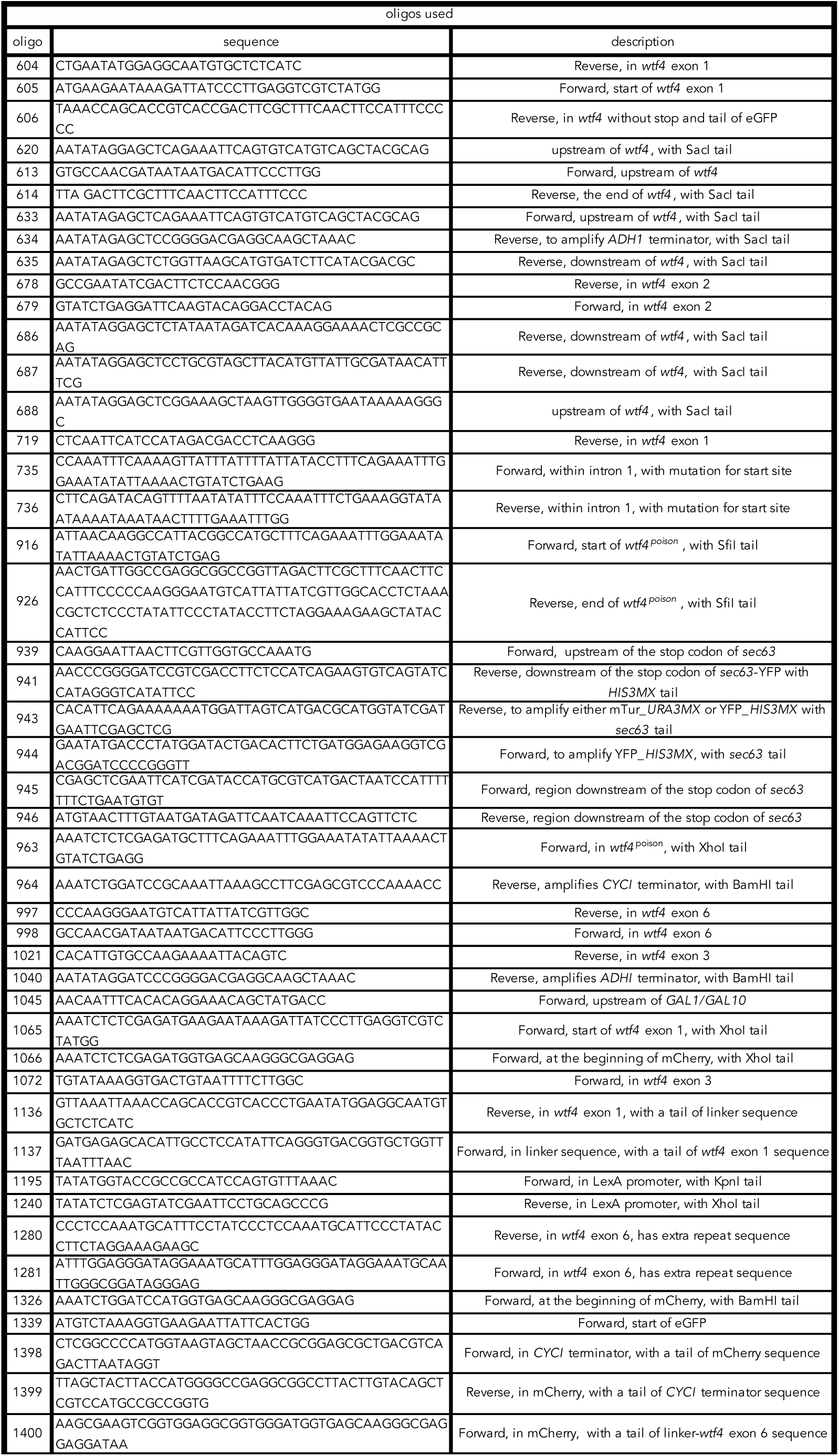
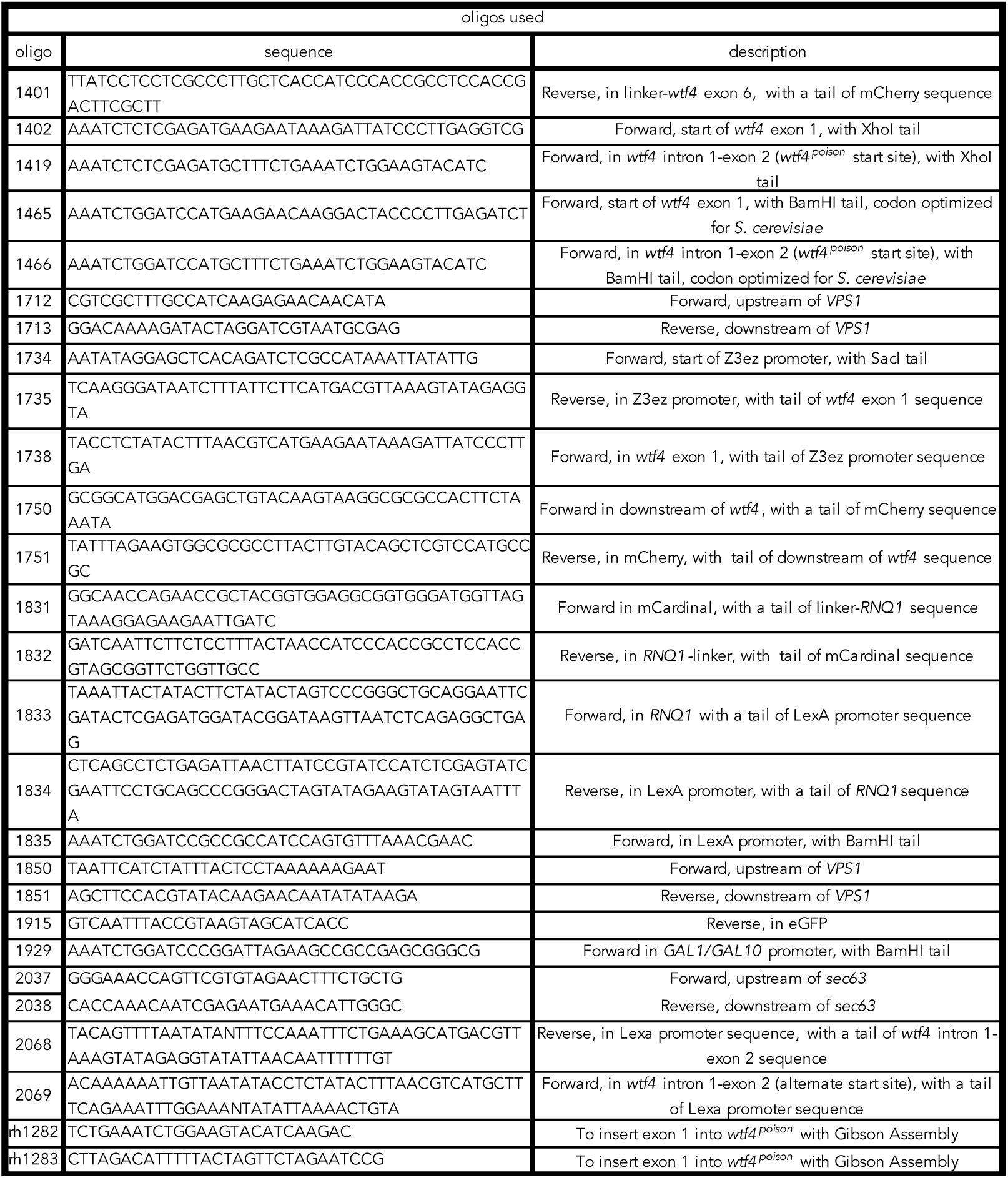
**Oligos Used**

## References

1. J. Adler and I. Parmryd, Quantifying colocalization by correlation: The Pearson correlation coefficient is superior to the Mander’s overlap coefficient. Cytometry 77A: 733–742 (2010).

2. T. Akera, L. Chmátal, E. Trimm, K. Yang, C. Aonbangkhen, D.M. Chenoweth, et al., Spindle asymmetry drives non-Mendelian chromosome segregation. Science 358(6363):668–72 (2017).

3. M. Arrasate, S. Mitra, E.S. Schweitzer, M.R. Segal, S. Finkbeiner, Inclusion body formation reduces levels of mutant huntingtin and the risk of neuronal death. Nature 431(7010):805–810 (2004).

4. K. Bagola, T. Sommer, Protein quality control: on IPODs and other JUNQ. Current Biology 18(21):R1019–21 (2008).

5. H. Bauer, N. Véron, J. Willert, and B.G. Herrmann, The t-complex-encoded guanine nucleotide exchange factor Fgd2 reveals that two opposing signaling pathways promote transmission ratio distortion in the mouse. Genes & development 21(2):143–147 (2007).

6. H. Bauer, S. Schindler, Y. Charron, J. Willert, B. Kusecek, B.G. Herrmann, The nucleoside diphosphate kinase gene *Nme3* acts as quantitative trait locus promoting non-Mendelian inheritance. PLOS Genetics 8(3):e1002567. (2012)

7. I. Belevich, M. Joensuu, D. Kumar, H. Vihinen and E. Jokitalo, Microscopy Image Browser: A platform for segmentation and analysis of multidimensional datasets. PLOS Biology 14(1):e1002340 (2016).

8. M.A. Bravo Núñez, I.M. Sabbarini, M.T. Eickbush, Y. Liang, J.J. Lange, A.M. Kent, S.E. Zanders, Dramatically diverse *S. pombe wtf* meiotic drivers all display high gamete-killing efficiency. PLOS Genetics (2020) in press.

9. M.A. Bravo Núñez, J.J. Lange, S.E. Zanders, A suppressor of a *wtf* poison-antidote meiotic driver acts via mimicry of the driver’s antidote. PLOS Genetics 14(11):e1007836 (2018).

10. M.A. Bravo Núñez, N. Nuckolls, S.E. Zanders, Genetic Villains: Killer Meiotic Drivers. Trends in Genetics 34(6):424–433. (2018).

11. A. Burt and A. Crisanti, Gene drive: Evolved and Synthetic. ACS Chem. Biol. 13:343–346 (2018).

12. A. Burt, Heritable strategies for controlling insect vectors of disease. Philosophical Transactions of the Royal Society B 369(1645):20130432 (2014).

13. A. Burt, R. Trivers, Genes in conflict: the biology of selfish genetic elements. (Belknap Press of Harvard University Press, Cambridge, Mass., 2006), pp. viii, 602 p., 608 p. of plates.

14. D. Carmona-Gutierrez, T. Eisenberg, S. Buttner, C. Meisinger, G. Kroemer F. Madeo, Apoptosis in yeast: triggers, pathways, subroutines. Cell Death and Differentiation 17:763– 773 (2010).

15. J. Chen, J. Ding, Y. Ouyang, H. Du, J. Yang, K. Cheng, et al., A triallelic system of S5 is a major regulator of the reproductive barrier and compatibility of indica–japonica hybrids in rice. Proceedings of the National Academy of Sciences of the United States of America 105(32):11436–41. (2008).

16. B. Chen, M. Retzlaff, T. Roos, J. Frydman, Cellular Strategies of Protein Quality Control. Cold Spring Harb Perspectives in Biology 3(8):a004374 (2011).

17. M. Coelho, S.J. Lade, S. Alberti, T. Gross, I.M. Tolić, Fusion of protein aggregates facilitates asymmetric damage segregation. PLOS Biology 12(6):e1001886 (2014).

18. M.R. Cookson and M. van der Brug, Cell systems and the toxic mechanism(s) of alpha-synuclein. Exp Neurol. 209(1):5–11. (2007).

19. J.F. Crow, Why is Mendelian segregation so exact? Bioessays 13:305–312 (1991).

20. H.J. Dalstra, K. Swart, A.J. Debets, S.J. Saupe, R.F. Hoekstra, Sexual transmission of the [Het-S] prion leads to meiotic drive in Podospora anserina. Proceedings of the National Academy of Sciences of the United States of America 100(11):6616–21. (2003).

21. H.J. Dalstra, R. van der Zee, K. Swart, R.F. Hoekstra, S.J. Saupe, A.J. Debets, Non-mendelian inheritance of the HET-s prion or HET-s prion domains determines the het-S spore killing system in Podospora anserina. Fungal Genetics and Biology 42(10):836–47 (2005).

22. S.K. Dasgupta, S. Jain, D. Kaushal and A.K. Tyagi, Expression systems for study of mycobacterial gene regulation and development of recombinant BCG vaccines. Biochem. Biophys. Res. Commun. 246(3):797–804 (1998).

23. R.K. Dawe, E.G. Lowry, J.I. Gent, M.C. Stitzer, K.W. Swentowsky, D.M. Higgins, et al., A kinesin-14 motor activates neocentromeres to promote meiotic drive in maize. Cell 173(4):839–50.e18.

24. J.P. Didion, A.P. Morgan, A.M. Clayshulte, R.C. McMullan, L. Yadgary, P.M. Petkov, et al., A multi-megabase copy number gain causes maternal transmission ratio distortion on mouse chromosome 2. PLOS Genetics 11(2):e1004850. (2015).

25. K.A. Dyer, B. Charlesworth, J. Jaenike, Chromosome-wide linkage disequilibrium as a consequence of meiotic drive. Proceedings of the National Academy of Sciences United States of America 104(5):1587–1592 (2007).

26. M.T. Eickbush, J.M. Young, S.E. Zanders, Killer Meiotic Drive and Dynamic Evolution of the *wtf* Gene Family. Molecular Biology and Evolution 36(6):1201–1214 (2019).

27. R. Elble, A simple and efficient procedure for transformation of yeasts. Biotechniques 13(1):18–20 (1992).

28. A.H. Enyenihi and W.S. Saunders, Large-Scale Functional Genomic Analysis of Sporulation and Meiosis in *Saccharomyces cerevisiae*. Genetics 163(1):47–54 (2003).

29. K.M. Esvelt, A.L. Smidler, F. Catteruccia, G.M. Church, Concerning RNA-guided gene drives for the alteration of wild populations. eLife 3:e03401 (2014).

30. J.M. Friederichs, S. Ghosh S, C.J. Smoyer, S. McCroskey, B.D. Miller, K.J. Weaver, K.M. Delventhal, J. Unruh, B.D. Slaughter, S.L. Jaspersen, The SUN protein Mps3 is required for spindle pole body insertion into the nuclear membrane and nuclear envelope homeostasis. PLOS Genetics 7(11):e1002365 (2011).

31. B. Fuchs, E. Mylonakis, Our paths might cross: the role of the fungal cell wall integrity pathway in stress response and cross talk with other stress response pathways. Eukaryotic Cell 8(11):1616–25. (2009).

32. V.M. Gantz, N. Jasinskiene, O. Tatarenkova, A. Fazekas, V.M. Macias, E. Bier, A. James, Highly efficient Cas9-mediated gene drive for population modification of the malaria vector mosquito Anopheles stephensi. Proceedings of the National Academy of Sciences of the United States of America 112(49):E6736–43 (2015).

33. R. García, J.M. Rodríguez-Peña, C. Bermejo, C. Nombela, J. Arroyo, The high osmotic response and cell wall integrity pathways cooperate to regulate transcriptional responses to zymolyase-induced cell wall stress in *Saccharomyces cerevisiae*. The Journal of Biological Chemistry 284(16):10901–1. (2009).

34. R.D. Gietz, R.H. Schiestl, A.R. Willems, R.A. Woods, Studies on the transformation of intact yeast cells by the LiAc/SS-DNA/PEG procedure. Yeast 11(4):355–60 (1995).

35. C. Grey, F. Baudat, D. de Massy, PRDM9, a driver of the genetic map. PLOS Genetics 14(8):e1007479 (2018).

36. P. Grognet, H. Lalucque, F. Malagnac, P. Silar, Genes that bias Mendelian segregation. PLOS Genetics 10:e1004387 (2014).

37. J. Guan, P.E. Stromhaug, M.D. George, P. Habibzadegah-Tari, A. Bevan, W.A. Dunn Jr, D.J. Klionsky, Cvt18/Gsa12 is required for cytoplasm-to-vacuole transport, pexophagy, and autophagy in Saccharomyces cerevisiae and Pichia pastoris. Molecular Biology of the Cell 12(12):3821–38 (2001).

38. M.F. Hammer, J. Schimenti, L.M. Silver, Evolution of mouse chromosome 17 and the origin of inversions associated with t haplotypes. Proceedings of the National Academy of Sciences of the United States of America 86(9):3261–3265 (1989).

39. T.M. Hammond, D.G. Rehard, H. Xiao, P.K. Shiu, Molecular dissection of Neurospora spore killer meiotic drive elements. Proceedings of the National Academy of Sciences of the United States of America 109(30):12093–8 (2012).

40. C. He, H. Song, T. Yorimitsu, I. Monastyrska, W.L. Yen, J.E. Legakis, D.J. Klionsky, Recruitment of Atg9 to the preautophagosomal structure by Atg11 is essential for selective autophagy in budding yeast. Journal of Cell Biology 175(6):925–935 (2006).

41. B. Herrmann, B. Koschorz, K. Wertz, et al., A Protein Kinase Encoded by the t-Complex Responder Gene Causes Non-Mendelian Inheritance. Nature 402(6758):141–146. (1999).

42. S.M. Hill, S. Hanzén, T. Nyström, Restricted access: spatial sequestration of damaged proteins during stress and aging. EMBO Reports 18(3):377–391 (2017).

43. S.M. Hill, X. Hao, J. Gronvall, S. Spinkings-Nordby, et al., Asymmetric Inheritance of Aggregated Proteins and Age Reset in Yeast Are Regulated by Vac17-Dependent Vacuolar Functions. Cell Reports 16(3):826–38 (2016).

44. C.S. Hoffman, V. Wood, P.A. Fantes, An Ancient Yeast for Young Geneticists: A Primer on the Schizosaccharomyces pombe Model System. Genetics 201(2):403–423 (2015)

45. W. Hu, Z. Jiang, F. Suo, J. Zheng, W. He, L. Du, A large gene family in fission yeast encodes spore killers that subvert Mendel’s law. eLife 6:e26033 (2017).

46. C. Janke, M. Magiera, N. Rathfelder, C. Taxis, S. Reber, H. Maekawa, et al., A versatile toolbox for PCR-based tagging of yeast genes: new fluorescent proteins, more markers and promoter substitution cassettes. Yeast 21:947–962 (2004).

47. C. Jin, SK Kim, SD Willis, KF Cooper, The MAPKKKs Ste11 and Bck1 jointly transduce the high oxidative stress signal through the cell wall integrity MAP kinase pathway. Microb Cell 2(9):329–342 (2015).

48. B. Johnson, D. Snead, J.J. Lee, J.M. McCaffery, J. Shorter, A.D. Gitler, TDP-43 is intrinsically aggregation-prone, and amyotrophic lateral sclerosis-linked mutations accelerate aggregation and increase toxicity. The Journal of Biological Chemistry 284(30):20329–39 (2009).

49. D. Kaganovich, R. Kopito, J. Frydman, Misfolded proteins partition between two distinct quality control compartments. Nature 454(7208):1088–1095 (2008).

50. J.F.R. Kerr, A.H. Wyllie, A.R. Currie, Apoptosis: a basic biological phenomenon with wide-ranging implication in tissue kinetics. British Journal of Cancer 26(4):239–57 (1972).

51. T. Khan, et al., Quantifying Nucleation In Vivo Reveals the Physical Basis of Prion-like Phase Behavior. Molecular Cell 71(1):155–168 (2018).

52. M. Klutstein, A. Fennell, A. Fernández-Álvarez, J.P. Cooper, The telomere bouquet regulates meiotic centromere assembly. Nat Cell Biol. 17(4):458–69(2015).

53. T.A. Kohda, K. Tanaka, M. Konomi, M. Sato, M. Osumi, M. Yamamoto, Fission yeast autophagy induced by nitrogen starvation generates a nitrogen source that drives adaptation processes. Genes to Cells 12(2):155–170 (2007).

54. J. Kremer, D. Mastronarde, and J. McIntosh, Computer visualization of three-dimensional image data using IMOD. Journal of Structural Biology 116(1):71–76 (1996).

55. A. Krogh, B. Larsson, G. von Heijne, EL. Sonnhammer, Predicting transmembrane protein topology with a hidden Markov model: application to complete genomes. Journal of Molecular Biology 305:567–580 (2001).

56. A.N. Kruger, M.A. Brogley, J.L. Huizinga, J.M. Kidd, D.G. de Rooij, Y-C. Hu, J.L. Mueller, A Neofunctionalized X-linked Ampliconic Gene Family Is Essential for Male Fertility and Equal Sex Ratio in Mice. Current Biology 29(21):3699–3706 (2019).

57. D. Kryndushkin, G. Ihrke, T.C. Piermartiri, F. Shewmaker, A yeast model of optineurin proteinopathy reveals a unique aggregation pattern associated with cellular toxicity. Molecular Microbiology 86(6):1531–47 (2012).

58. Z. Kuang, J.D. Boeke, S. Canzar, The dynamic landscape of fission yeast meiosis alternative-splice isoforms. Genome Res. 27(1):145–156 (2016).

59. R. Kumar, P.P. Nawroth, J. Tyedmers, Hitchhiking vesicular transport routes to the vacuole: Amyloid recruitment to the Insoluble Protein Deposit (IPOD). Prion 11(2):71–81 (2017).

60. R. Kumar, P.P. Nawroth, J. Tyedmers, Prion Aggregates Are Recruited to the Insoluble Protein Deposit (IPOD) via Myosin 2-Based Vesicular Transport. PLOS Genetics 12(9): e1006324 (2016)

61. A.M. Larracuente and D.C. Presgraves, The selfish Segregation Distorter gene complex of Drosophila melanogaster. Genetics 192(1):33–53 (2012).

62. X. Li, T. Ohmori, K. Irie, Y. Kimura, Y. Suda, T. Mizuno, K. Irie, Different regulations of ROM2 and LRG1 expression by Ccr4, Pop2, and Dhh1 in the Saccharomyces cerevisiae cell wall integrity pathway. mSphere 1(5):e00250–16 (2016).

63. C.J. Lin, F. Hu, R. Dubruille, J. Vedanayagam, J. Wen, P. Smibert, B. Loppin, E.C. Lai, The hpRNA/RNAi Pathway Is Essential to Resolve Intragenomic Conflict in the Drosophila Male Germline. Developmental Cell 46(3):316–326 (2018).

64. A.K. Lindholm, K.A. Dyer, R.C. Firman, L. Fishman, W. Forstmeier, L. Holman, H. Johannesson, U. Knief, H. Kokko, A.M. Larracuente, A. Manser, C. Montchamp-Moreau, V.G. Petrosyan, A. Pomiankowski, D.C. Presgraves, L.D. Safronova, A. Sutter, R.L. Unckless, R.L. Verspoor, N. Wedell, et al., The Ecology and Evolutionary Dynamics of Meiotic Drive. Trends in Ecology and Evolution 31(4):315–326 (2016).

65. G. Liu, A.N. Coyne, F. Pei, S. Vaughan, M. Chaung, D.C. Zarnescu, J.R. Buchan, Endocytosis regulates TDP-43 toxicity and turnover. Nature communications 8(1):2092 (2017).

66. B. Liu, L. Larsson, A. Caballero, X. Hao, D. Oling, J. Grantham, T. Nystrom, The polarisome is required for segregation and retrograde transport of protein aggregates. Cell 140(2):257–267 (2010).

67. Y. Long, L. Zhao, B. Niu, J. Su, H. Wu, et al., Hybrid male sterility in rice controlled by interaction between divergent alleles of two adjacent genes. Proceedings of the National Academy of Sciences of the United States of America 105(48):18871–6 (2008).

68. J.F. López Hernández, S. E. Zanders, Veni, vidi, vici: the success of *wtf* meiotic drivers in fission yeast. Yeast 35(7):447–453 (2018).

69. A. Lupas, M. Van Dyke, and J. Stock, Predicting Coiled Coils from Protein Sequences. Science 252:1162–1164 (1991).

70. R.S. Marshall, F. McLoughlin, R.D. Vierstra, Autophagic Turnover of Inactive 26S Proteasomes in Yeast Is Directed by the Ubiquitin Receptor Cue5 and the Hsp42 Chaperone. Cell 16(6): 1717–1732 (2016).

71. A. Matsuyama, R. Arai, Y. Yashiroda, et al., ORFeome cloning and global analysis of protein localization in the fission yeast *Schizosaccharomyces pombe*. Nature Biotechnology 24: 841–847 (2006).

72. A.M. Neiman, Sporulation in the Budding Yeast *Saccharomyces cerevisiae*. Genetics 189(3): 737–765 (2011).

73. N.L. Nuckolls, M.A. Bravo Núñez, M.T. Eickbush, J.M. Young, J.J. Lange, J.S. Yu, G.R. Smith, S.L. Jaspersen, H.S. Malik, S.E. Zanders, *wtf* genes are prolific dual poison– antidote meiotic drivers. eLife 6:e26033 (2017).

74. M.J. Ohira, D.G. Hendrickson, R.S. McIsaac, N. Rhind, An estradiol-inducible promoter enables fast, graduated control of gene expression in fission yeast. Yeast 34(8):323–334 (2017).

75. D.S. Ottoz, F. Rudolf, J. Stelling, Inducible, tightly regulated and growth condition-independent transcription factor in Saccharomyces cerevisiae. Nucleic Acids Research 42(17):e130 (2014).

76. F. Pardo-Manuel de Villena, C. Sapienza, Female meiosis drives karyotypic evolution in mammals. Genetics 159(3):1179–1189 (2001).

77. K.E. Pieper, R.L. Unckless, K.A. Dyer, A fast-evolving X-linked duplicate of importin-α2 is overexpressed in sex-ratio drive in *Drosophila neotestacea*. Molecular Ecology 27(24):5165–79 (2018).

78. N. Rhind, Z. Chen, M. Yassour, et al., Comparative functional genomics of the fission yeasts. Science 332(6032):930–6 (2011).

79. N.A. Rhoades, A.M. Harvey, D.A. Samarajeewa, J. Svedberg, A. Yusifov, A. Abusharekh, et al., Identification of *rfk-1*, a meiotic driver undergoing RNA editing in *Neurospora*. Genetics 212(1):93–110 (2019).

80. R. Riek. S.J. Saupe, The HET-S/s prion motif in the control of programmed cell death. Cold Spring Harb. Perspect. Biol. 8(9):a023515 (2016).

81. A.D. Roeder, J.M. Shaw, Vacuole partitioning during meiotic division in yeast. Genetics 144(2):445–58 (1996).

82. O. Ronneberger, P. Fischer, T. Brox, U-Net: Convolutional Networks for Biomedical Image Segmentation. In: N. Navab, J. Hornegger, W. Wells, A. Frangi (eds) Medical Image Computing and Computer-Assisted Intervention. Lecture Notes in Computer Science 9351, Springer, Cham (2015).

83. S. Rothe, A. Prakash, J. Tyedmers, The Insoluble Protein Deposit (IPOD) in Yeast. Frontiers in Molecular Neuroscience 11:237 (2018).

84. L. Ruan, C. Zhou, E. Jin, A. Kucharavy, Y. Zhang, Z. Wen, L. Florens, R. Li, Cytosolic proteostasis through importing of misfolded proteins into mitochondria. Nature 543(7645):443–446 (2017).

85. L. Sandler, E. Novitski, Meiotic Drive as an Evolutionary Force. The American Naturalist 91:105–110 (1957).

86. J. Schimenti, Segregation distortion of mouse *t* haplotypes the molecular basis emerges. Trends in Genetics 16(6):240–243 (2000).

87. R.B. Sekar, A. Periasamy, Fluorescence resonance energy transfer (FRET) microscopy imaging of live cell protein localizations. Journal of Cell Biology 160(5):629–633 (2003).

88. R. Shen, L. Wang, X. Liu, et al., Genomic structural variation-mediated allelic suppression causes hybrid male sterility in rice. Nature Communications 8:1310 (2017).

89. R.S. Sikorski, P. Hieter, A system of shuttle vectors and yeast host strains designed for efficient manipulation of DNA in *Saccharomyces cerevisiae*. Genetics 122(1):19–27 (1989).

90. G. Singh, AJ. Klar, The 2.1-kb inverted repeat DNA sequences flank the mat2,3 silent region in two species of *Schizosaccharomyces* and are involved in epigenetic silencing in *Schizosaccharomyces pombe*. Genetics 162:591–602 (2002).

91. B.D. Slaughter, J.R. Unruh, A. Das, S.E. Smith, B. Rubinstein, R. Li, Non-uniform membrane diffusion enables steady-state cell polarization via vesicular trafficking. Nature Communications 4:1380 (2013).

92. G.R. Smith, Genetic analysis of meiotic recombination in *Schizosaccharomyces pombe*. Methods Mol Biol. 557:65–76 (2009).

93. J.H. Soper, S. Roy, A. Stieber, E. Lee, R.B. Wilson, J.Q. Trojanowski, C.G. Burd, V.M. Lee, α-Synuclein–induced Aggregation of Cytoplasmic Vesicles in *Saccharomyces cerevisiae*. Molecular Biology of the Cell 19(3):1093–1103 (2008).

94. M.S. Stewart, S.A. Krause, J. McGhie, J.V. Gray, Mpt5p, a stress tolerance- and lifespan-promoting PUF protein in *Saccharomyces cerevisiae*, acts upstream of the cell wall integrity pathway. Eukaryotic Cell 6(2):262–70. (2007).

95. Y. Suda, H. Nakanishi, E. Mathieson, A. Neiman, Alternative Modes of Organellar Segregation during Sporulation in *Saccharomyces cerevisiae*. Eukaryotic Cell 6(11):2009–2017 (2007).

96. L.L. Sun, M. Li, F. Suo, et al., Global Analysis of Fission Yeast Mating Genes Reveals New Autophagy Factors. PLOS Genetics 9(8):e1003715 (2013).

97. K. Suzuki, Y. Ohsumi, Current knowledge of the pre-autophagosomal structure (PAS). FEBS Letters 584(7):1280–1286 (2010).

98. J.P. Taylor, F. Tanaka, J. Robitschek, C.M. Sandoval, A. Taye, S. Markovic-Plese, K.H. Fischbeck, Aggresomes protect cells by enhancing the degradation of toxic polyglutamine-containing protein. Human Molecular Genetics 12(7):749–757 (2003).

99. K. Tomita, J.P. Cooper, The telomere bouquet controls the meiotic spindle. Cell 130(1):113–26 (2007).

100. J. Tyedmers, S. Treusch, J. Dong, J.M. McCaffery, B. Bevis, S. Lindquist, Prion induction involves an ancient system for the sequestration of aggregated proteins and heritable changes in prion fragmentation. Proceedings of the National Academy of Sciences of the United States of America 107(19):8633–8638 (2010).

101. A.A. Vogan, S.L. Ament-Velásquez, A. Granger-Farbos, J. Svedberg, E. Bastiaans, A.J. Debets, V. Coustou, H. Yvanne, C. Clavé, S.J. Saupe, H. Johannesson, Combinations of Spok genes create multiple meiotic drivers in Podospora. eLife 8:e46454 (2019).

102. S. Willingham, T.F. Outeiro, M.J. DeVit, S.L. Lindquist, P.J. Muchowski, Yeast genes that enhance the toxicity of a mutant huntingtin fragment or alpha-synuclein. Science 302(5651):1769–72 (2003).

103. Y. Xie, J. Tang, X. Xie, X. Li, J. Huang, et al., An asymmetric allelicinteraction drives allele transmission bias in interspecific rice hybrids. Nature Communications 10(1):2501 (2019).

104. X. Yu, Z. Zhao, X. Zheng, J. Zhou, et al., A selfish genetic element confers non-Mendelian inheritance in rice. Science 360:1130–1132 (2018).

105. S.E. Zanders, M.T. Eickbush, J.S. Yu, J. Kang, K. Fowler, G.R. Smith, H.S. Malik, Genome rearrangements and pervasive meiotic drive cause hybrid infertility in fission yeast. eLife 3:e02630 (2014).

106. D. Zhang, A. Vjestica, S. Oliferenko, Plasma membrane tethering of the cortical ER necessitates its finely reticulated architecture. Current Biology 22(21): 2048–52 (2012).

107. H. Zhang, S. Elbaum-Garfinkle, E. Langdon, N. Taylor, P. Occhipinti, A. Bridges, C. P. Brangwynne, and A.S. Gladfelter, RNA controls PolyQ protein phase transitions. Molecular Cell 60(2):220–230 (2015).

108. D. Zhao, et al., Atg20- and Atg24-family proteins promote organelle autophagy in fission yeast. Journal of Cell Science 129:4289–4304 (2016).

109. C. Zhou, B.D. Slaughter, J.R. Unruh, F. Guo, Yu Z, K. Mickey, A. Narkar, R.T. Ross, M. McClain, R. Li R, Organelle-based aggregation and retention of damaged proteins in asymmetrically dividing cells. Cell 159(3):530–42 (2014).

